# Unveiling the Promise of Peptide Nucleic Acids as Functional Linkers for Riboglow RNA Imaging Platform

**DOI:** 10.1101/2024.10.03.616516

**Authors:** Aleksandra J. Wierzba, Erin M. Richards, Shelby R. Lennon, Robert T. Batey, Amy E. Palmer

**Author notes:** To whom correspondence should be addressed. R.T.B.; A.E.P.

## Abstract

Linkers in chemical biology provide more than just connectivity between molecules; their intrinsic properties can be harnessed to enhance the stability and functionality of chemical probes. In this study, we explored the incorporation of a peptide nucleic acid (PNA)-based linker into RNA-targeting probes to improve their affinity and specificity. By integrating a PNA linker into a small molecule probe of Riboglow platform, we enabled dual binding events: cobalamin (Cbl)-RNA structure-based recognition and sequence-specific PNA-RNA interaction. We show that incorporating a six-nucleotide PNA sequence complementary to the region of wild type RNA aptamer (*env*8) results in a 30-fold improvement in binding affinity compared to the probe with nonfunctional PEG linker. Even greater improvements are observed when the PNA probe was tested against truncated versions of the RNA aptamer, with affinity increasing by up to 280-fold. Additionally, the PNA linker is able to rescue Cbl-RNA interaction even when the cobalamin binding pocket is compromised. We demonstrated that PNA probes effectively bind RNA both *in vitro* and in live cells, enhancing visualization of RNA in stress granules and U-bodies at low concentrations. The modular nature of the Riboglow platform allows for flexible modifications of the PNA linker, fluorophore and RNA tag, while maintaining high specificity and affinity. This work establishes a new approach for enhancing RNA imaging platforms through the use of PNA linkers, highlighting the potential of combining short oligonucleotides with small molecules to improve the affinity and specificity of RNA-targeting probes. Furthermore, this dual-binding approach presents a promising strategy for driving advancements in RNA-targeted drug development.

**Table of Contents graphic:** 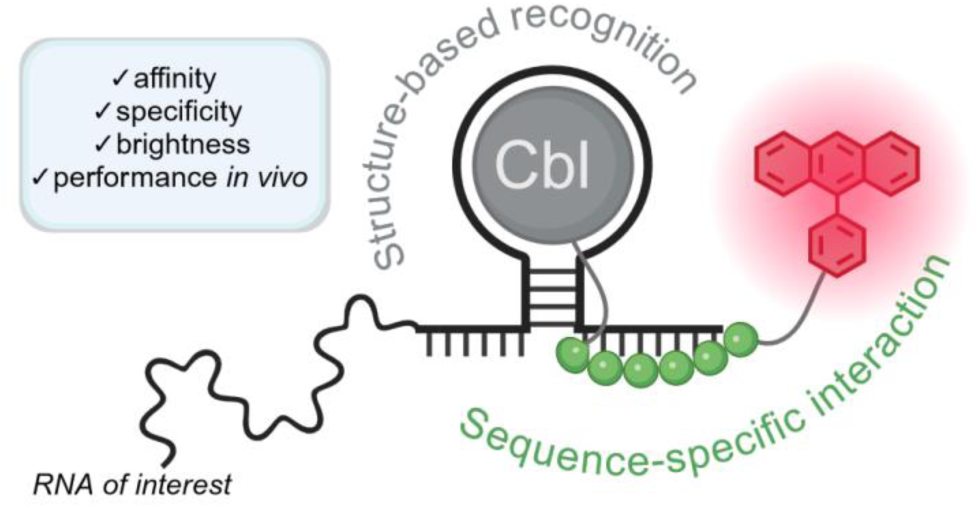

## INTRODUCTION

The ability to visualize macromolecules in live cells has driven many fundamental discoveries in biology over the last few decades. Tagging proteins with fluorophores has revolutionized our understanding of the behavior of individual proteins in the cellular context, in large part due to a diverse and easy-to-use set of tools that continues to expand to meet new experimental challenges.^1^ Conversely, while the roles of RNA in human biology are myriad and diverse, the ability to visualize RNA in live cells is much more limited. The most widely employed RNA imaging tool to date is the MS2-MCP system which uses fluorescently-tagged proteins to track individual RNA molecules in live cells.^2,3^ However, this system has a set of well-established technical limitations that have inspired the development of additional live-cell RNA imaging tools.^4–6^ Over the past decade, much effort has focused on the development of tools that employ a small-molecule binding aptamer as a tag that is incorporated into the RNA of interest.^7,8^ While these next generation RNA imaging tools are beginning to make an impact in addressing important biological questions, their lack of widespread adoption indicates a need for improvements in their capabilities and performance.

One class of RNA probes relies on binding of a conjugated fluorophore-quencher pair to an RNA aptamer, where the binding event separates the fluorophore from the quencher, giving rise to fluorescence turn-on. For example, a conjugate between tetramethylrhodamine (TMR) and a dinitroaniline quencher was created by connecting the pair with a small flexible polyethylene glycol (PEG) linker. When TMR binds to an *in vitro* selected RNA aptamer,^9^ the quencher is separated from the fluorophore, causing an increase in fluorescence.^10^ Fluorophore-binding aptamers suffer from the limitation that a new aptamer needs to be selected for every new fluorophore. To overcome this limitation, some RNA probes are designed so that the quencher binds the RNA tag.^11–13^ For example, Riboglow is an RNA imaging tool in which the RNA aptamer binds to the quencher, enabling the use of a wide array of fluorescent dyes.^13^ In this case, the quencher is cobalamin (Cbl) that binds a biological RNA aptamer derived from a bacterial mRNA riboswitch that represses translation of the message in response to cobalamin binding. In each case, the fluorophore and quencher are connected by a PEG linker, which was chosen for its flexibility and ease of installation into the probe.

Linkers are an indispensable tool in chemical biology, providing not only structural connectivity but also functional versatility in the design and synthesis of bioconjugates.^14–16^ PEG is widely used in chemical probes due to its biological inertness, hydrophilicity and flexibility, all of which make it an attractive choice as a linker between two functional parts of a chemical probe, where it is presumed to be innocuous. We speculated, however, that using PEG linkers in RNA imaging probes might represent a missed opportunity to add chemical functionality to the linker. A functionalized linker could interact with the target or influence the probe’s behavior. In our previous work on Riboglow, we tested a limited set of different linkers, focusing primarily on their length as a factor affecting probe properties, particularly the optimal fluorophore-quencher separation.^13^ We demonstrated that the optimal linker length resulted in effective quenching/dequenching platform properties. Moreover, a more rigid linker consisting of four glycine units (4xGly linker), compared to 5xPEG, improved the probe’s turn-on efficiency. Building on this, we became particularly interested in incorporating a short, antisense oligonucleotide (ASO) that could base pair with a target RNA. This is a new approach to probe design that provides additional binding capabilities that could impact the functionality of RNA-targeted probes.

Peptide nucleic acids (PNAs) hold great promise as a nucleic acid mimic.^17–19^ These nucleic acid analogs have a backbone composed of repeating *N*-(2-aminoethyl)-glycine units linked by peptide bonds rather than sugar-phosphate groups. As a consequence, they exhibit enhanced stability against enzymatic degradation and form stronger duplexes with complementary DNA or RNA than either DNA or RNA homoduplexes due to the lack of charge repulsion between the strands.^19^ PNAs have been shown to function as antisense agents, gene-editing tools, and crosslinkers,^17^ as well as fluorogenic probes for nucleic acid sensing when tethered to a fluorescent dye (molecular beacons).^20^ Despite these advantages, there are notable challenges that deter scientists from using PNAs, such as poor cell penetration, limited solubility, and a tendency to form aggregates.^21^ However, these issues can be mitigated by coupling PNAs with appropriate molecules.^22,23^ Previous work has shown that PNAs can be coupled to cobalamin to facilitate their uptake through the bacterial cobalamin import system and act as an ASO to repress gene expression.^24–28^ Additionally, conjugation of PNA to hydrophilic cobalamin significantly improves the solubility of the synthetic oligonucleotide. Together, these observations suggest that PNAs may serve as ideal candidates as “smart” linkers in contact quenching RNA probes.

In this work, we explore the impact of functional linkers on an RNA imaging platform by substituting the neutral PEG linker in Riboglow with PNA linkers designed to interact with the target RNA via base pairing. We designed a six-nucleotide PNA sequence complementary to a single-stranded region within the aptamer that does not engage in cobalamin binding, which is crucial for maintaining cobalamin recognition. Using chemical probing, we demonstrate the direct engagement of the PNA linker with the target RNA sequence along with effective cobalamin binding. Base pairing of the PNA linker with the RNA aptamer combined with the cobalamin-RNA binding significantly increases the probe affinity. *In vitro* fluorescence turn-on assays revealed the K_D_ decreases 30-fold to 0.11 ± 0.02 nM for the full-length wild type *env*8 cobalamin riboswitch aptamer when compared to a probe with a 5xPEG linker. We show that scrambling the PNA sequence in the probe abolishes its interaction with the RNA, but the sequence specific interaction can be rescued by making a compensatory change in the RNA. An even more beneficial impact of PNA was observed for truncated versions of the RNA aptamer, in which introduction of a PNA binding site on either the 3’ or 5’ end led to an increase in affinity by 60 to 80-fold (for two different sequences at the 3’ end) or 220 to 280-fold (for two different sequences at the 5’ end). Further, we show that the PNA linker can drive binding between the probe and the RNA, even when the binding pocket for cobalamin is significantly compromised. Cell studies show the superiority of the PNA probes compared to those with nonfunctional linkers. The PNA probes show stronger enrichment in RNA-protein granules (stress granules or U-bodies) in live cells expressing aptamer-tagged RNA-of-interest. In summary, we establish a new pathway for improvement of RNA imaging tools as well as the potential for using short oligonucleotides conjugated to small molecules to improve affinity and specificity of chemical probes targeting RNA.

## RESULTS

### Synthesis of cobalamin-PNA-fluorophore conjugates

We set out to install chemical functionality in the linker of cobalamin-fluorophore probes to improve the affinity and specificity of the Riboglow RNA imaging platform. To accomplish this, we replaced the flexible polyethylene glycol linker in the original Riboglow probes^13^ with a PNA linker. Based on the crystal structure and secondary structural elements, we designed the PNA sequence to target the single-stranded J1/13 region in the wild type cobalamin *env*8 riboswitch (Supporting Figure S1).^29^ The targeted site within the riboswitch does not engage in cobalamin binding; therefore, it was envisioned as a suitable landing pad for the complementary linker. We chose a PNA linker rather than one of the other nucleic acid analogs because cobalamin-PNA conjugates have been previously synthesized and shown to function in bacterial cells as an antisense oligonucleotide.^24–28^ The mode of action of Cbl-PNA probe and the RNA tag, involving both cobalamin binding by the riboswitch and the engagement of sequence-specific interaction between the linker and the complementary RNA region, is shown in Figure 1A.

**Figure 1.**
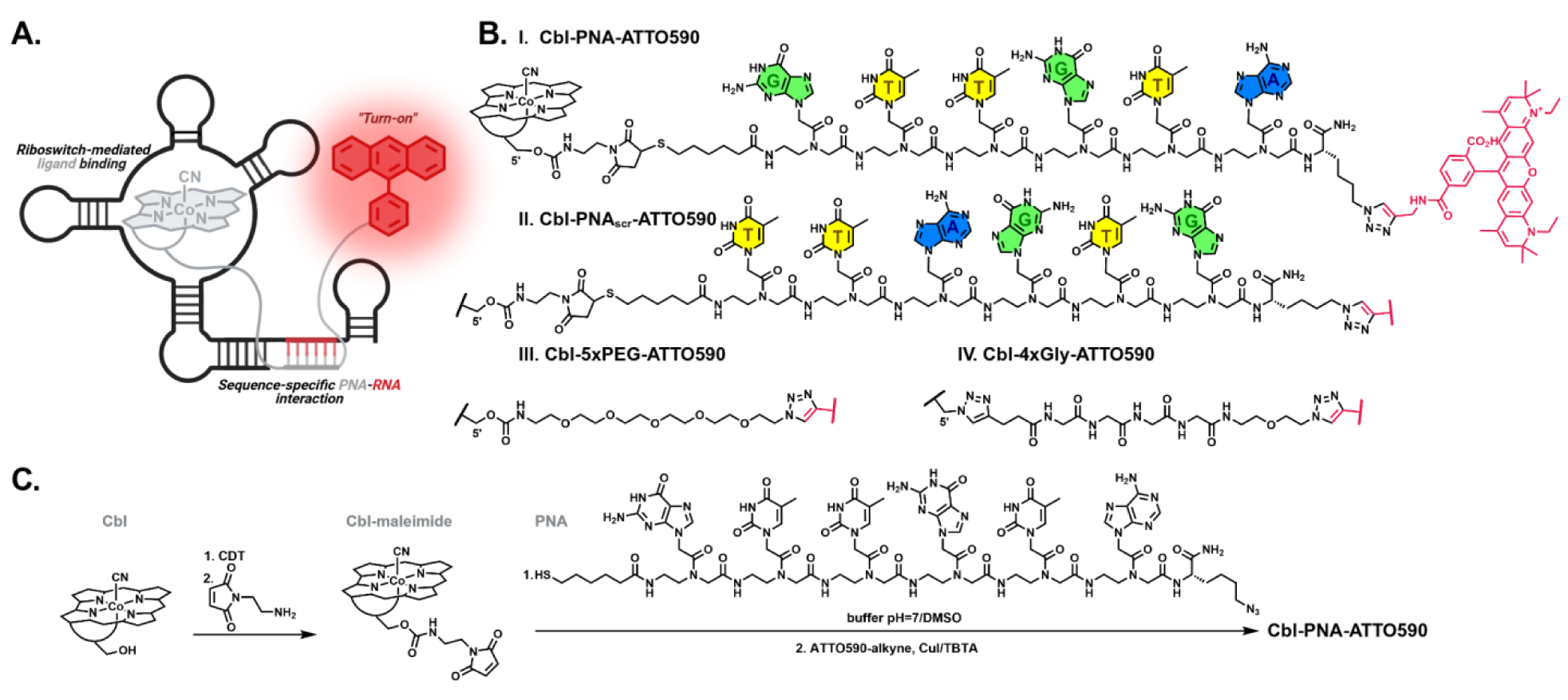
**A**. General concept of the mode of action of cobalamin-based probes comprising a PNA linker. **B**. Chemical structure of the cobalamin-based fluorescent probes used in the study (PNA = GTTGTA sequence complementary to J1/13 region in wild type *env*8, PNA_scr_ = PNA linker with scrambled sequence TTAGTG, 5xPEG = linker of 5 polyethylene glycol units, 4xGly = linker of 4 glycine units). **C**. Synthetic pathway leading to the Cbl-PNA-ATTO590 probe.

Cbl-PNA-dye probes were synthesized by preparing a specifically designed cobalamin derivative, followed by the subsequent attachment of a suitably tailored PNA oligomer to Cbl and the addition of a fluorescent dye using previously established methods (Figure 1B and C).^13,24^ PNA oligomers were manually synthesized using solid-phase synthesis (Supporting Information 3.2). The strand was obtained by coupling six monomers together through Fmoc chemistry and tagging the N-terminus and C-terminus with two distinct functionalities: a sulfhydryl group and an azide group, respectively. To obtain the thiol-reactive cobalamin, its primary hydroxyl group was modified via carbamate chemistry, leading to the introduction of a maleimide moiety (Cbl-maleimide, Figure 1C). This derivative was then successfully coupled with the thiol end of the PNA strand, yielding the thiosuccinimide product, the Cbl-PNA conjugate (Supporting Figure S2). The target Cbl-PNA-dye probe was obtained via copper(I)-catalyzed azide-alkyne cycloaddition between the alkyne-modified dye and the azide end of the PNA, resulting in a covalently linked hybrid (Figure 1B and C, Supporting Information 3.8).

The Cbl-PNA-ATTO590 probe possesses a sequence targeting the complementary single stranded region of the wild type *env*8 cobalamin riboswitch (*env*8-FL-3’*anti*PNA, Supporting Figure S4). To assess the specificity of PNA to base pair with the targeted RNA region, the Cbl-PNA-ATTO590 probe was tested against a “scrambled” variant, Cbl-PNA_scr_-ATTO590 (Figure 1B, Supporting Figure S3), which is equipped with the same set of nucleobases, but arranged in a different order to abrogate base-pairing with the RNA aptamer. Additionally, we compared the PNA probe to the previously published probes with neutral chemical linkers, namely Cbl-5xPEG-ATTO590 and Cbl-4xGly-ATTO590 (Figure 1B, Supporting Figure S3).^13^

### The PNA probe productively base pairs with the riboswitch

To visualize RNA-PNA pairing, we used a chemical footprinting approach called “Selective 2’-hydroxyl acylation analyzed by primer extension” (SHAPE).^30,31^ This technique exploits small electrophilic reagents (here *N*-methylisatoic anhydride, NMIA) that react with 2’-hydroxyl groups in flexible regions of the RNA to interrogate its structure at single-nucleotide resolution. Thus, when a given nucleotide is involved in secondary structural elements (i.e., base pairing to form a helix), the 2’-hydroxyl has strongly reduced reactivity towards NMIA. We visualized chemical reactivity of each 2’-hydroxyl group in *env*8 RNA using radiolabeled primer extension and electrophoresis of the resulting cDNAs on a sequencing gel.

SHAPE chemical probing revealed Cbl and PNA interactions between the Cbl-conjugates (Supporting Figure S2) and *env*8-FL-3’*anti*PNA. The Cbl-specific signature of protection within the L5, J6/3, and L13 RNA regions of the *env*8 riboswitch was the same for Cbl-PNA, Cbl- PNA_scr_, and Cbl alone (Figure 2A, left gel, Supporting Figure S5). This reveals that conjugation of the PNA to Cbl does not impact the ability of Cbl to interact productively with the RNA. In contrast, the J1/13 region revealed ligand-dependent protection between Cbl-PNA and wild type *env*8-FL-3’*anti*PNA (Figure 2A, left gel) but not with the Cbl-PNA_scr_. This suggests that the PNA interacts with the RNA in a sequence-specific fashion.

**Figure 2.**
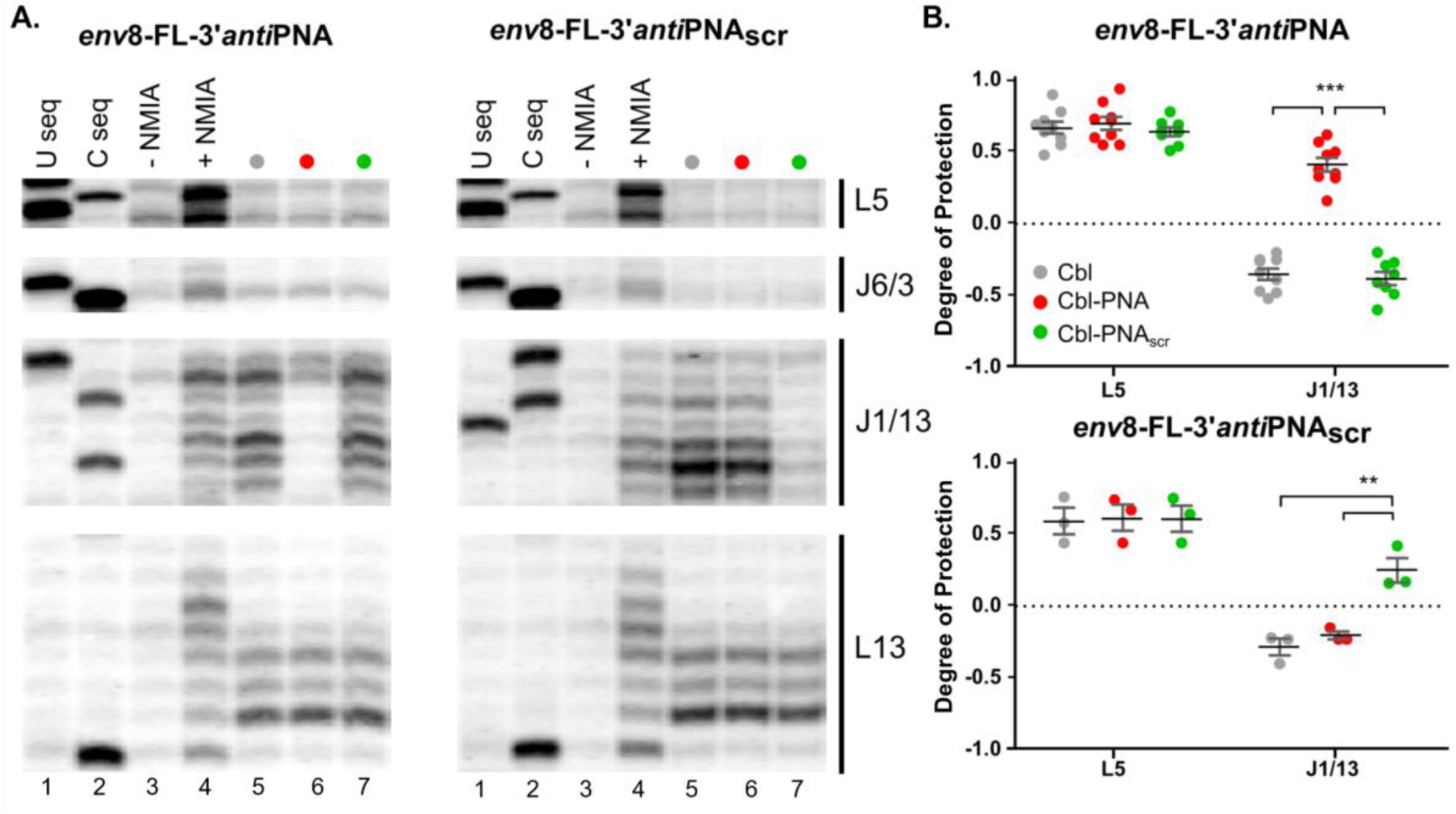
Cbl-PNA conjugates bind to wild type *env*8 in a sequence specific manner. **A.** Representative regions of wild type *env*8-FL-3’*anti*PNA (left) and *env*8-FL-*anti*PNA_scr_ (right) SHAPE gels in the presence of Cbl (gray), Cbl-PNA (red), and Cbl-PNA_scr_ (green) ligands. Lanes are loaded according to the key above. Cbl binding of all three ligands is seen by ligand dependent changes in regions L5, J6/3, and L13 (lanes 5, 6, & 7 compared to lane 4). Sequence specific annealing of the PNA is seen by ligand dependent protections in J1/13 (lane 6 in A and lane 7 in B compared to lane 4). **B.** Quantifications of L5 and J1/13 regions of *env*8-FL-3’*anti*PNA (top) and *env*8-FL-*anti*PNAscr (bottom) SHAPE gels with the same three ligands as A. Degree of protection is in comparison to the +NMIA condition. Unpaired t-tests, *** p ≤ 0.001, ** p < 0.01. Not significant p-values (>0.5) are not marked. n= 3-9. Error bars represent SEM.

To demonstrate conclusively that the PNA-RNA interaction is sequence specific, the same chemical probing experiments were performed using an RNA whose J1/13 sequence was altered to be complementary to the PNA_scr_ sequence. These data revealed that the PNA-mediated base pairing interaction is rescued when the sequence of J1/13 was altered to match that of Cbl-PNA_scr_ (*env*8-FL-3’*anti*PNA_scr_, Figure 2A, right gel, Supporting Figure S6). Hence both Cbl-PNA and Cbl-PNA_scr_ conjugates engaged in base pairing only when a complementary sequence was present within the RNA structure. Quantification of reactivity in the J1/13 regions of *env*8-FL-3’*anti*PNA and *env*8-FL-*anti*PNA_scr_ indicated significant protections of this fragment for both complementary pairs (Figure 2B, top and bottom, respectively) compared to the ligands lacking the sequence-specific fragment. Together, these data reveal specific RNA-PNA hybridization in the context of a small molecule-PNA conjugate. Further, we show the sequence of the RNA aptamer can be mutated to create a PNA binding site, indicating that PNA sequences are not limited to those that pair with wild type RNA but could be expanded to a large number of potential PNA sequences that pair with a complementary region of RNA.

### Quantitative binding analysis reveals that the PNA linker enhances binding affinity of the probe

Since the PNA creates additional interactions with the RNA through base pairing, we predicted that the PNA linker would increase the affinity of Cbl-PNA probes compared to probes in which the linker does not interact with the RNA. To test this, we performed a series of titrations that leverage probe turn-on upon binding due to separation of the fluorophore from its quencher (Figure 3). The affinity was determined by titrating an increasing amount of the RNA into a solution containing a constant amount of Cbl probe and the dissociation constant (K_D_) was determined by fitting the fluorescence data to a quadratic binding equation.^32^

**Figure 3.**
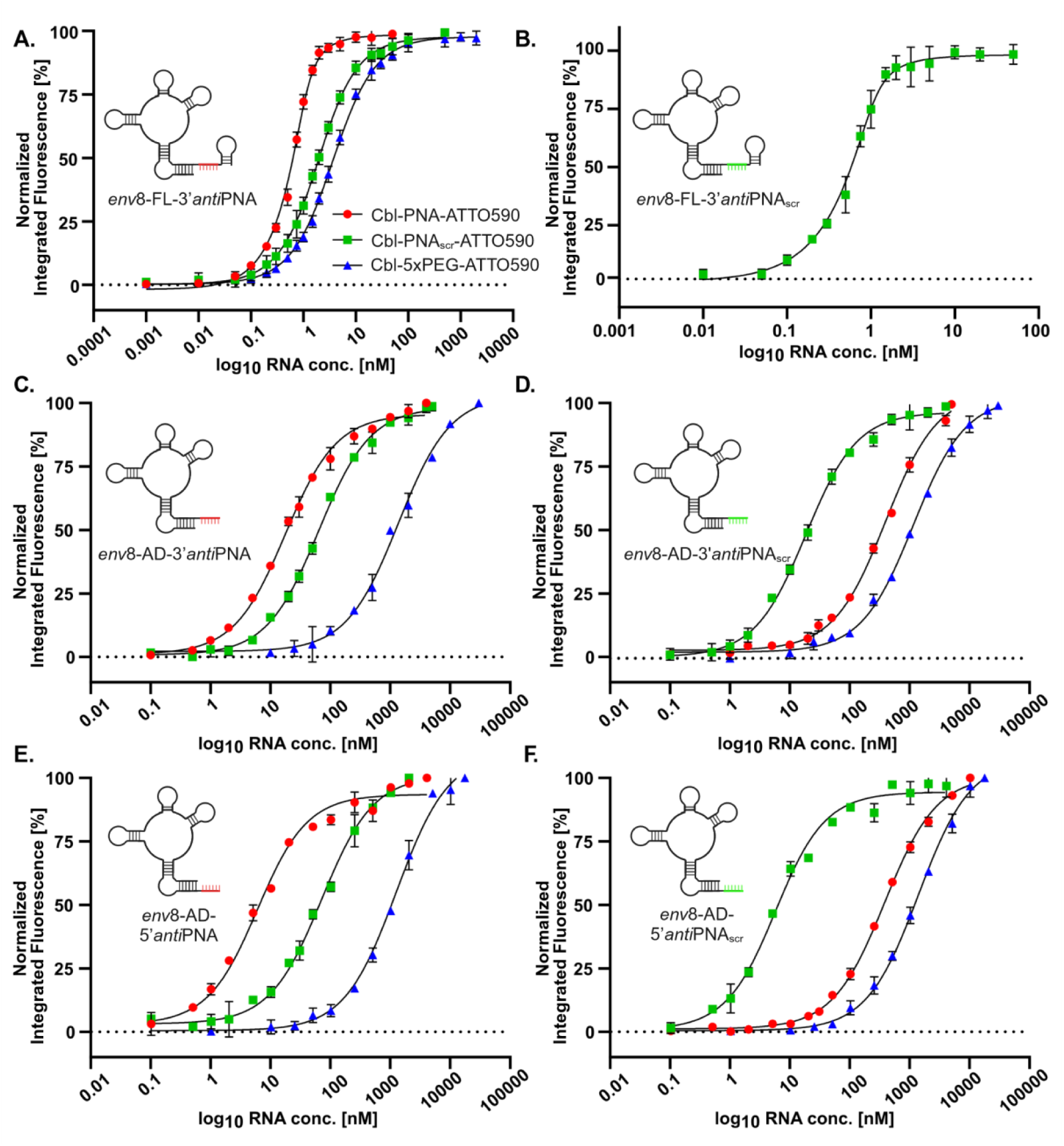
Fluorescence turn-on studies between *env*8 variants and cobalamin-based probes demonstrate the advantage of the PNA linker over non-functional alternatives. **A.** Titration of Cbl probes with wild type *env*8 (c_probe_ = 1 nM, red dot = Cbl-PNA-ATTO590, green square = Cbl-PNA_scr_-ATTO590, blue triangle = Cbl-5xPEG-ATTO590). **B**. Titration of Cbl-PNA_scr_-ATTO590 probe with wild type *env*8 equipped with a modified J1/13 fragment complementary to the PNA_scr_ linker. **C** and **D.** Titration of Cbl probes with truncated *env*8 variants, each containing either the *anti*PNA or *anti*PNA_scr_ fragment installed at the 3’ end. **E** and **F**. Titration of Cbl probes with truncated *env*8 variants, each containing either the *anti*PNA or *anti*PNA_scr_ fragment installed at the 5’ end. n=4-8. Error bars represent SEM.

Binding analysis demonstrated that PNA significantly increased the affinity of Cbl probes for the RNA riboswitch. Cbl-PNA-ATTO590 exhibited tighter binding with wild type *env*8-FL-3’*anti*PNA, with a K_D_ of 0.11 ± 0.02 nM, in comparison to Cbl-PNA_scr_-ATTO590 (K_D_ = 1.4 ± 0.4 nM) and Cbl-5xPEG-ATTO590 (K_D_ = 3.2 ± 0.2 nM) (Figure 3A, Table 1). The Cbl-PNA_scr_-ATTO590 probe bound slightly tighter compared to Cbl-5xPEG-ATTO590 which could be attributed to nonspecific interactions between the RNA and noncomplementary PNA_scr_ sequences. Mutation of *env*8 J1/13 to make it complementary to the Cbl-PNA_scr_-ATTO590 probe (*env8*-FL-3’*anti*PNA_scr_) resulted in comparably tighter binding between the two with a K_D_ of 0.12 ± 0.07 nM (Figure 3B, Table 1), further demonstrating that complementary sequence in the linker leads to significant enhancement of the probe affinity.

**Table 1.** Summary of all K_D_ values [nM] obtained from fluorescence turn-on studies between full length and truncated *env*8 variants and cobalamin-based probes. n=4-8. Error bars represent SEM. nd-value not determined.

| Probe\RNA | *env8*-FL-3'*anti*PNA | *env8*-FL-3'*anti*PNA<sub>scr</sub> | *env8*-AD-3'*anti*PNA | *env8*-AD-3'*anti*PNA<sub>scr</sub> | *env8*-AD-5'*anti*PNA | *env8*-AD-5'*anti*PNA<sub>scr</sub> |
| --- | --- | --- | --- | --- | --- | --- |
| Cbl-PNA-ATTO590 | 0.11 ± 0.02 | nd | 17.2 ± 1.2 | 370 ± 21 | 5.4 ± 0.8 | 373 ± 27 |
| Cbl-PNA<sub>scr</sub>-ATTO590 | 1.4 ± 0.4 | 0.12 ± 0.07 | 65 ± 6 | 17.9 ± 1.4 | 67 ± 9 | 5.0 ± 0.8 |
| Cbl-5xPEG-ATTO590 | 3.2 ± 0.2 | nd | 1400 ± 200 | 1100 ± 100 | 1200 ± 100 | 1400 ± 100 |

### Truncated aptamer domain Cbl riboswitches display enhanced binding by PNA linker probes

The RNA imaging field is in pursuit of relatively small RNA tags to minimally perturb the behavior of the RNA of interest. We previously tested a truncated *env*8 containing only the aptamer domain (*env*8-AD)^13^ for binding to Cbl-5xPEG-ATTO590 probe and found it had significantly reduced affinity (K_D_ = 1.3 ± 0.6 µM) compared to a full length aptamer (K_D_ = 3.2 ± 0.2 nM). Here we tested whether addition of an *anti*PNA binding site could enhance the binding affinity of the truncated *env*8. In the case of the aptamer domain (AD), which lacks the P13 element, the PNA-pairing sequence could be installed on either the 5’- or 3’-side of the aptamer providing more opportunities to optimize the enhancement in affinity between the RNA and probe (Figure 3C-F).

Addition of the *anti*PNA sequence to the 3’ end of truncated *env*8 (*env*8-AD-3’*anti*PNA) led to an 80-fold decrease in the K_D_ value for the PNA probe compared to the 5xPEG probe (Figure 3C, Table 1). The probe with the scrambled PNA sequence showed weaker binding to the noncomplementary aptamer, which was rescued when the complementary sequence was added to the 3’ end (*env*8-AD-3’*anti*PNA_scr_) (Figure 3C, D). A similar trend was observed when the PNA sequence was added to the 5’ end of the truncated *env*8 (*env*8-AD-5’*anti*PNA), with an even greater decrease in the K_D_ value, approximately 280-fold, compared to the 5xPEG probe (Figure 3E, F). As in our previous work,^13^ the probe with the 5xPEG linker exhibited a K_D_ in the micromolar range, whereas probes with complementary PNA sequences showed K_D_ values in the low nanomolar range (Figure 3C-F, Table 1). Inserting the *anti*PNA complementary sequence at the 5’ end resulted in higher affinity (K_D_ ∼ 5 nM) compared to the 3’ end (K_D_ ∼ 17 nM) (Table 1), indicating that binding can be further optimized by altering the placement of the PNA landing pad within the RNA aptamer.

### The PNA linker can drive the binding of cobalamin to an impaired riboswitch

Given that incorporation of an *anti*PNA landing pad into an RNA aptamer could decrease the K_D_ up to 280- fold, we evaluated whether complementary PNA interactions could be the primary driver of binding between a probe and RNA rather than a structure-based recognition by the cobalamin. For this purpose we synthesized an *env*8 variant with four mutations in the Cbl binding pocket that debilitate binding to Cbl but retain the complementary sequence to the PNA (*env*8_mut_-FL- 3’*anti*PNA, Figure 4A). To evaluate Cbl-PNA conjugates binding to the mutated RNA, we used SHAPE chemical probing (Figure 4B, Supporting Figure S7). Ligand-induced protections in the L5 and J6/3 regions of *env*8 are indicative of Cbl binding, as seen in Figure 2. Quantification of the degree of protection in these regions revealed that as the ligand concentration was lowered from 30 µM to 0.3 µM, the Cbl-PNA conjugate showed significantly greater protection compared to Cbl or Cbl-PNA_scr_. This indicates that the PNA linker facilitates Cbl binding to the debilitated aptamer and demonstrates that the PNA can indeed drive binding (Figure 4B-C). As observed for *env*8_mut_-FL-3’*anti*PNA, the J1/13 region showed significant protection upon interaction with Cbl- PNA, but not the other two non-complementary conjugates, confirming specific base pairing between PNA strand in the probe and its complementary sequence in the aptamer.

**Figure 4.**
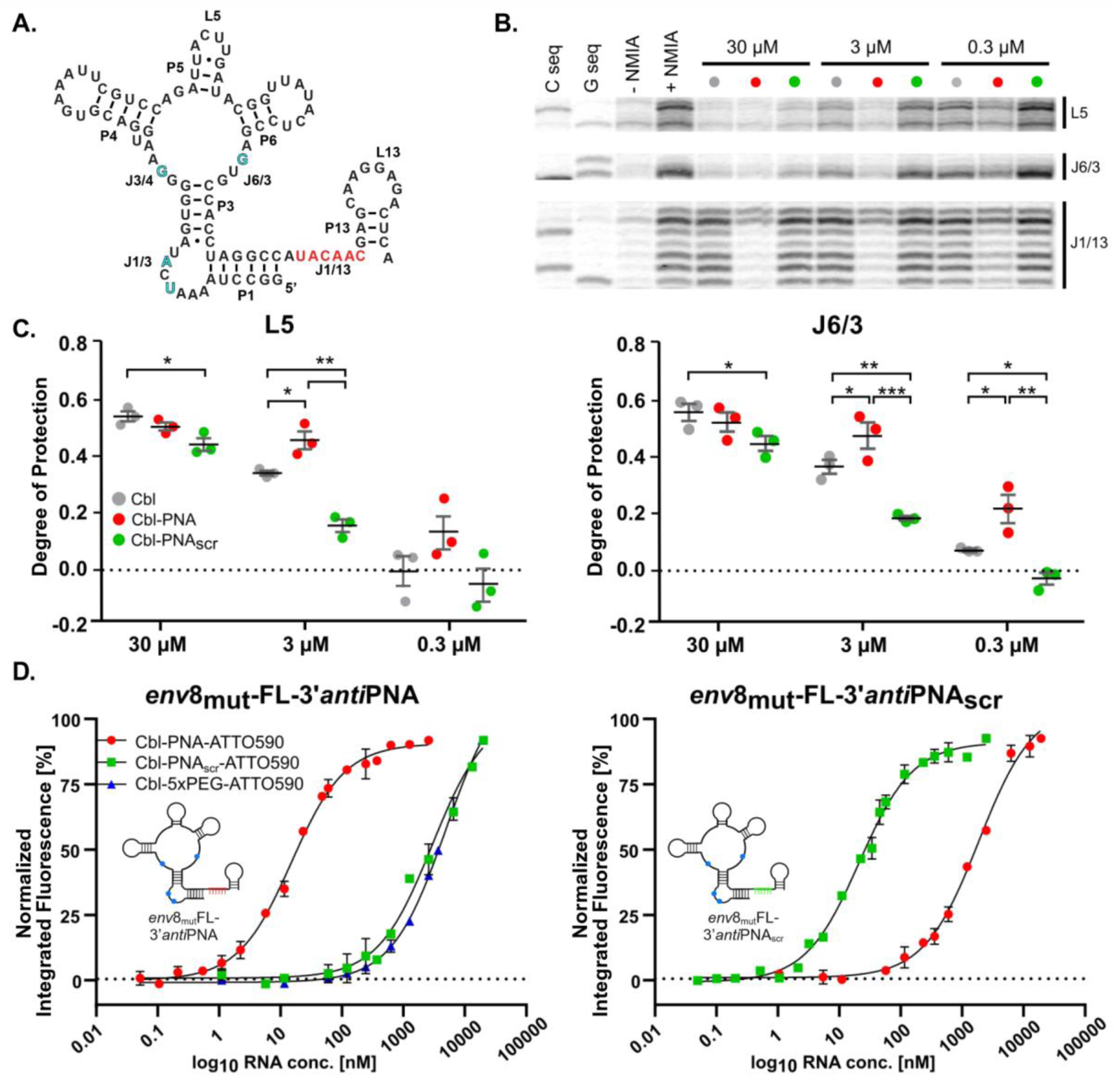
PNA can be used to enhance affinity between a weak RNA-Cbl binder and confer specificity. **A**. 2D representation of mutated variant of wild type *env*8 (mutation sites marked in blue). **B**. Representative SHAPE gel regions of *env*8_mut_-FL- 3’*anti*PNA in the presence of three concentrations of Cbl (grey), Cbl-PNA (red), and Cbl-PNA_scr_ (green) ligands. Cbl binding of all three ligands is seen by ligand dependent protections in regions L5 and J6/3. Sequence specific annealing of the PNA is seen by ligand dependent protections in J1/13. Lanes are loaded according to the key. **C**. Quantifications of L5 and J6/3 SHAPE gel regions with the same ligands as B. Ligands are represented by colored dots according to the key. Degree of protection is in comparison to the +NMIA condition. Unpaired t-tests, ** p < 0.01, * p < 0.05. Not significant p-values (>0.5) are not marked. n= 2-3. Error bars represent SEM. **D**. Titration of Cbl probes with mutated *env*8 variants, each containing either the *anti*PNA (left graph) or *anti*PNA_scr_ fragment (right graph) installed at the 3’ end (c_probe_ = 1 nM, red dot = Cbl-PNA-ATTO590, green square = Cbl-PNA_scr_-ATTO590, blue triangle = Cbl-5xPEG-ATTO590). n= 4. Error bars represent SEM.

Measurement of binding affinities for the mutant *env*8 via fluorescence turn-on studies revealed that the probe containing the complementary PNA linker was able to bind the mutated RNA within the nanomolar range (K_D_ for Cbl-PNA-ATTO590 to *env*8_mut-_FL-3’*anti*PNA was 12.6 ± 1.2 nM and for the scrambled pair was 18.4 ± 1.6 nM) (Figure 4D, Table 2). This result shows a more than 300-fold increase in affinity over the Cbl-5xPEG-ATTO590 probe which bound to the mutant with K_D_ = 3.8 ± 0.3 µM, demonstrating that PNA binding to a complementary sequence in the aptamer significantly enhances affinity between a weak RNA- Cbl binder.

**Table 2.** Summary of all K_D_ values [nM] obtained from fluorescence turn-on studies between mutated variants of wild type *env*8 and cobalamin-based probes. n=4. Error bars represent SEM.

| Probe\RNA | *env8*<sub>mut</sub>-FL-3'antiPNA | *env8*<sub>mut</sub>-FL-3'antiPNA<sub>scr</sub> |
| --- | --- | --- |
| Cbl-PNA-ATTO590 | 12.6 ± 1.2 | 1600 ± 200 |
| Cbl-PNA<sub>scr</sub>-ATTO590 | 2300 ± 400 | 18.4 ± 1.6 |
| Cbl-5xPEG-ATTO590 | 4000 ± 300 | nd |

### The Cbl-PNA conjugate accommodates different fluorescent probes

The modular nature of our synthetic method allows us to install any type of alkyne-modified fluorophore at the end of the PNA strand. To demonstrate the application of an alternative fluorophore, we synthesized the PNA probe with the ATTO488 dye (Supporting Figure S3, Supporting Information 3.8) and measured its affinity towards a set of RNAs (Figure 5A). Like the ATTO590 variant, Cbl-PNA- ATTO488 displayed high affinity binding to the complementary full length *env*8 aptamer (K_D_ 0.12 ± 0.02 nM), the truncated variants and the *env*8 mutant (Figure 5A). The results reveal that the binding affinity is independent of the type of dye, and the probe can be tuned according to spectral needs.

**Figure 5.**
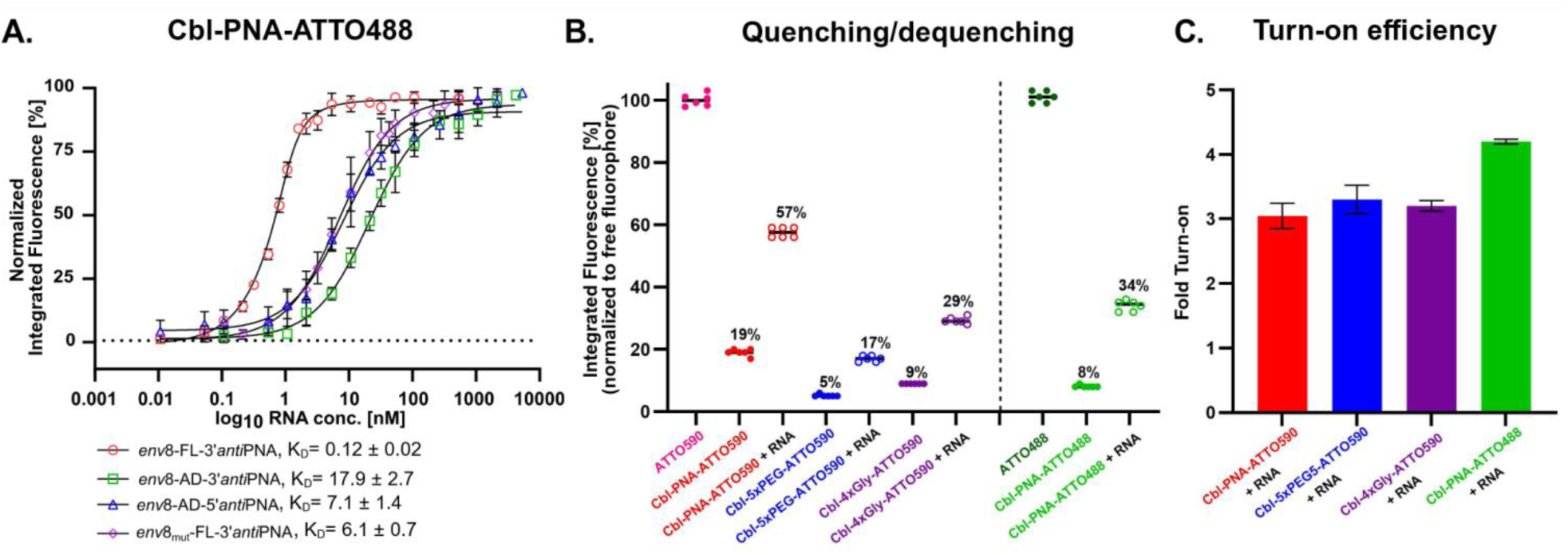
Altering the fluorophore preserves the high affinity of the PNA-based probe and can beneficially influence the photophysical properties of the platform. **A.** Titration of the Cbl-PNA-ATTO488 probe with various RNAs (c_probe_ = 1 nM) and K_D_ values [nM] obtained from fluorescence turn-on studies. n=3-6. Error bars represent SEM. **B.** The levels of probe quenching and dequenching in the presence of *env*8-FL-3’*anti*PNA (c_probe_ = 1 nM, c_RNA_ = 50 nM for Cbl-PNA-ATTO590 and Cbl-PNA- ATTO488, c_RNA_ = 500 nM for Cbl-5xPEG-ATTO590 and Cbl-4xGly-ATTO590). n=6. **C.** Fold turn-on derived from quenching/dequenching studies (RNA = *env*8-FL-3’*anti*PNA). n=4-6. Error bars represent SEM.

The specific fluorophore, however, may have different turn-on properties due to different quenching/dequenching mechanisms. We measured the level of probe quenching in the absence of the RNA and dequenching when the RNA was present and normalized the values to the free fluorophore (Figure 5B). The Cbl-PNA-ATTO590 probe exhibited a higher level of residual fluorescence in its quenched state (19%) compared to the 5xPEG (5%) or 4xGly (9%) probe, likely due to less efficient quenching possibly caused by the increased length and/or increased rigidity of the PNA linker. Nevertheless, once dequenched, the probe turned on by ∼3-fold, comparable to the probes with neutral chemical linkers (Figure 5C). In contrast, the probe with ATTO488 dye exhibited improved turn on (4.2-fold) compared to other linkers (Figure 5C), demonstrating that the quenching/dequenching propensity can be fine-tuned by altering the fluorophore while maintaining a high binding affinity. To verify whether the PNA linker contributes to the quenching process, we synthesized a PNA-ATTO590 conjugate (Supporting Figure S3, Supporting Information 3.10) and tested it against wild type *env*8 in a fluorescence turn-on assay. No significant turn-on was observed, indicating that cobalamin is responsible and essential for quenching the fluorophore (Supporting Figure S8).

### Cbl-PNA probes show improved visualization of RNA in live mammalian cells

While the Cbl-PNA-ATTO590 probe performs better *in vitro* than probes with a neutral linker, we sought to test the probe in live human cells. To accomplish this, we visualized recruitment of mRNA to stress granules (SGs) in live cells; under sodium arsenite stress, cells sequester non-translating cytosolic RNA and proteins into SGs.^33^ This assay has been widely used to test RNA-tagging tools.^13,34–40^ U-2 OS cells stably expressing GFP-G3BP1, a SG marker protein,^41^ were transfected with a plasmid encoding ACTB tagged with four copies of *env*8-FL-3’*anti*PNA. The cells were subsequently beadloaded with 5 µM solution of Cbl-based probe, imaged on a laser scanning confocal microscope, and the enrichment ratio of the probe in SGs relative to the surrounding cytosol was measured (Figure 6A and Supporting Figure S9). Riboglow probes with PNA, 5xPEG, and 4xGly linkers all showed significantly increased enrichment ratios compared to the negative control (untagged RNA), indicating effective binding and recruitment to SGs. However, probes with PNA and 4xGly linkers exhibited higher enrichment ratios and greater statistical significance compared to the 5xPEG probe (Figure 6B, C). All images were collected with the same imaging settings and at the same probe concentration, but 4xGly probe was dimmer in cells than the PNA or 5xPEG probes (data not shown). Therefore, the PNA probe outperformed both the 5xPEG and 4xGly probes in the SG assay.

**Figure 6.**
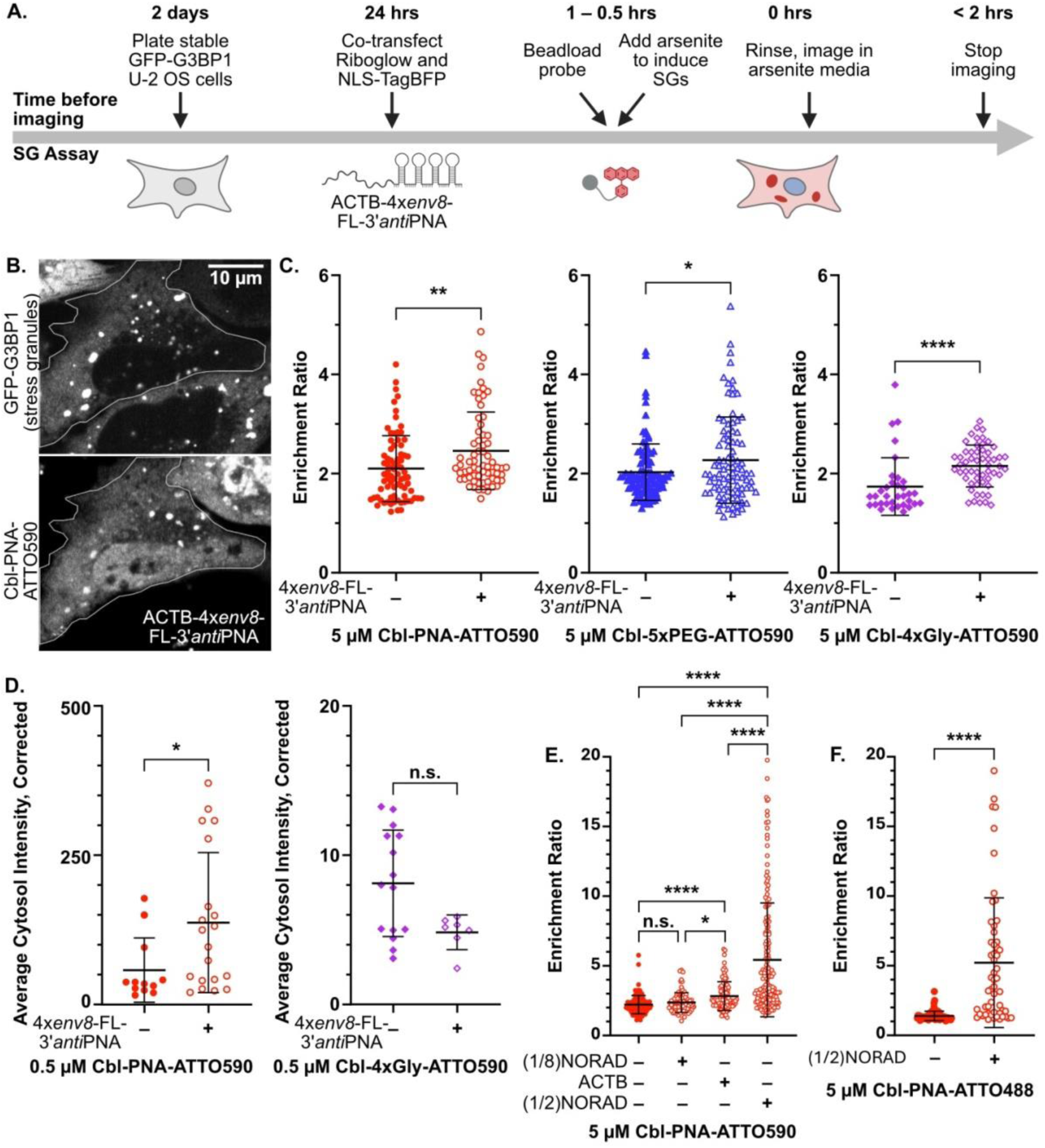
**A**. Schematic showing the timeline of the SG assay. **B**. Representative images of a cell stably expressing GFP-G3BP1, co-transfected with NLS-TagBFP and ACTB-4x*env*8-FL-3’*anti*PNA, and beadloaded with 5 µM Cbl-PNA-ATTO590. Outline of cell in white. Scale bar = 10 µm. **C**. Dot plots of enrichment ratios in cells co-transfected with a fluorescent marker without or with ACTB-4x*env*8-FL-3’*anti*PNA and beadloaded with 5 µM Cbl-PNA-ATTO590, Cbl-5xPEG-ATTO590, or Cbl-4xGly- ATTO590. Each dot represents one SG. n= 34-106 SGs across 7-16 cells in 3-4 imaging dishes for each condition. p-values from Kolmogorov-Smirnov test (nonparametric cumulative distribution t-test). **D**. Dot plots of average cytosol intensity in cells co-transfected with NLS-TagBFP without or with ACTB-4x*env*8-FL-3’*anti*PNA and beadloaded with 0.5 µM Cbl-PNA-ATTO590 or Cbl-4xGly-ATTO590. Each dot represents one cell. n= 7-20 cells across 2-4 imaging dishes for each condition. p-values from Kolmogorov-Smirnov test (nonparametric cumulative distribution t-test). **E**. Dot plots of enrichment ratios in cells co-transfected with NLS-TagBFP without or with an RNA-of-interest tagged with 4x*env*8-FL-3’*anti*PNA and beadloaded with 5 µM Cbl-PNA- ATTO590. Each dot represents one SG. n= 91–170 SGs across 19–29 cells in 3 imaging dishes for each condition. p-values from Kruskal-Wallis test (nonparametric one-way ANOVA) with each condition compared against each other condition. **F**. Dot plots of enrichment ratios in cells co-transfected NLS-TagBFP without or with (1/2)NORAD-4x*env*8-FL-3’*anti*PNA and beadloaded with 5 µM Cbl-PNA-ATTO488. Each dot represents one SG. n= 55–71 SGs across 11 cells in 3–5 imaging dishes for each condition. p-values from Kolmogorov-Smirnov test (nonparametric cumulative distribution t-test). For all panels, bars show mean and standard deviation. p < 0.0001 = ****, p < 0.001 = ***, p < 0.01 = **, p < 0.05 = *, p ≥ 0.05 = n.s.

The Cbl-PNA-ATTO590 probe is brighter when dequenched and has a higher binding affinity *in vitro* compared to probes with nonfunctional linkers (Table 1, Figure 5B). To transfer these advantages to an *in vivo* setting, we decreased the amount of probe 10-fold and performed the SG assay. The 4xGly probe was too dim to analyze the enrichment ratio, so instead we compared the percent colocalization of the probe with SGs. The PNA probe had a much higher percent of colocalization with the SG marker compared to 4xGly probe (mean of 58% versus 15%, respectively, Supporting Figure S10). The average cytosol intensity of cells with PNA probe was higher in the presence of tagged RNA, showing fluorescence turn-on of the PNA probe upon binding (Figure 6D). This result is consistent with findings from another study using different Riboglow probes.^40^ With the same imaging settings, the average cytosol intensity of cells with 4xGly probe did not show a difference in the presence of tagged RNA (Figure 6D). The increased brightness and higher affinity of the PNA probe enable it to work at low probe concentrations while the probe with nonfunctional 4xGly linker fails.

While recruitment of ACTB to SGs has been widely used to evaluate RNA imaging tools,^13,35–40^ a study that purified SGs and sequenced their RNA revealed that only a small percent of endogenous ACTB RNA localizes to SGs.^42^ To increase the dynamic range in the SG assay, we tested (1/2)NORAD, the first half of the NORAD ncRNA which was shown to be strongly localized to SGs during stress.^40,43^ As a negative control, we examined (1/8)NORAD, the first eighth of the NORAD ncRNA which has weaker SG colocalization. All RNAs used in this assay have the same 4x*env*8-FL-3’*anti*PNA array appended to their 3’ end. The enrichment ratio of the Cbl-PNA-ATTO590 probe increased significantly in the presence of (1/2)NORAD, far above ACTB, but remained unchanged in the presence of (1/8)NORAD (Figure 6E). In addition to demonstrating that the PNA probe functions with other RNA targets, we show that (1/2)NORAD is a more robust target than ACTB in the SG assay, leading to higher enrichment in SGs, as has been shown recently with FLIM and other Cbl probes.^40^

Because the RNA aptamer does not bind the fluorophore directly, the fluorophore can be easily exchanged. The Cbl-PNA-ATTO488 probe performs on par with the Cbl-PNA-ATTO590 *in vitro*, so we also tested Cbl-PNA-ATTO488 probe in live cells. Cbl-PNA-ATTO488 performs similarly to Cbl-PNA-ATTO590 and shows increased enrichment in SGs in the presence of (1/2)NORAD-4x*env*8-FL-3’*anti*PNA (Figure 6F). As was shown with the *in vitro* experiments, the probe fluorophore can be switched without altering the functionality of the system *in vivo*.

The Cbl-PNA-ATTO590 probe confers more than 200-fold increased affinity *in vitro* for the truncated aptamer by base-pairing with a complementary region (Figure 3, Table 1). Therefore, we tested whether PNA could provide a distinct advantage for tracking RNAs with a truncated aptamer in live cells using a U-body assay. Under thapsigargin stress, cells accumulate small noncoding U-rich spliceosomal RNAs and associated proteins into cytoplasmic U-bodies.^44,45^ Previously, we used the truncated aptamer *env*8-AD-U1 and a Cbl-5xPEG-ATTO590 probe to track enrichment of the U1 snRNA into U-bodies upon stress.^13^ Importantly, this small snRNA cannot tolerate large fusions without perturbing its processing,^46^ making the truncated aptamer essential. In this U-body assay, we transfected U-2 OS cells with GFP-SMN1, a U-body marker protein,^44^ and tagged U1 with either *env*8-AD or *env*8-AD-5’*anti*PNA. The cells were then beadloaded with 50 μM PNA probe, imaged using a laser scanning confocal microscope, and analyzed for colocalization of the PNA probe with the U-body marker (Figure 7A, Supporting Figure S11). A significantly higher percentage of the PNA probe colocalized with U- bodies in the presence of *env*8-AD-5’*anti*PNA-U1 compared to *env*8-AD-U1 and cells with *env*8- AD-5’*anti*PNA showed very high colocalization between PNA probe and U-bodies (Figure 7B, C). Binding of the Cbl portion alone (*env*8-AD-U1) is sufficient to observe colocalization, but the further increase in colocalization in cells transfected with RNA capable of engaging the PNA linker (*env*8-AD-U1-5’*anti*PNA) strongly suggests that the PNA probe’s sequence-specific binding behavior observed *in vitro* is recapitulated in live cells.

**Figure 7.**
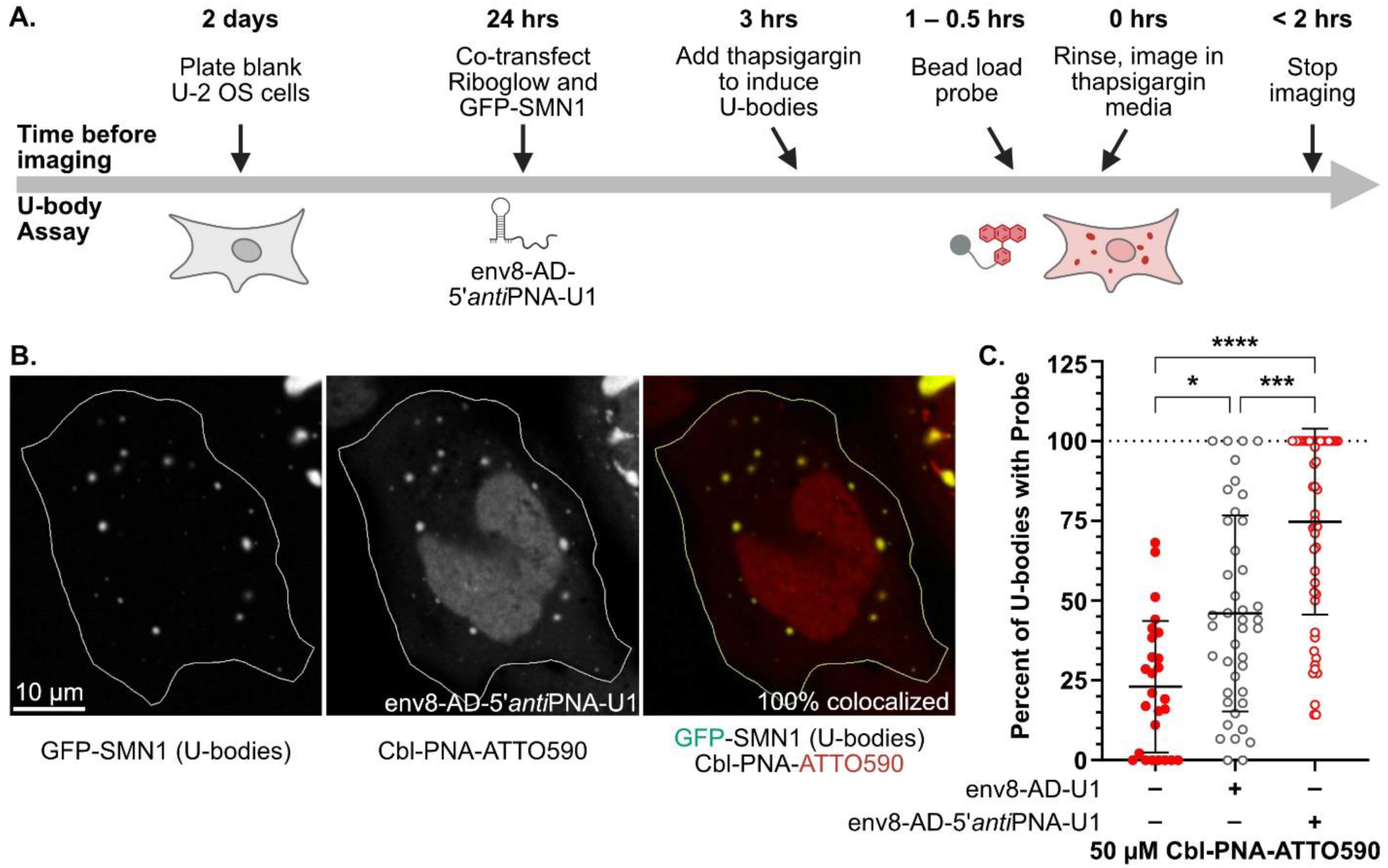
**A.** Schematic showing the timeline of the U-body assay. **B.** Representative images of a cell co-transfected with GFP- SMN1 and *env*8-AD-5’*anti*PNA and beadloaded with 50 µM Cbl-PNA-ATTO590. In merged image, GFP-SMN1 (U-body) channel in green, Cbl-PNA-ATTO590 in red, and overlap in yellow. Outline of cell in white. Scale bar = 10 µm. **C.** Dot plot of percent of U-bodies that show visible colocalization in cells co-transfected with GFP-SMN1 without or with *env*8-AD-U1 or *env*8-AD-5’*anti*PNA-U1 and beadloaded with 50 µM Cbl-PNA-ATTO590. Each dot represents one cell. n= 26-52 cells across 3 imaging dishes for each condition. Bars show mean and standard deviation. p-values from Kruskal-Wallis test (nonparametric one-way ANOVA) with each condition compared against each other condition. p < 0.0001 = ****, p < 0.001 = ***, p < 0.01 = **, p < 0.05 = *, p ≥ 0.05 = n.s.

## Discussion

Linkers are vital in chemical biology, connecting functional groups, molecules, and biomolecules to form complex, multifunctional systems crucial for studying biological processes. However, their role extends well beyond mere connectivity, as their intrinsic chemical properties can be leveraged to enhance the stability, reactivity, and functionality of these systems.^14^ In our work, we explored the possibility of improving the affinity and specificity of chemical probes targeting RNA by incorporating a linker capable of interacting with the RNA target with high specificity. To achieve this, we integrated a PNA-based linker into the small molecule probe, which functions through contact-quenching. The PNA linker serves as an additional anchoring point for the probe, augmenting the RNA-ligand (Cbl) structure-based recognition. Therefore, by incorporating the chemical functionality within the linker, we enable two distinct binding events to operate in concert, resulting in increased affinity and specificity for the RNA imaging platform.

Significantly tighter binding was observed for PNA probes compared to probes with neutral linkers, and the combined involvement of the two binding interactions (Cbl-RNA and PNA linker-complementary RNA) was confirmed through chemical probing. The difference in affinity was even more pronounced when the RNA aptamer was truncated. Additionally, the effect was preserved regardless of whether the PNA binding site is installed at the 5’ or 3’ end of the aptamer, indicating the adaptability of the approach. *In vivo* studies demonstrated the advantages of PNA probes, allowing for more robust visualization of RNA recruitment to stress granules and U-bodies. The increased affinity and brightness allowed the PNA probes to be used at much lower concentrations compared to those with non-functional linkers. Additionally, we have shown that the PNA-RNA sequence-specific interaction observed *in vitro* is also successfully occurring in live cells. We demonstrated that a linker as short as six nucleobases can effectively restore binding to a significantly impaired aptamer, increasing affinity by over 300- fold compared to a probe with a non-functional 5xPEG linker. This proves that specificity of the probe can be driven by the PNA linker. Moreover, this property can be utilized to achieve orthogonality by designing multiple aptamers with significantly different affinities and targeting them selectively with distinct PNA probes. This strategy paves the way for simultaneous multicolor tracking of various RNA types, a concept we intend to investigate further in the future.

The Riboglow platform can be readily adapted, and with the installation of the PNA linker, its versatility expands even further. We have demonstrated that both the PNA linker and the fluorophore can be modified while preserving high affinity and specificity. Additionally, the RNA tag itself can be altered to match the sequence present within the linker and/or modify the ligand interaction. However, there is still room for improvement and exploration. In this context, special emphasis can be placed on the modularity of PNAs, as their capabilities are not limited to the selective recognition of single-stranded RNAs. PNAs can be chemically modified^47^ and altered nucleobases can serve as a handle for bioactive molecule installation (e.g. fluorophore).^48^ Specifically designed PNAs can recognize double-stranded regions (triplex formation), form triplex invasion structures, duplex invasion, and double-duplex invasion structures.^49–51^ Furthermore, the synthetic nature of PNA allows for non-canonical pairings to optimize PNA-RNA interactions.^52^ Therefore, we believe that combining a small molecule with a PNA strand has the potential to enhance existing RNA imaging tools that rely on collisional quenching, as well as inspire the development of new ones.

In this work we establish a new approach for improving RNA imaging tools and demonstrate the potential for using short oligonucleotides conjugated to small molecules to improve affinity and specificity of chemical probes targeting RNA. Apart from new opportunities to improve RNA imaging tools, we speculate that the approach described here could also be exploited in RNA-targeted drug development. Though much effort has been devoted to targeting RNA, the field still remains in its infancy.^53–55^ Two main therapeutic strategies including antisense oligonucleotides (∼20-30 nucleotides) and small molecules have been employed to target disease-causing RNAs.^56,57^ Among small molecules, bifunctional structures such as dimeric miRNA binders enhance potency and selectivity by simultaneously targeting a functional site and an adjacent druggable motif.^56^ On the other hand, RNA cleavers and degraders, such as bleomycin A5-conjugates and RIBOTACs, offer novel therapeutic modalities to reduce levels of disease-causing RNAs.^56^ These examples support the importance of introducing more than one functionality within the RNA-targeting molecule to achieve the desired effects. Similarly, relying on just one binding event may not be sufficient and could result in binding promiscuity. Synthetic oligonucleotides open new possibilities in this area and can significantly improve specific RNA targeting. A recent example on PNA-based fluorogenic probes for sensing of the panhandle structure of the influenza A virus (IAV) RNA promoter region utilizes the PNA’s ability to form triplexes with double-stranded RNA region.^48^ Consistent with our findings, conjugation of 8-mer PNA sequence equipped with a fluorogenic nuclobase to 7-dimethoxy-2-(1-piperazinyl)-4- quinazolinamine (DPQ) improved affinity toward RNA compared to the small molecule alone. Therefore, in the pursuit of more defined RNA-small molecule interactions, not only for imaging purposes, the combination of two different binding events operating in concert is essential. We foresee that the combination of a small molecule targeting RNA via structure-based recognition, along with a sequence-specific synthetic oligonucleotide serving as an additional anchoring point, could advance the field of RNA drugging further.

## Conclusions

Our work demonstrates the potential of using short peptide nucleic acid (PNA)-based linkers to enhance RNA-targeting chemical probes. By integrating a PNA linker into the Riboglow platform, we achieved significant improvements in probe affinity and specificity through dual binding events. The PNA linker allows for stronger, specific binding, enhancing probe performance both *in vitro* and *in vivo*. These improvements enable lower probe concentrations and more robust visualization of RNA dynamics in cellular environments.

Our findings highlight the versatility of PNA linkers in RNA imaging platforms and suggest broader applications, including the design of orthogonal probes for multicolor tracking of different RNA species. Additionally, the adaptability of the Riboglow platform, combined with the tunable nature of PNAs, provides a pathway for further enhancing RNA imaging tools and developing new strategies for targeting RNA.

We also propose that this approach can be extended beyond imaging to RNA-targeted drug development. The ability to combine multiple binding interactions within a single probe opens new avenues for improving the selectivity and potency of RNA-targeting molecules. This dual-binding approach represents a promising strategy for advancing RNA drugging and holds potential for future innovations in both research and therapeutic applications.

## Author contribution

AJW: synthesis and characterization of the probes, RNA synthesis and purification, fluorescence turn-on assay, data analysis

EMR: preparation of in vivo constructs, in vivo imaging and quantification

SRL: RNA synthesis and purification, SHAPE experiments and quantification

AJW, RTB, AEP: conceptualization of the project

AJW, EMR, SRL, RTB, AEP: manuscript preparation

RTB, AEP: supervision and funding

The manuscript was written through contributions of all authors. All authors have given approval to the final version of the manuscript.

### Notes

The authors declare the following competing financial interest(s): R.T.B. serves on the Scientific Advisory Boards of Expansion Therapeutics, SomaLogic, and MeiraGTx.

## Acknowledgements

We would like to acknowledge the following financial support: R01 GM133184 (AEP and RTB), R35 GM139644 (AEP), R35 GM152029 (RTB), T32 GM065103 (EMR, SRL), F31 5F31ES033919 (EMR), Polish National Agency for Academic Exchange PPN/BEK/2020/1/00219/U/0000 (AJW).

We thank the Shared Instruments Pool (RRID: SCR_018986) of the Department of Biochemistry at the University of Colorado Boulder for the use of the Typhoon 5. The Typhoon 5 is funded by NIH Shared Instrumentation Grant S10OD034218-01.

The imaging work was performed at the BioFrontiers Institute’s Advanced Light Microscopy Core (RRID: SCR_018302). Laser scanning confocal microscopy was performed on an Nikon A1R microscope supported by NIST-CU Cooperative Agreement award number 70NANB15H226. We would like to thank Dr. Joe Dragavon of the BioFrontiers Institute Advanced Light Microscopy Core for assistance with microscopy.

We would like to acknowledge the University of Colorado Biochemistry Cell Culture Core Facility, especially Dr. Theresa Nahreini, for providing resources and support for all our cell work.

We would like to thank Dr. Esther Braselmann for helpful comments regarding the stress granule assay.

## Supporting information

Experimental details for chemical probe synthesis and characterization are provided in supporting information. In addition, we provide the crystal structure of the *env*8 aptamer highlighting the PNA binding site, chemical structures of the PNA linkers and Cbl probes, secondary structures of the RNAs, full SHAPE gels and associated quantification, pipeline for SG assay, quantification of percent SGs with probe, pipeline for U-body assay, and validation of *env*8 RNA.

## Methods

### Probe synthesis

Synthesis of fluorophore probes. Synthesis and characterization of probes are described in Supporting Information, see Supporting information: probe synthesis and characterization.

### RNA synthesis and purification

For all *in vitro* experiments, RNAs were synthesized using template DNA amplified through PCR. DNA templates were amplified using a series of overlapping oligonucleotides ordered from IDT (Supporting Table S2). Transcriptions used T7 RNA polymerase (for details see Supporting Table S3).^58^ The RNAs were purified using denaturing PAGE (8% acrylamide:bisacrylamide 29:1, 8 M urea, 1x Tris-Borate-EDTA (TBE) buffer). This was polymerized with 10% (w/v) ammonium persulfate (500 µL for 100 mL of gel) and tetramethylethlyenediamine (TEMED, 50 µL for 100 mL of gel). The gel was preheated at 14 W for about 30 minutes prior to loading. 1x TBE buffer was used as running buffer. RNAs were mixed with an equal volume of denaturing dye (85% formamide, 0.5x TBE, 50 mM EDTA, pH 8.0, with xylene cyanol and bromophenol blue tracking dyes) and heated at 90° C for 3 minutes prior to loading without cooling. RNA samples were loaded and the gel ran for about 2.5 hours at 14 W at room temperature. *In vitro* transcription of wild type *env*8 RNA led to two major products (Supporting Figure S12). Analysis of Cbl binding to the upper band, the lower band, or a mixture of two bands revealed the lower band to be the correct species (Supporting Figure S12). All subsequent binding experiments were conducted with the purified lower band. Full length transcripts were visualized by UV shadowing, excised from the gel, and eluted from the gel by soaking at 4 °C in MilliQ water. RNA was concentrated into MilliQ water using centrifugal concentrators (Amicon) with 10 KDa molecular weight cutoff. Final RNA concentrations were calculated using A260 and molar extinction coefficients determined from the summation of the individual bases. For selective 2’-hydroxyl acylation analyzed by primer extension (SHAPE) assays, RNA was gel purified twice and stored at -80 °C until use. Other gel purified RNA was purified once and stored at -20 °C until use. Secondary structures and sequences of all riboswitch RNAs used in this study are shown in Supporting Figure S4 and Supporting Table S1, respectively.

### SHAPE experiments

Chemical structure probing was performed using *N*-methylisatoic anhydride (NMIA) as previously described^31^ with modifications. RNAs were synthesized with additional 5’ and 3’ sequences, called the SHAPE cassette, for purposes of reverse transcription and visualization.^31^ RNAs were refolded by heating at 90 °C for 3 minutes and placed on ice for a minimum of 10 minutes. RNA was added to folding buffer (333 mM Na-HEPES pH 8.0, 333 mM NaCl, 20 mM MgCl_2_) along with a ligand (dissolved in DMSO) and incubated at room temperature for 10 minutes before adding NMIA dissolved in DMSO or neat DMSO for control reactions. The final 10 μL probing reactions had concentrations of 100 nM RNA, 100 mM Na-HEPES pH 8.0, 100 mM NaCl, 6 mM MgCl_2_, 30 μM ligand unless stated otherwise, and 13 mM NMIA. Probing reactions were incubated at 37 °C for 41 minutes (5 half-lives). Following probing, reactions were moved straight into primer extension using reverse transcriptase as previously described^59^ without ethanol precipitation. A ^32^P 5’-end labeled reverse transcription primer complementary to the 3’ SHAPE cassette was used (Supporting Table S2) along with the enzyme mix (250 mM KCl, 167 mM Tris-HCl at pH 8.3, 1.67 mM each dNTP, 17 mM DTT, 0.33 units Superscript III).^59^ Products were run on a 12% denaturing polyacrylamide gel at 55 W for several (4-6) hours until the areas of interest could be resolved. Gels were exposed to a storage phosphor screen at least overnight and for up to two weeks, and screens were imaged using a Typhoon FLA 9500 PhosphoroImager (GE) or an Amersham Typhoon 5 (Cytiva). Gel images were straightened and quantified by the SAFA software package.^60^ Data were also normalized by the software, using the built-in invariant residue determination. This normalizes all sequencing gel band intensities to those of nucleotides which do not show any change in response to ligand, to account for any loading differences. Finally, degree of protection (DOP) was calculated using the equation 𝐷𝑂𝑃 = –((𝐶−𝑁)/𝑁) where C is the sum of all bands of a region of interest in the Cbl lane and N is the sum of all bands of a region of interest in the NMIA lane.

### Fluorescence turn-on experiments

All *in vitro* experiments were conducted in RNA buffer (100 mM KCl, 10 mM NaCl, 1 mM MgCl_2_, 50 mM HEPES, pH 8.0) with the addition of 0.01% nonidet P40 and 10% DMSO (v/v). For the Cbl-fluorophore probes, the extinction coefficient of the fluorophores was used to determine the concentration (120000 M^-1^cm^-1^ for ATTO590 and 90000 M^-1^cm^-1^ for ATTO488, source: ATTO-TEC). To determine the binding affinity of each probe to the RNA of interest, a series of titration experiments were performed. Relevant amounts of RNA were titrated into 120 μL reactions into Eppendorf tubes, ensuring a final concentration of 1 nM cobalamin probe (for details, see Supporting Table S4). All reactions were performed in technical duplicate and in at least biological triplicate. The reactions were pipetted into a Corning 384-well plate (2 x 55 μL). The reactions were allowed to equilibrate for 60 minutes at room temperature in the dark before reading. Extended equilibration times (up to 24 hrs) did not significantly alter the K_D_ for the titration of *env*8-FL-3’*anti*PNA and Cbl-PNA-ATTO590, therefore we settled on the 60 min time point. ATTO590 fluorescence was excited at 590 ± 8 nm and fluorescence emission was from collected from 615-675 nm using a BMG Labtech CLARIOstarPLUS microplate reader. ATTO488 was excited at 485 ± 5 nm and fluorescence emission was from collected from 505-565 nm using the same instrument. Fluorescence values were background corrected by subtracting the fluorescence values of a buffer control at each wavelength and integrated over all wavelengths. The corrected and integrated fluorescence values were plotted versus log(nM RNA) in GraphPad Prism. Technical replicates were fit to the quadratic binding equation with one transition 𝑌 = 𝑚 + (𝑛 – 𝑚) ∗ (((𝑐 + 𝑥 + 𝐾) – 𝑠𝑞𝑟𝑡(𝑠𝑞𝑟(𝑐 + 𝑥 + 𝐾) – (4 ∗ 𝑐 ∗ 𝑥)))/(2 ∗ 𝑐)) where Y is the corrected integrated fluorescence value, m is the lower baseline, n is the upper baseline, c is the probe concentration, x is the RNA concentration, and K is the K_D_.^32^ Calculated K_D_s are the average of multiple biological replicates, and errors were calculated from those average K_D_s. Biological replicates were combined for the final graphs shown in the figures. To combine replicates and calculate the fraction of probe bound to RNA, each was normalized to values between 0 and 100, where 0 was the value of the lower baseline and 100 was the value of the upper baseline as calculated by the quadratic binding equation. The fold turn-on of 1 nM probes was calculated by dividing the fluorescence value obtained from titration point at c_RNA_ = 50 nM (for Cbl-PNA-ATTO590 and Cbl-PNA-ATTO488) and c_RNA_ = 500 nM (for Cbl-5xPEG-ATTO590 and Cbl-4xGly-ATTO590) by the fluorescence value obtained for the free probe. Same fluorescence values were used to determine quenching/dequenching levels for each probe. Quenching levels were derived from titration point at c_RNA_ = 50 nM (for Cbl-PNA- ATTO590 and Cbl-PNA-ATTO488) and c_RNA_ = 500 nM (for Cbl-5xPEG-ATTO590 and Cbl-4xGly- ATTO590) while dequenching levels from fluorescence values obtained for free probe at 1 nM. Each value was divided by the fluorescence level obtained for the free fluorophore (c_free dye_ = 1 nM) to obtain quenching/dequenching levels. Data were normalized to values between 0 and 100, where 0 was the value of the lower baseline and 100 was the value of the upper baseline.

### Cloning plasmids for mammalian expression

To create PB-HaloTag-ACTB-0x, pUC57-HaloTag-ACTB-18x*env*8-FL was ordered as a custom plasmid from GeneWiz. Lyophilized plasmid was dissolved and transformed into C3040 recombinase-deficient *E. coli* chemically competent cells (NEB). Colonies were selected on LB+Amp agar plates and plasmid was isolated for downstream use (Qiagen Midiprep Kit). pUC57-HaloTag-ACTB-18x*env*8-FL and PB510B-1 (System Biosciences) were digested with the unique restriction sites SwaI and XbaI (NEB). Fragments were gel purified (Macherey Nagel Gel and PCR Cleanup Kit) and ligated with T4 DNA ligase (NEB). The ligation reaction was then transformed into C3040 cells and selected on LB+Amp agar plates. Colonies were screened for the insertion and sent for sequencing. PB-HaloTag-ACTB-18x*env*8 was then digested with the unique restriction sites BlpI and PmeI (NEB) to remove the 18x*env*8 fragment, blunted with DNA polymerase I Klenow fragment (Thermo Fisher Scientific), and ligated with T4 DNA ligase. The ligation reaction was then transformed into C3040 cells and selected on LB+Amp agar plates. Colonies were screened for deletion of the array and sent for whole plasmid nanopore sequencing.

To create *env*8-AD-5’*anti*PNA-U1, *env*8-AD-U1 (Addgene plasmid #112059) was first transformed into C2925 *dam-/dcm- E. coli* chemically competent cells (NEB) and selected on LB+Kan agar plates. Plasmid was isolated for downstream use. A custom gBlock of *env8*-AD-5’*anti*PNA and primers to amplify it were ordered from IDT (Supporting Table S2). *env*8-AD-5’*anti*PNA was PCR amplified, ethanol precipitated, digested with the unique restriction sites BglII and BclI (NEB), and purified. *env8*-AD-U1 was digested with the unique restriction sites BglII and BclI, CIP (NEB) treated, and gel purified. Fragments were ligated with T7 DNA ligase (NEB), transformed into Stellar *E. coli* chemically competent cells (Takara Bio), and selected on LB+Kan agar plates. Colonies were screened and sent for whole plasmid nanopore sequencing.

### Cell lines and cell culture

U-2 OS cells were obtained from ATCC (ATCC number HTB-96) and cultured in McCoy’s 5A medium supplemented with 10% FBS and 1% P/S. Cells were routinely passaged into 10-cm cell culture-treated plates before they reached confluency and were mycoplasma free for the course of these experiments. All cells used in the following experiments were used on or before their 25th passage. The GFP-G3BP1 stable U-2 OS cells and the HaloTag-G3BP1 CRISPR stable U-2 OS cells were generated previously.^13^

### Stress granule (SG) assay setup and imaging

175,000 U-2 OS cells stably expressing GFP-G3BP1 or HaloTag-G3BP1 were seeded into home-made 3.5-cm cell culture-treated imaging dishes with a ∼10 mm center hole covered by cover glass (No. 1.5, VWR) 2 days before imaging. About 24 hours before imaging, cells were transfected using TransIT (Mirus Bio) and ∼292 fmol ACTB-4x*env*8-FL-3’*anti*PNA (as described previously,^13^ Addgene plasmid #112058), (1/2)NORAD-4x*env*8-FL-3’*anti*PNA (Addgene plasmid #199208), or (1/8)NORAD-4x*env*8- FL-3’*anti*PNA (Addgene plasmid #199209). For all experiments, unless noted otherwise, NLS-TagBFP (Addgene plasmid #55265) was used in an equimolar amount as a co-transfection marker which was shown to co-express from the same cell as the other transfected plasmid 94% of the time.^13^ In the 5 µM Cbl-4xGly-ATTO590 experiments only, an equimolar amount of PB-HaloTag-ACTB-0x was used instead as a co-transfection marker. On the day of imaging, cells were beadloaded as previously described^13^ with 3 µL of 5 µM or 0.5 µM probe (2% DMSO, 98% D-PBS by volume). 2 mL of cell culture media with 0.5 mM sodium arsenite (Sigma-Aldrich) was added immediately after beadloading to induce stress granules. For the 5 µM Cbl-4xGly-ATTO590 experiments only, 10 nM HaloTag-JF669 (Lavis Labs) and 2 drops of NucBlue live cell stain (Life Technologies) were also added at the time of beadloading to stain the HaloTag-ACTB co-transfection protein, and illuminate the nucleus, respectively. For the 5 µM Cbl-PNA- ATTO488 experiments, cells were beadloaded with 6 µL of 5 µM probe and 100 nM HaloTag-JF669 ligand and 2 drops of NucBlue live cell stain (Life Technologies) were added at the time of beadloading to stain the HaloTag-G3BP1 protein and illuminate the nucleus. Cells were allowed to recover at 37 °C and 5% CO_2_ for 30 minutes to 1 hour. Then cells were prepped for imaging by removing the media, rinsing once with 1 mL FluoroBrite DMEM (Fisher) supplemented with 10% FBS and 0.5 mM sodium arsenite, and imaged in 1 mL FluoroBrite DMEM supplemented with 10% FBS and 0.5 mM sodium arsenite. Cells were then imaged in a LiveCell stage top environmental chamber (Pathology Devices, Inc.) at 37 °C, 5% CO_2_, and 95% humidity for up to 2 hours. Images were collected on a Nikon Ti-E A1R laser scanning confocal microscope with a 100X (1.45 NA) Plan Apo Lambda oil objective (Nikon) and 405-nm (Coherent OBIS), 488-nm (Coherent OBIS), 561-nm (Coherent Sapphire), and 640-nm (Coherent OBIS) lasers. Signal from excitation with 405-nm and 488-nm lasers were collected with Nikon PMT detectors and signal from excitation with 561-nm and 640-nm lasers were collected with GaAsp PMT detectors. Imaging settings are defined in Supporting Table S5. Expression of transfected plasmids for longer than 30 hours or collection of a z-stack that was then maximum-intensity projected for analysis resulted in significantly decreased dynamic range between the probe alone and plus aptamer RNA conditions.

### Stress granule (SG) assay analysis

Confocal images as .nd2 files were imported into Fiji/ImageJ using the Bio-Formats Importer. Channels were separated and the expression of the co-transfection marker and presence of beadloaded probe were confirmed. The line selection tool was used to draw a line across a SG in the SG marker channel, save the selection in the ROI Manager, and make a line plot of the fluorescence intensities with Plot Profile. The ROI manager was used to bring up the same line selection in the probe channel and make a line plot of the fluorescence intensities with Plot Profile. The two line profiles were aligned and two max probe intensities inside the SG (as defined by high SG marker intensity) were recorded in an Excel spreadsheet with the image date, construct, probe identity, probe concentration, cell number, and SG number. The same was done for two average probe intensities outside of the SG (as defined by low SG marker intensity). Line profiles of each SG in the cytosol with well-defined edges and larger than 2 px-by-2 px were collected, and probe intensities inside and outside the SG were recorded. To correct for imaging media fluorescence and detector background noise, three boxes larger than 15 px-by-15 px were drawn outside of cells in the image using the rectangle selection tool. The mean probe intensity of each background box was determined with Measure and recorded in the Excel spreadsheet for that image. The average background probe intensity, the average SG max probe intensity, the average local cytosol probe intensity, the background-corrected average SG max probe intensity, the background-corrected average local cytosol probe intensity, and the enrichment ratio were calculated in Excel with the equations in Supporting Figure S8B. For each imaging set, the enrichment ratio was plotted as a function of the background-corrected average local cytosol probe intensity. Thresholds for the background-corrected average local cytosol probe intensity were determined by the shape of the data, setting the lower threshold to remove high enrichment values caused by a small denominator and setting the higher threshold to remove low enrichment values caused by a large denominator. Statistical tests were only performed on data sets that used the same imaging settings and threshold values. A Q-test was performed on each data set to determine if the highest or lowest enrichment ratio should be removed from the thresholded data set. No more than one data point was removed from each thresholded data set. A Kolmogorov-Smirnov test (nonparametric cumulative distribution t-test) was performed on graphs with two data sets, and a Kruskal-Wallis test (nonparametric one-way ANOVA) was performed with each condition compared against each other condition on graphs with more than two data sets. Graphing of data and statistical tests were done in GraphPad Prism.

### U-body assay setup and imaging

175,000 U-2 OS cells were seeded into home-made 3.5-cm cell culture-treated imaging dishes with a ∼10 mm center hole covered by cover glass (No. 1.5, VWR) 2 days before imaging. About 24 hours before imaging, cells were transfected using TransIT (Mirus Bio) and ∼448 fmol *env8*-AD-U1 as described previously,^13^ Addgene plasmid #112059) or *env*8-AD-5’*anti*PNA-U1 (described above). ∼65 fmol EGFP- SMN1 (Addgene plasmid #37057) was used as a co-transfection marker and a U-body marker protein. On the day of imaging, 1 mL of cell culture media with 10 µM thapsigargin (Calbiochem/VWR) was added 3 hours before imaging to induce U-bodies. One hour before imaging, cells were beadloaded as previously described^13^ with 3 µL of 50 µM probe (25% DMSO, 75% D-PBS by volume). 1 mL of cell culture media with 10 µM thapsigargin and 2 drops of NucBlue live cell stain (Life Technologies) were added immediately after beadloading to keep inducing U-bodies and to stain the nucleus. Cells were allowed to recover at 37 °C and 5% CO_2_ for 30 minutes to 1 hour. Cells were prepped for imaging by removing the media, rinsing once with 1 mL FluoroBrite DMEM (Fisher) supplemented with 10% FBS and 10 µM thapsigargin. Cells were imaged in 1 mL FluoroBrite DMEM supplemented with 10% FBS and 10 µM thapsigargin in a LiveCell stage top environmental chamber (Pathology Devices, Inc.) at 37 °C, 5% CO_2_, and 95% humidity for up to 2 hours. Images were collected on a Nikon Ti-E A1R laser scanning confocal microscope with a 100X (1.45 NA) Plan Apo Lambda oil objective (Nikon) and 405-nm (Coherent OBIS), 488-nm (Coherent OBIS), 561-nm (Coherent Sapphire), and 640-nm (Coherent OBIS) lasers. Signal from excitation with 405-nm and 488-nm lasers were collected with Nikon PMT detectors and signal from excitation with 561-nm and 640-nm lasers were collected with GaAsp PMT detectors. Imaging settings are defined in Supporting Table S5.

### U-body assay analysis

Confocal images as .nd2 files were imported into Fiji/ImageJ using the Bio-Formats Importer. Channels were separated and the expression of the co-transfection marker/U-body marker protein and presence of beadloaded probe were confirmed. The U-body marker protein channel was initialized in Cell Counter and channels were synced with Synchronize Windows. One counter type in Cell Counter was used to mark U-bodies that visibly had probe and another counter type was used to mark U-bodies that did not visibly have probe. This was repeated for all U-bodies in the cytosol with well-defined edges and larger than 2 px-by-2 px. The number of U-bodies with visible colocalization and the number of U-bodies with no visible colocalization were recorded in an Excel spreadsheet with the image date, construct, probe identity, probe concentration, and cell number. For each cell, three boxes larger than 4 px-by-4 px were drawn inside the cytosol using the rectangle selection tool. The mean probe intensity of each cytosol box was determined with Measure and recorded in Excel. The percent visible colocalization was calculated with the equation in Supporting Figure S11 in Excel. For each imaging set, the percent visible colocalization was plotted as a function of average cytosol intensity, but it was determined that no threshold for the average cytosol intensity was required. Statistical tests were only performed on data sets that used the same imaging settings. A Q-test was performed on each data set to determine if the highest or lowest percent visible colocalization should be removed from the data set. No more than one data point was removed from each thresholded data set. A Kruskal-Wallis test (nonparametric one-way ANOVA) was performed with each condition compared against each other condition. Graphing of data and statistical tests were done in GraphPad Prism.

## References

(1) Dean, K. M.; Palmer, A. E. Advances in Fluorescence Labeling Strategies for Dynamic Cellular Imaging. Nat. Chem. Biol. 2014, 10 (7), 512–523.

(2) Tutucci, E.; Livingston, N. M.; Singer, R. H.; Wu, B. Imaging MRNA in Vivo, from Birth to Death. Annu. Rev. Biophys. 2018, 47, 85–106.

(3) Tutucci, E.; Vera, M.; Biswas, J.; Garcia, J.; Parker, R.; Singer, R. H. An Improved MS2 System for Accurate Reporting of the MRNA Life Cycle. Nat. Methods 2018, 15 (1), 81– 89.

(4) Wu, B.; Miskolci, V.; Sato, H.; Tutucci, E.; Kenworthy, C. A.; Donnelly, S. K.; Yoon, Y. J.; Cox, D.; Singer, R. H.; Hodgson, L. Synonymous Modification Results in Highfidelity Gene Expression of Repetitive Protein and Nucleotide Sequences. Genes Dev. 2015, 29 (8), 876–886.

(5) Garcia, J. F.; Parker, R. Ubiquitous Accumulation of 3′ MRNA Decay Fragments in Saccharomyces Cerevisiae MRNAs with Chromosomally Integrated MS2 Arrays. Rna 2016, 22 (5), 657–659.

(6) Garcia, J. F.; Parker, R. MS2 Coat Proteins Bound to Yeast MRNAs Block 5′ to 3′ Degradation and Trap MRNA Decay Products: Implications for the Localization of MRNAs by MS2-MCP System. Rna 2015, 21 (8), 1393–1395.

(7) Braselmann, E.; Rathbun, C.; Richards, E. M.; Palmer, A. E. Illuminating RNA Biology: Tools for Imaging RNA in Live Mammalian Cells. Cell Chem. Biol. 2020, 27 (8), 891– 903.

(8) Le, P.; Ahmed, N.; Yeo, G. W. Illuminating RNA Biology through Imaging. Nat. Cell Biol. 2022, 24 (6), 815–824.

(9) Sunbul, M.; Lackner, J.; Martin, A.; Englert, D.; Hacene, B.; Grün, F.; Nienhaus, K.; Nienhaus, G. U.; Jäschke, A. Super-Resolution RNA Imaging Using a Rhodamine-Binding Aptamer with Fast Exchange Kinetics. Nat. Biotechnol. 2021, 39 (6), 686–690.

(10) Sunbul, M.; Jäschke, A. Contact-Mediated Quenching for RNA Imaging in Bacteria with a Fluorophore-Binding Aptamer. Angew. Chemie - Int. Ed. 2013, 52 (50), 13401–13404.

(11) Murata, A.; Ichi Sato, S.; Kawazoe, Y.; Uesugi, M. Small-Molecule Fluorescent Probes for Specific RNA Targets. Chem. Commun. 2011, 47 (16), 4712–4714.

(12) Arora, A.; Sunbul, M.; Jäschke, A. Dual-Colour Imaging of RNAs Using Quencher- and Fluorophore-Binding Aptamers. Nucleic Acids Res. 2015, 43 (21).

(13) Braselmann, E.; Wierzba, A. J.; Polaski, J. T.; Chromiński, M.; Holmes, Z. E.; Hung, S. T.; Batan, D.; Wheeler, J. R.; Parker, R.; Jimenez, R.;, et al. A Multicolor Riboswitch-Based Platform for Imaging of RNA in Live Mammalian Cells. Nat. Chem. Biol. 2018, 14 (10), 964–971.

(14) Yang, X.; Pan, Z.; Choudhury, M. R.; Yuan, Z.; Anifowose, A.; Yu, B.; Wang, W.; Wang, B. Making Smart Drugs Smarter: The Importance of Linker Chemistry in Targeted Drug Delivery. Med. Res. Rev. 2020, 40 (6), 2682–2713.

(15) Borsari, C.; Trader, D. J.; Tait, A.; Costi, M. P. Designing Chimeric Molecules for Drug Discovery by Leveraging Chemical Biology. J. Med. Chem. 2020, 63 (5), 1908–1928.

(16) Leriche, G.; Chisholm, L.; Wagner, A. Cleavable Linkers in Chemical Biology. Bioorganic Med. Chem. 2012, 20 (2), 571–582.

(17) Saarbach, J.; Sabale, P. M.; Winssinger, N. Peptide Nucleic Acid (PNA) and Its Applications in Chemical Biology, Diagnostics, and Therapeutics. Curr. Opin. Chem. Biol. 2019, 52, 112–124.

(18) Brazil, R. Peptide Nucleic Acids Promise New Therapeutics and Gene Editing Tools. ACS Cent. Sci. 2023, 9 (1), 3–6.

(19) Nielsen, P. E.; Egholm, M.; Berg, R. H.; Buchardt, O. Sequence-Selective Recognition of DNA by Strand Displacement with a Thymine-Substituted Polyamide. Science (80-). 1991, 254 (5037), 1497–1500.

(20) Vilaivan, T. Fluorogenic PNA Probes. Beilstein J. Org. Chem. 2018, 14, 253–281.

(21) Koppelhus, U.; Nielsen, P. E. Cellular Delivery of Peptide Nucleic Acid (PNA). Adv. Drug Deliv. Rev. 2003, 55 (2), 267–280.

(22) Tsylents, U.; Siekierska, I.; Trylska, J. Peptide Nucleic Acid Conjugates and Their Antimicrobial Applications—a Mini-Review. Eur. Biophys. J. 2023, 52 (6–7), 533–544.

(23) Hawner, M.; Ducho, C. Cellular Targeting of Oligonucleotides by Conjugation with Small Molecules; 2020; Vol. 25.

(24) Równicki, M.; Wojciechowska, M.; Wierzba, A. J.; Czarnecki, J.; Bartosik, D.; Gryko, D.; Trylska, J. Vitamin B12 as a Carrier of Peptide Nucleic Acid (PNA) into Bacterial Cells. Sci. Rep. 2017, 7 (1), 7644.

(25) Równicki, M.; Dąbrowska, Z.; Wojciechowska, M.; Wierzba, A. J.; Maximova, K.; Gryko, D.; Trylska, J. Inhibition of Escherichia Coli Growth by Vitamin B 12 –Peptide Nucleic Acid Conjugates. ACS Omega 2019, 4 (1), 819–824.

(26) Giedyk, M.; Jackowska, A.; Równicki, M.; Kolanowska, M.; Trylska, J.; Gryko, D. Vitamin B 12 Transports Modified RNA into E. Coli and S. Typhimurium Cells. Chem. Commun. 2019, 55 (6), 763–766.

(27) Pieńko, T.; Czarnecki, J.; Równicki, M.; Wojciechowska, M.; Wierzba, A. J.; Gryko, D.; Bartosik, D.; Trylska, J. Vitamin B 12-Peptide Nucleic Acids Use the BtuB Receptor to Pass through the Escherichia Coli Outer Membrane. Biophys. J. 2021, 120 (4).

(28) Wierzba, A. J.; Wojciechowska, M.; Trylska, J.; Gryko, D. Vitamin B12 – Peptide Nucleic Acid Conjugates BT - Peptide Conjugation: Methods and Protocols; Hussein, W. M., Stephenson, R. J., Toth, I., Eds.; Springer US: New York, NY, 2021; pp 65–82.

(29) Johnson Jr, J. E.; Reyes, F. E.; Polaski, J. T.; Batey, R. T. B12 Cofactors Directly Stabilize an MRNA Regulatory Switch. Nature 2012, 492 (7427), 133–137.

(30) Merino, E. J.; Wilkinson, K. A.; Coughlan, J. L.; Weeks, K. M. RNA Structure Analysis at Single Nucleotide Resolution by Selective 2‘-Hydroxyl Acylation and Primer Extension (SHAPE). J. Am. Chem. Soc. 2005, 127 (12), 4223–4231.

(31) Wilkinson, K. A.; Merino, E. J.; Weeks, K. M. Selective 2′-Hydroxyl Acylation Analyzed by Primer Extension (SHAPE): Quantitative RNA Structure Analysis at Single Nucleotide Resolution. Nat. Protoc. 2006, 1 (3), 1610–1616.

(32) Jarmoskaite, I.; Alsadhan, I.; Vaidyanathan, P. P.; Herschlag, D. How to Measure and Evaluate Binding Affinities. Elife 2020, 9, 1–34.

(33) Unworth, H.; Raguz, S.; Edwards, H. J.; Higgins, C. F.; Yagüe, E. MRNA Escape from Stress Granule Sequestration Is Dictated by Localization to the Endoplasmic Reticulum. FASEB J. 2010, 24 (9), 3370–3380.

(34) Mollet, S.; Cougot, N.; Wilczynska, A.; Dautry, F.; Kress, M.; Bertrand, E.; Weil, D. Translationally Repressed MRNA Transiently Cycles through Stress Granules during Stress. Mol. Biol. Cell 2008, 19 (10), 4469–4479.

(35) Nelles, D. A.; Fang, M. Y.; O’Connell, M. R.; Xu, J. L.; Markmiller, S. J.; Doudna, J. A.; Yeo, G. W. Programmable RNA Tracking in Live Cells with CRISPR/Cas9. Cell 2016, 165 (2), 488–496.

(36) Zurla, C.; Lifland, A. W.; Santangelo, P. J. Characterizing MRNA Interactions with RNA Granules during Translation Initiation Inhibition. PLoS One 2011, 6 (5).

(37) Wu, J.; Zaccara, S.; Khuperkar, D.; Kim, H.; Tanenbaum, M. E.; Jaffrey, S. R. Live Imaging of MRNA Using RNA-Stabilized Fluorogenic Proteins. Nat. Methods 2019, 16 (9), 862–865.

(38) Chen, X.; Zhang, D.; Su, N.; Bao, B.; Xie, X.; Zuo, F.; Yang, L.; Wang, H.; Jiang, L.; Lin, Q.;, et al. Visualizing RNA Dynamics in Live Cells with Bright and Stable Fluorescent RNAs. Nat. Biotechnol. 2019, 37 (11), 1287–1293.

(39) Li, X.; Kim, H.; Litke, J. L.; Wu, J.; Jaffrey, S. R. Fluorophore-Promoted RNA Folding and Photostability Enables Imaging of Single Broccoli-Tagged MRNAs in Live Mammalian Cells. Angew. Chemie - Int. Ed. 2020, 59 (11), 4511–4518.

(40) Sarfraz, N.; Moscoso, E.; Oertel, T.; Lee, H. J.; Ranjit, S.; Braselmann, E. Visualizing Orthogonal RNAs Simultaneously in Live Mammalian Cells by Fluorescence Lifetime Imaging Microscopy (FLIM). Nat. Commun. 2023, 14 (1), 1–9.

(41) Tourrière, H.; Chebli, K.; Zekri, L.; Courselaud, B.; Blanchard, J. M.; Bertrand, E.; Tazi, J. The RasGAP-Associated Endoribonuclease G3BP Mediates Stress Granule Assembly. J. Cell Biol. 2023, 222 (11), e200212128072023new.

(42) Khong, A.; Matheny, T.; Jain, S.; Mitchell, S. F.; Wheeler, J. R.; Parker, R. The Stress Granule Transcriptome Reveals Principles of MRNA Accumulation in Stress Granules. Mol. Cell 2017, 68 (4), 808–820.e5.

(43) Matheny, T.; Van Treeck, B.; Huynh, T. N.; Parker, R. RNA Partitioning into Stress Granules Is Based on the Summation of Multiple Interactions. Rna 2021, 27 (2), 174–189.

(44) Liu, J. L.; Gall, J. G. U Bodies Are Cytoplasmic Structures That Contain Uridine-Rich Small Nuclear Ribonucleoproteins and Associate with P Bodies. Proc. Natl. Acad. Sci. U. S. A. 2007, 104 (28), 11655–11659.

(45) Tsalikis, J.; Tattoli, I.; Ling, A.; Sorbara, M. T.; Croitoru, D. O.; Philpott, D. J.; Girardin, S. E. Intracellular Bacterial Pathogens Trigger the Formation of U Small Nuclear RNA Bodies (U Bodies) through Metabolic Stress Induction. J. Biol. Chem. 2015, 290 (34), 20904–20918.

(46) McCloskey, A.; Taniguchi, I.; Shinmyozu, K.; Ohno, M. HnRNP C Tetramer Measures RNA Length to Classify RNA Polymerase II Transcripts for Export. Science (80-.). 2012, 335 (6076), 1643–1646.

(47) Singh, G.; Monga, V. Peptide Nucleic Acids: Recent Developments in the Synthesis and Backbone Modifications. Bioorg. Chem. 2023, 141, 106860.

(48) Sato, Y.; Miura, H.; Tanabe, T.; Okeke, C. U.; Kikuchi, A.; Nishizawa, S. Fluorescence Sensing of the Panhandle Structure of the Influenza A Virus RNA Promoter by Thiazole Orange Base Surrogate-Carrying Peptide Nucleic Acid Conjugated with Small Molecule. Anal. Chem. 2022, 94 (22), 7814–7822.

(49) Zhan, X.; Deng, L.; Chen, G. Mechanisms and Applications of Peptide Nucleic Acids Selectively Binding to Double-Stranded RNA. Biopolymers 2022, 113 (2).

(50) Lohse, J.; Dahl, O.; Nielsen, P. E. Double Duplex Invasion by Peptide Nucleic Acid: A General Principle for Sequence-Specific Targeting of Double-Stranded DNA. Proc. Natl. Acad. Sci. U. S. A. 1999, 96 (21), 11804–11808.

(51) Bentin, T.; Larsen, H. J.; Nielsen, P. E. Combined Triplex/Duplex Invasion of Double-Stranded DNA by “Tail-Clamp” Peptide Nucleic Acid. Biochemistry 2003, 42 (47), 13987–13995.

(52) Kiliszek, A.; Banaszak, K.; Dauter, Z.; Rypniewski, W. The First Crystal Structures of RNA-PNA Duplexes and a PNA-PNA Duplex Containing Mismatches - Toward Anti- Sense Therapy against TREDs. Nucleic Acids Res. 2015, 44 (4), 1937–1943.

(53) Falese, J. P.; Donlic, A.; Hargrove, A. E. Targeting RNA with Small Molecules: From Fundamental Principles towards the Clinic. Chem. Soc. Rev. 2021, 50 (4), 2224–2243.

(54) Childs-Disney, J. L.; Yang, X.; Gibaut, Q. M. R.; Tong, Y.; Batey, R. T.; Disney, M. D. Targeting RNA Structures with Small Molecules. Nat. Rev. Drug Discov. 2022, 21 (10), 736–762.

(55) Shao, Y.; Zhang, Q. C. Targeting RNA Structures in Diseases with Small Molecules. Essays Biochem. 2020, 64 (6), 955–966.

(56) Meyer, S. M.; Williams, C. C.; Akahori, Y.; Tanaka, T.; Aikawa, H.; Tong, Y.; Childs-Disney, J. L.; Disney, M. D. Small Molecule Recognition of Disease-Relevant RNA Structures. Chem. Soc. Rev. 2020, 49 (19), 7167–7199.

(57) Pradeep, S. P.; Malik, S.; Slack, F. J.; Bahal, R. Unlocking the Potential of Chemically Modified Peptide Nucleic Acids for RNA-Based Therapeutics. Rna 2023, 29 (4), 434–445.

(58) Edwards, A. L.; Garst, A. D.; Batey, R. T. Determining Structures of RNA Aptamers and Riboswitches by X-Ray Crystallography BT - Nucleic Acid and Peptide Aptamers: Methods and Protocols; Mayer, G., Ed.; Humana Press: Totowa, NJ, 2009; pp 135–163.

(59) Stoddard, C. D.; Gilbert, S. D.; Batey, R. T. Ligand-Dependent Folding of the Three-Way Junction in the Purine Riboswitch. Rna 2008, 14 (4), 675–684.

(60) Das, R.; Laederach, A.; Pearlman, S. M.; Herschlag, D.; Altman, R. B. SAFA: Semi-Automated Footprinting Analysis Software for High-Throughput Quantification of Nucleic Acid Footprinting Experiments. Rna 2005, 11 (3), 344–354.

