## Supplementary information for "Unveiling the Promise of Peptide Nucleic Acids as Functional Linkers for Riboglow RNA Imaging Platform"

### 1. Supporting Figures

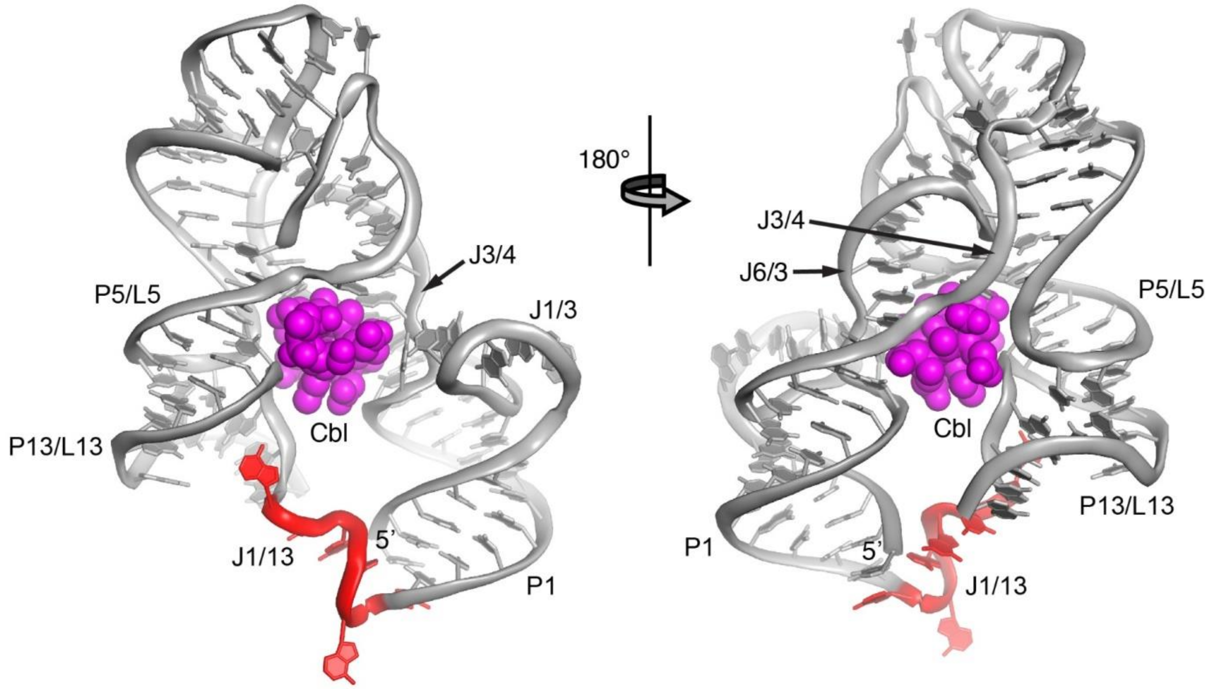

**Supporting Figure S1.** Crystal structure of wildtype *env*8 (*env*8-FL-3'*anti*PNA) bound to cobalamin (PDB: 4FRG).[1] Key structural regions denoted as P (paired), J (junction) and L (loop). J1/13 fragment is marked in red and cobalamin in magenta.

**PNA**

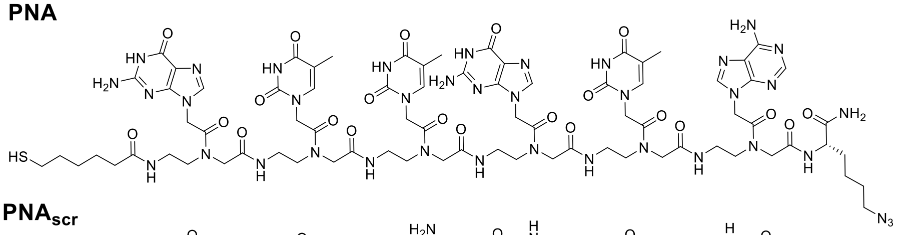

**PNA$_{scr}$**

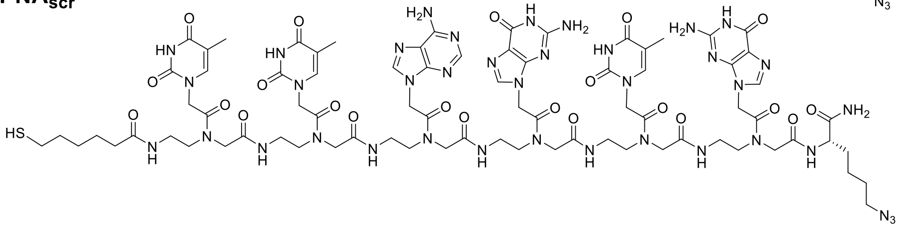

**PNA$_{truncated}$**

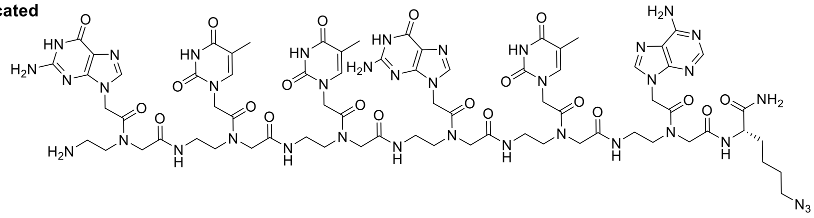

**Cbl-PNA-ATTO590**

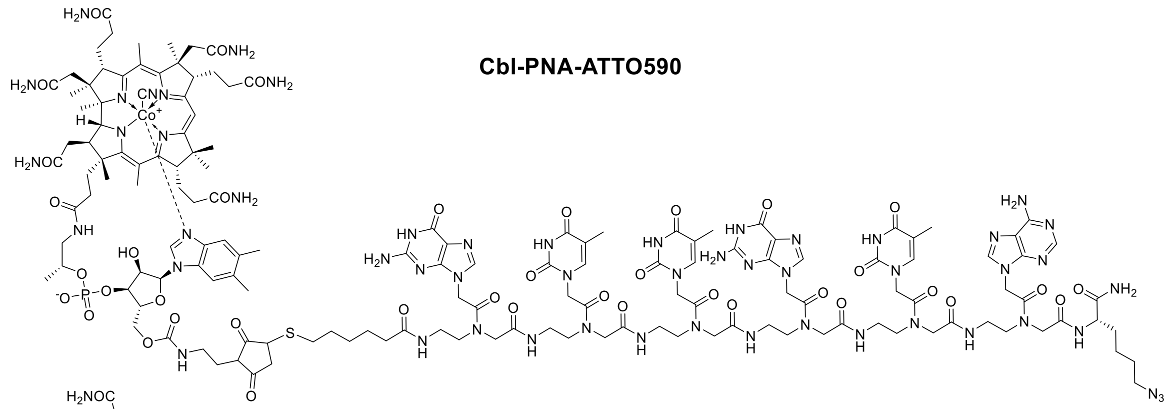

**Cbl-PNA$_{scr}$-ATTO590**

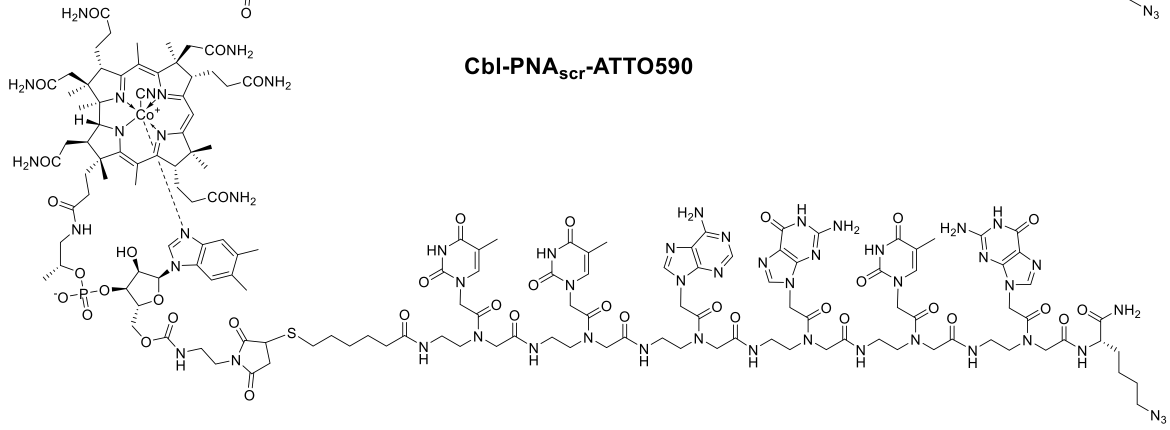

**Supporting Figure S2.** Chemical structures of PNA linkers and Cbl-PNA conjugates.

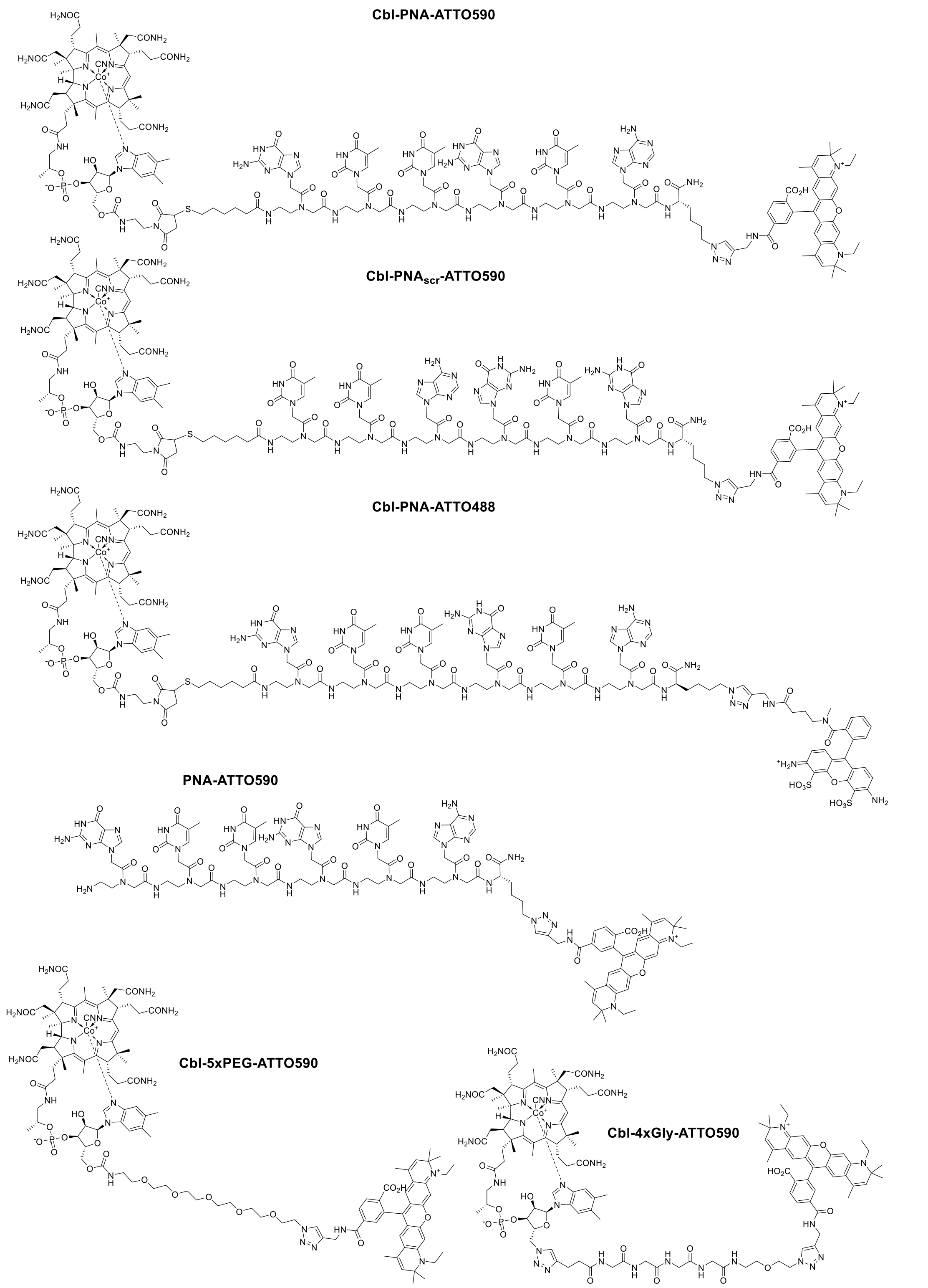

**Supporting Figure S3.** Chemical structures of Cbl-based probes and PNA-ATTO590 probe.

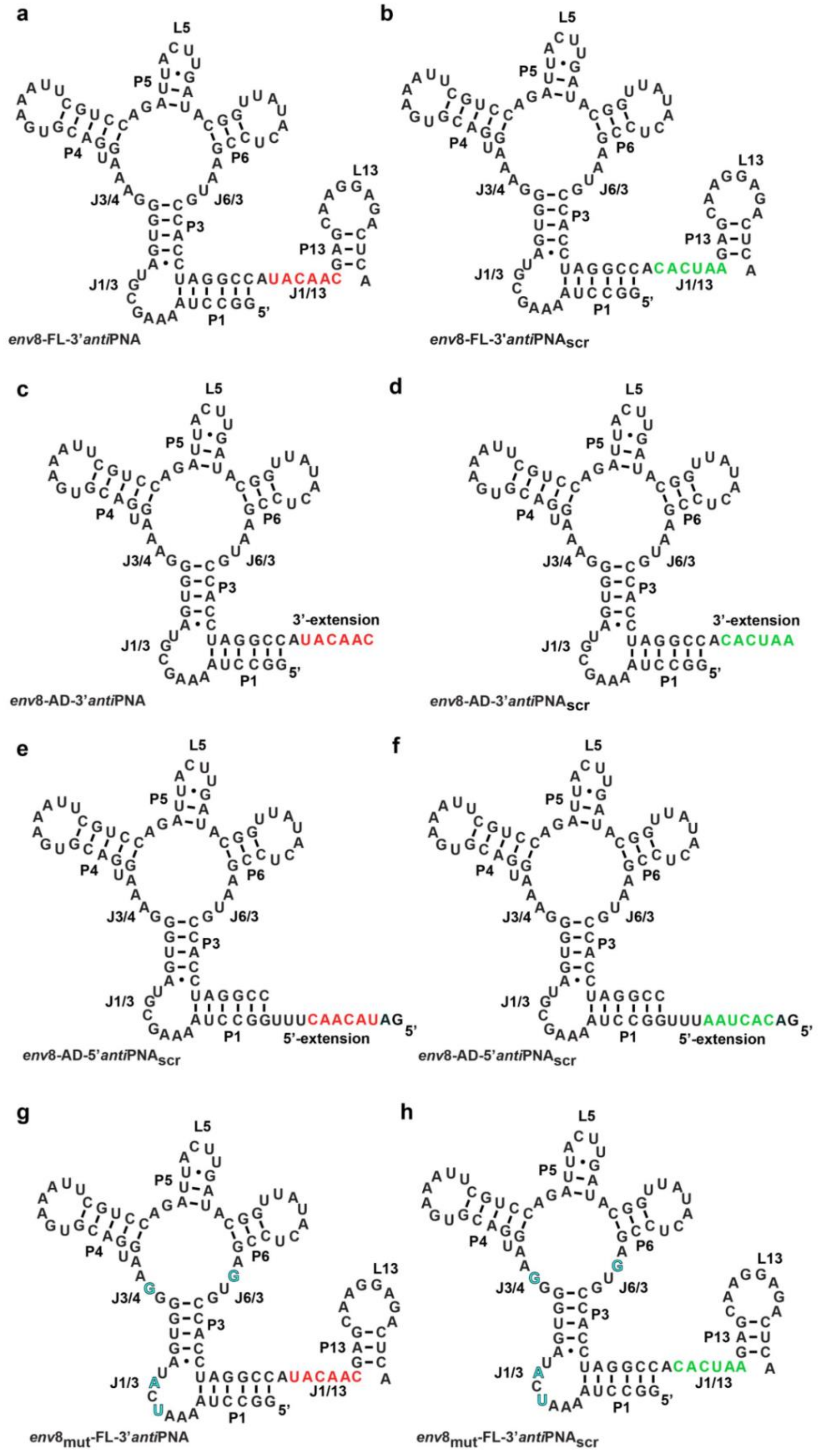

**Supporting Figure S4.** Secondary RNA structures used in the study. The *anti*PNA and *anti*$PNA_{scr}$ fragments are color-coded with red and green, respectively. Mutated nucleotides in **g** and **h** are colored blue.

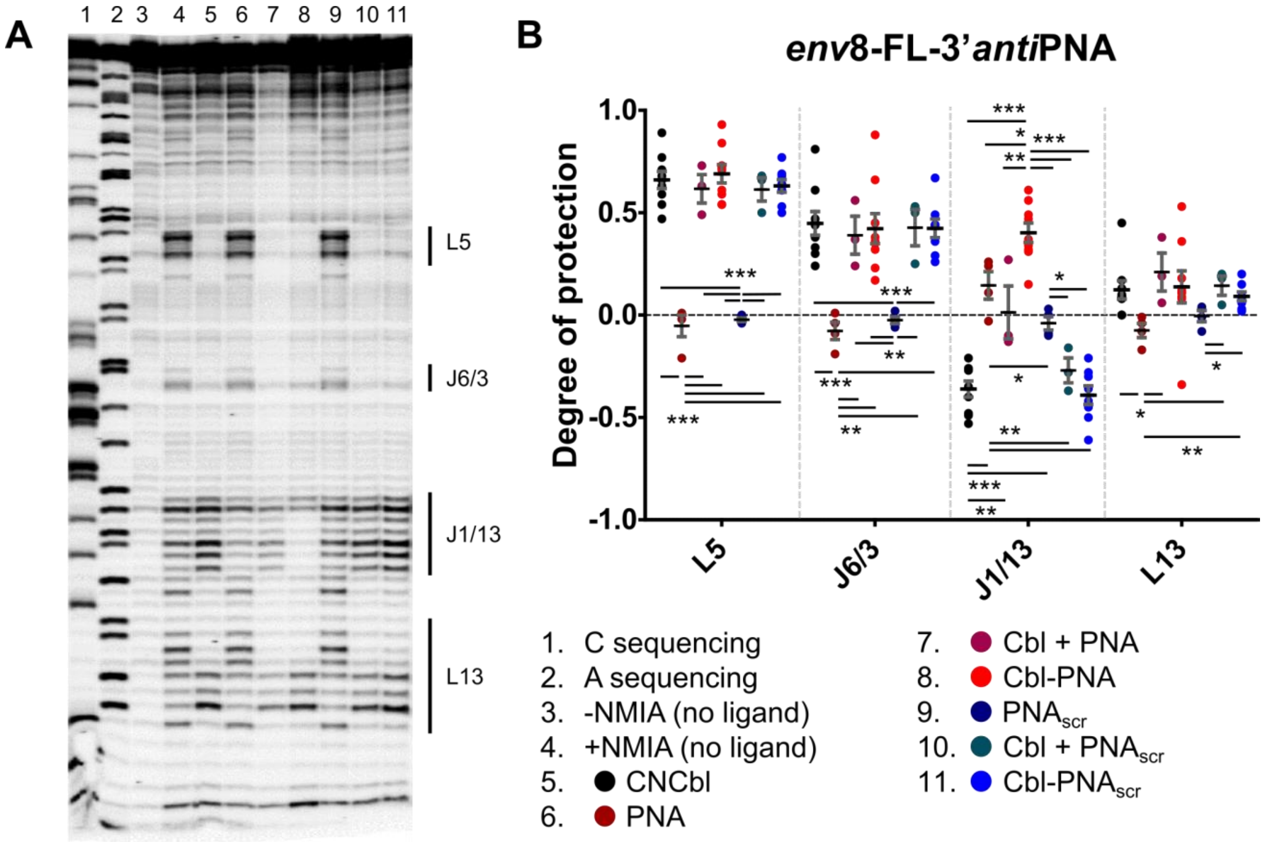

**Supporting Figure S5.** Extension of Figure 2. **A**. Full representative *env*8-FL-3'*anti*PNA SHAPE gel in the presence of several ligands. Lanes are loaded according to the key. Cbl-dependent SHAPE pattern changes in L5, J6/3, and L13 occur when Cbl is present in the ligand (lanes 5, 7, 8, 10, & 11 compared to lane 4). Sequence specific annealing of the PNA in the context of the Cbl-PNA ligand is seen by ligand dependent protections in J1/13 (lane 8 compared to lane 4). **B**. Quantifications of all marked SHAPE gel regions with the same ligands as A. Ligands are represented by colored dots according to the key. Degree of protection is in comparison to the +NMIA condition. Unpaired t-tests, *** $p < 0.001$, ** $p < 0.01$, * $p < 0.05$. Not significant p-values (>0.5) are not marked. n=3-9. Error bars represent SEM.

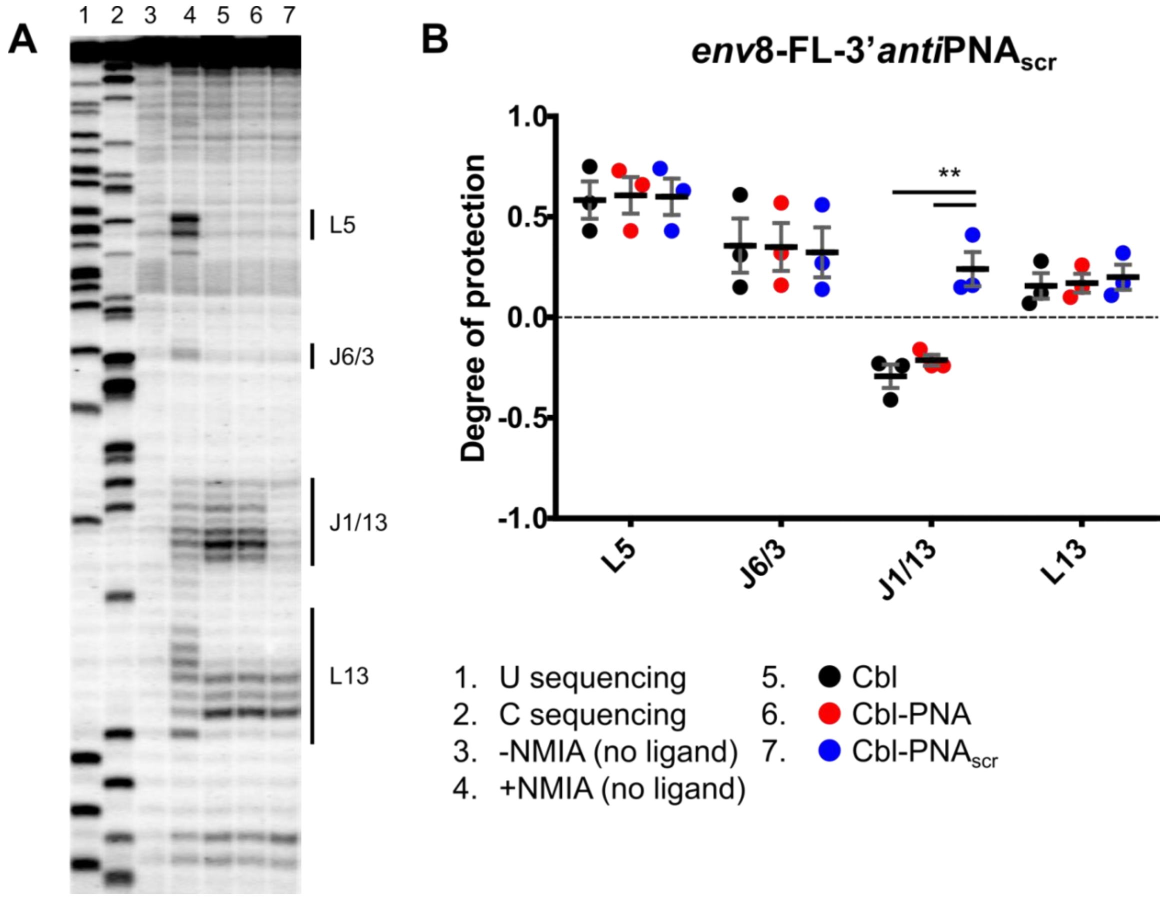

**Supporting Figure S6.** Extension of Figure 2. **A**. Full representative *env*8-FL-3'*anti*PNA$_{scr}$ SHAPE gel in the presence of several ligands. Lanes are loaded according to the key. Cbl-dependent SHAPE pattern changes in L5, J6/3, and L13 indicate Cbl binding (lanes 5, 6, & 7 compared to lane 4). Sequence specific annealing of the PNA$_{scr}$ is seen by ligand dependent protections in J1/13 (lane 7 compared to lane 4). **B**. Quantifications of all marked SHAPE gel regions with the same ligands as A. Ligands are represented by colored dots according to the key. Degree of protection is in comparison to the +NMIA condition. Unpaired t-tests, ** $p < 0.01$. Not significant p-values (>0.5) are not marked. n=3. Error bars represent SEM.

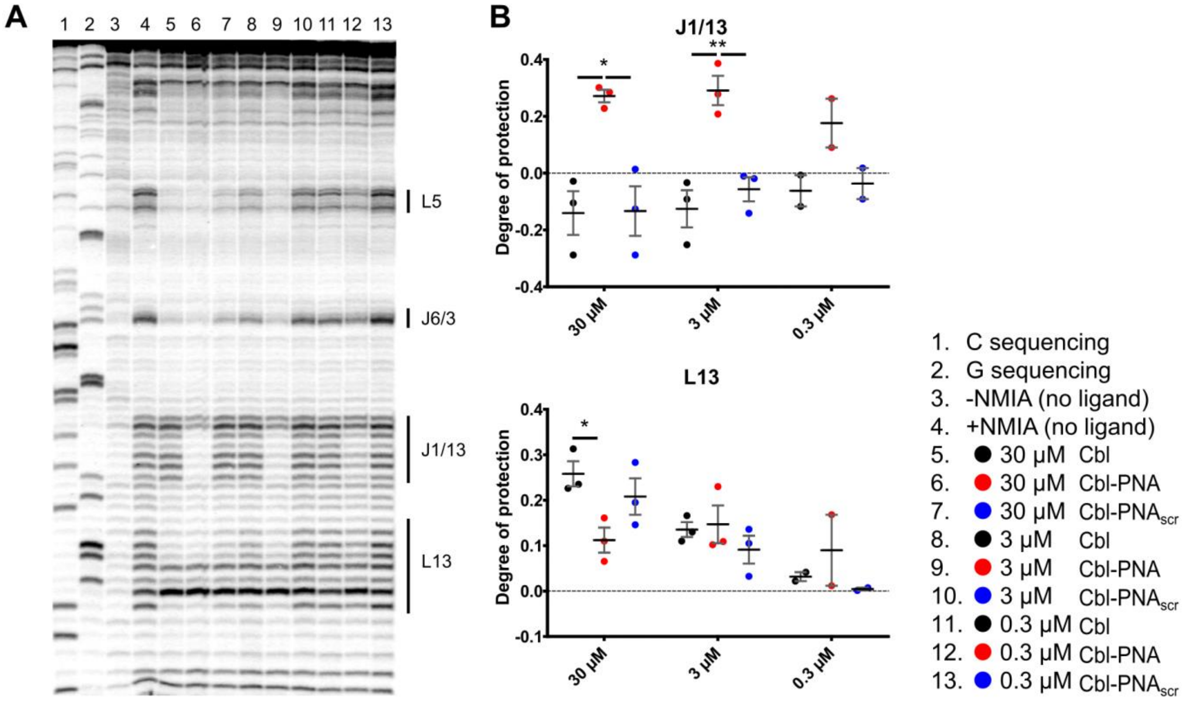

**Supporting Figure S7.** Extension of Figure 4. **A**. Full representative *env*8$_{mut}$-FL-3'*anti*PNA SHAPE gel in the presence of several ligands at different concentrations. Lanes are loaded according to the key. Cbl-dependent SHAPE pattern changes in L5, J6/3, and L13 indicate Cbl binding. Sequence specific annealing of the PNA is seen by ligand dependent protections in J1/13 (lanes 6, 9, & 12 compared to lane 4). **B**. Quantifications of marked SHAPE gel regions not quantified in Figure 6 with the same ligands and concentrations as A. Ligands are represented by colored dots according to the key. Degree of protection is in comparison to the +NMIA condition. Unpaired t-tests, ** $p < 0.01$, * $p < 0.05$. Not significant p-values (>0.5) are not marked. n=2-3. Error bars represent SEM.

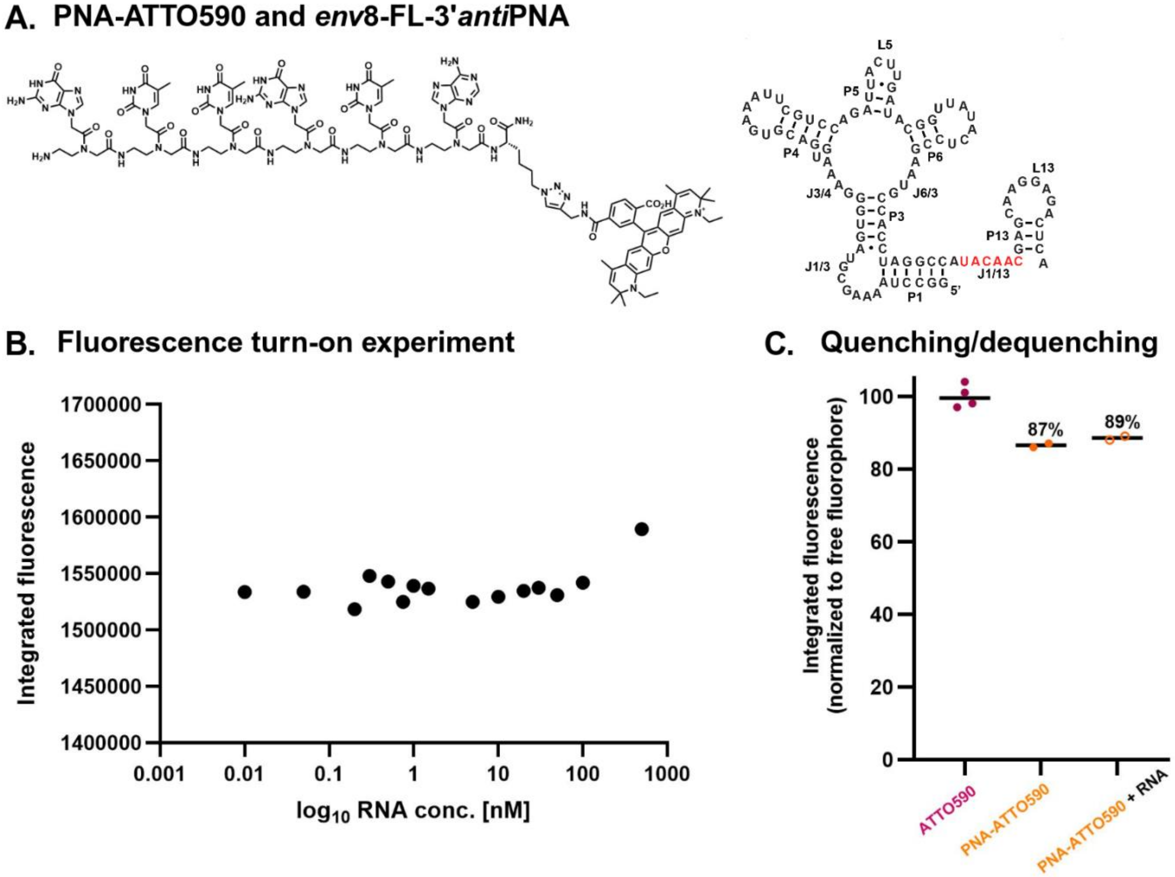

**Supporting Figure S8.** Control experiments indicating the relevance of cobalamin as a quencher in the Cbl-PNA-ATTO590 probe and PNA linker contribution towards quenching. **A.** Structures of PNA-ATTO590 probe and wild type *env*8. **B.** Titration of PNA-ATTO590 probe with *env*8-FL-3'*anti*PNA ($c_{PNA\text{-}ATTO590}$ = 1 nM). **C**. The levels of probe quenching and dequenching in the presence of RNA: *env*8-FL-3'*anti*PNA ($c_{probe}$ = 1 nM, $c_{RNA}$ = 500 nM).

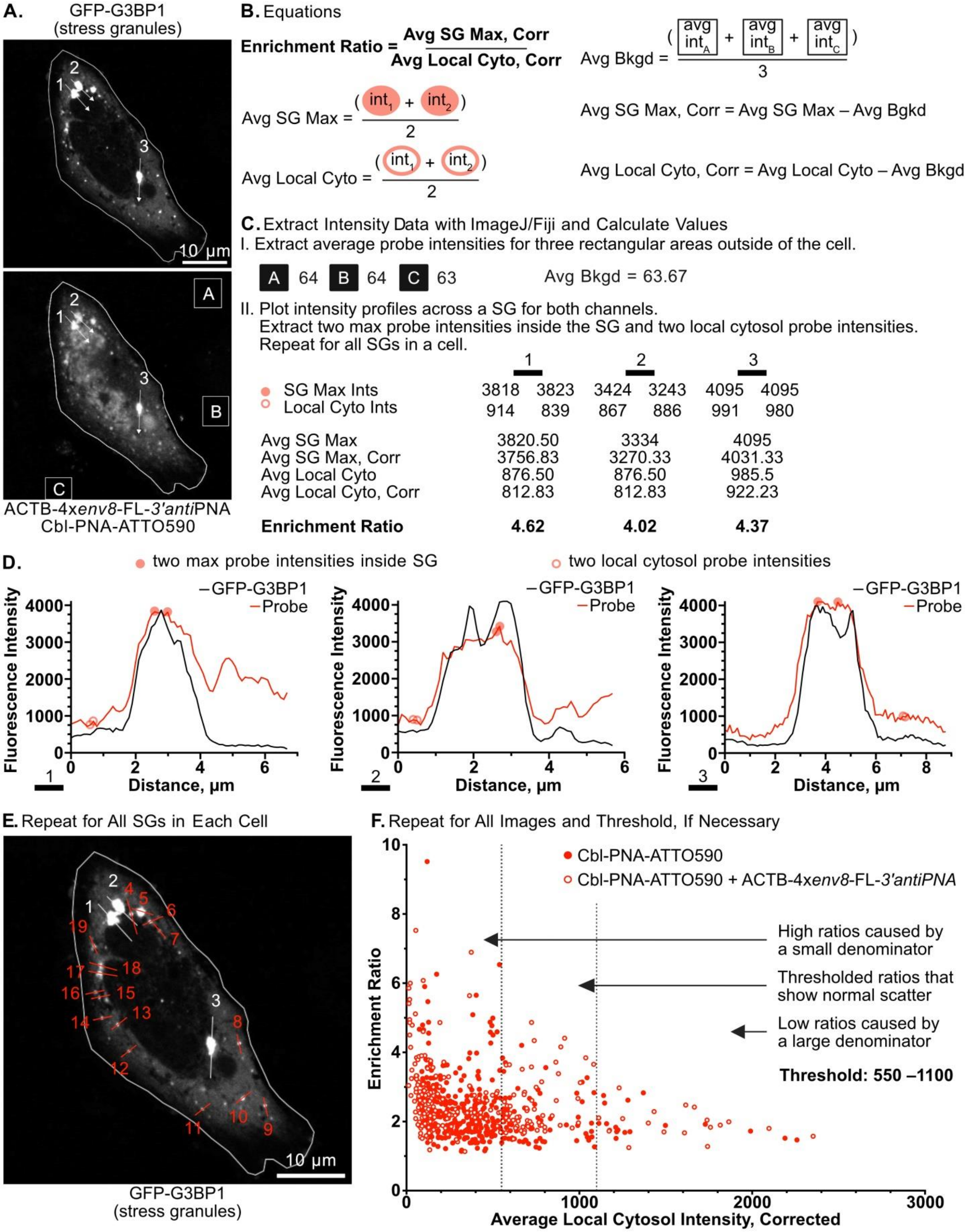

**Supporting Figure S9. A.** Representative images of a cell stably expressing GFP-G3BP1, co-transfected with NLS-TagBFP and ACTB-4x*env*8-FL-3'*anti*PNA, and beadloaded with 5 µM Cbl-PNA-ATTO590. Cell outline in white. Labeled white arrows 1, 2, and 3 showing example line scans across SGs. Labeled white squares showing example boxes outside of the cell. Scale bar = 10 µm. **B.** All equations necessary for calculation of enrichment ratio overlayed with shapes and colors to orient the reader to example intensity values in Panels C and D. **C.** (I) Values for the average probe intensities of the boxes outside the cell corresponding to the labeled boxes in Panel A and the calculation of average background probe intensity. (II) Values taken from the line scans in Panel D corresponding to the labeled arrows in Panel A and the calculation of average SG max probe intensities, background-corrected average SG max probe intensities, average local cytosol probe intensities, background-corrected average local cytosol probe intensities, and enrichment ratios. **D.** Line plots corresponding to the line scans of the labeled arrows in Panel A showing the fluorescence intensity in the GFP-G3BP1 (SG marker) channel and the probe channel versus distance in microns. Each plot shows two filled-in red circles for the two chosen max probe intensities inside the SG and two red outlined circles for the two chosen local cytosol probe intensities. **E.** GFP-G3BP1 channel for the same representative image as in Panel A with three white labeled arrows showing the three line scans with example values in Panels C and D and with more red labeled lines showing the remaining line scans that would be drawn to finish analysis of this cell. Scale bar = 10 µm. **F.** Dot plot of enrichment ratios versus background-corrected average local cytosol probe intensity. Each dot represents one SG. Filled in red circles are from cells without ACTB-4x*env*8-FL-3'*anti*PNA and red outlined circles are from cells co-transfected with ACTB-4x*env*8-FL-3'*anti*PNA. Dashed vertical lines at 550 and 1100 show the chosen thresholds for background-corrected average local cytosol probe intensity. These thresholds were chosen to avoid irregularities caused by particularly large or small denominators in the enrichment ratio equation.

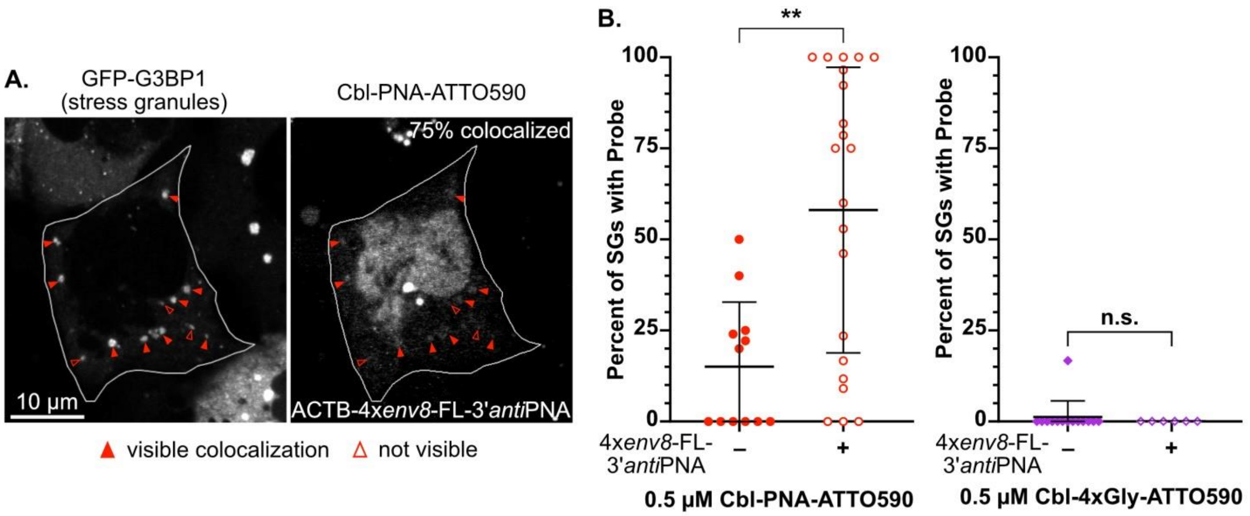

**Supporting Figure S10. A.** Representative image of a cell stably expressing GFP-G3BP1, co-transfected with NLS-TagBFP and ACTB-4x*env*8-FL-3'*anti*PNA, and beadloaded with 0.5 μM Cbl-PNA-ATTO590. Cell outline in white. Filled-in red triangles show SGs with visible colocalization while outlined red triangles show SGs without visible colocalization. Scale bar = 10 μm. **B.** Dot plots of percent of SGs with visible colocalization in cells co-transfected with NLS-TagBFP without or with ACTB-4x*env*8-FL-3'*anti*PNA and beadloaded with 0.5 μM Cbl-PNA-ATTO590. Each dot represents one cell. n= 6-21 cells across 2-4 imaging dishes for each condition as those quantified in Figure 5 Panel D. Bars show mean and standard deviation. p-values from Kolmogorov-Smirnov test (nonparametric cumulative distribution t-test). $p < 0.0001$ = ****, $p < 0.001$ = ***, $p < 0.01$ = **, $p < 0.05$ = *, $p \geq 0.05$ = n.s.

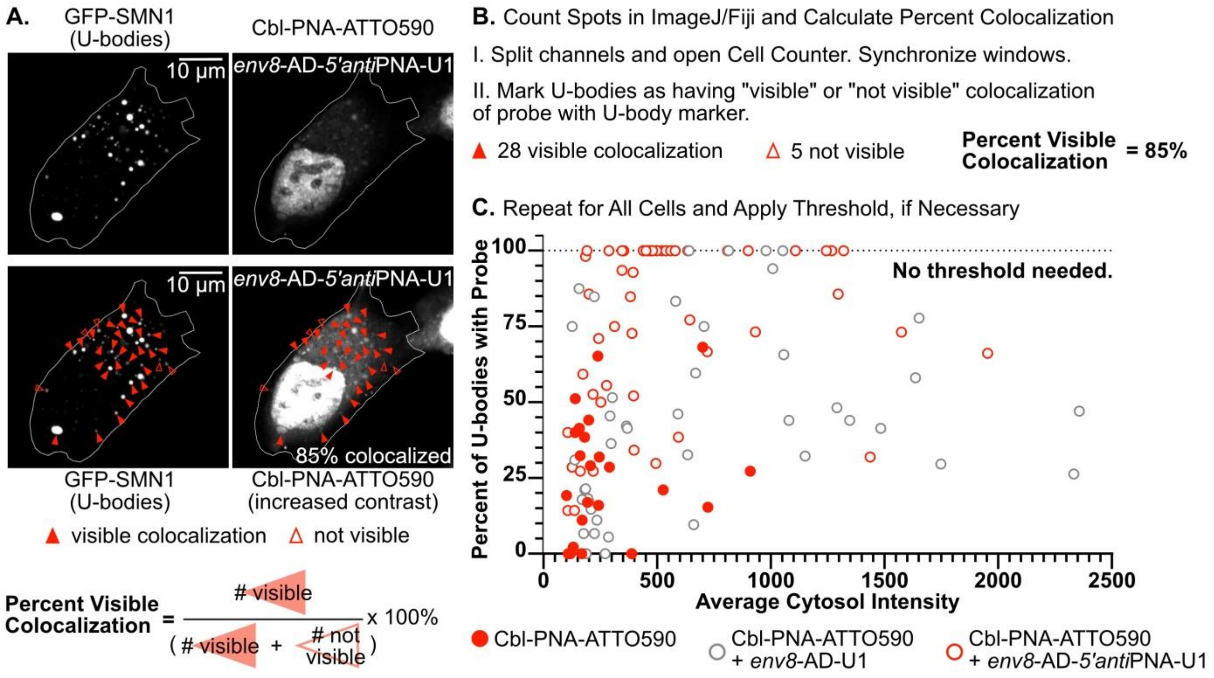

**Supporting Figure S11. A.** Representative images of a cell co-transfected with GFP-SMN1 and *env*8-AD-5'*anti*PNA-U1 and beadloaded with 50 µM Cbl-PNA-ATTO590. Cell outline in white. Filled-in red triangles show U-bodies with visible colocalization while outlined red triangles show U-bodies without visible colocalization. Scale bar = 10 µm. **B.** Equation to calculate percent visible, counts for the representative image in Panel A, and calculation of percent of U-bodies visible in the probe channel. **C.** Dot plot of percent visible versus average local cytosol probe intensity. Average local cytosol probe intensity was calculated by averaging the average probe intensities in three boxes inside the cytosol. Each dot represents one cell. Filled in red circles are from cells without Riboglow aptamer, grey outlined circles are from cells co-transfected with NLS-TagBFP and *env*8-AD-U1, and red outlined circles are from cells co-transfected with NLS-TagBFP and *env*8-AD-5'*anti*PNA-U1. There is no obvious structure to the data caused by average local cytosol intensity, so no thresholds were taken into account.

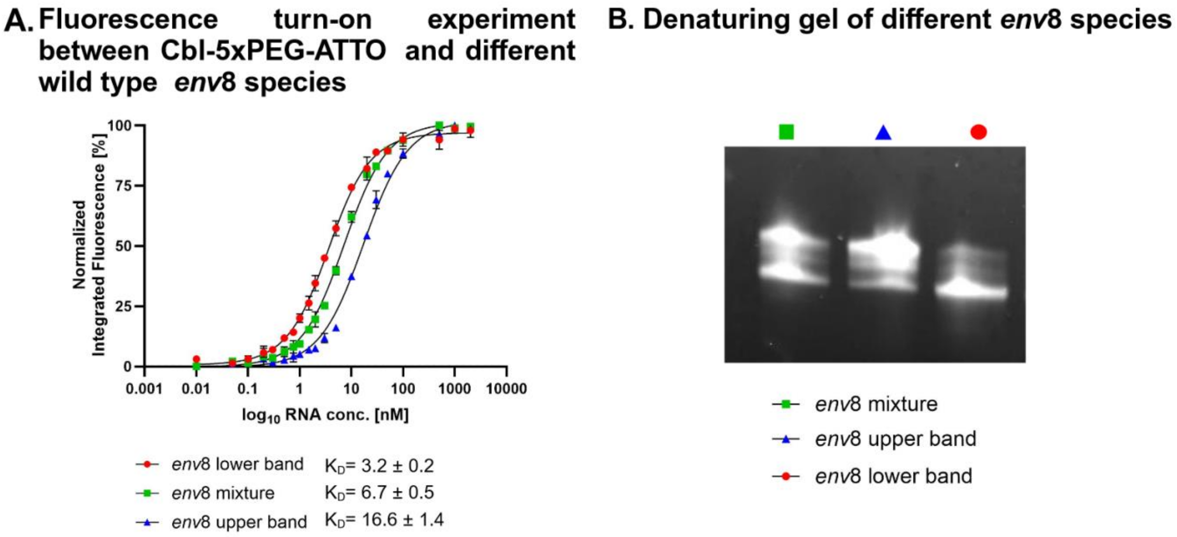

**Supporting Figure S12. A.** Fluorescence turn-on assay showing $K_D$ dependence on the purity of the wild type *env*8 ($c_{Cbl\text{-}5xPEG\text{-}ATTO590}$= 1 nM, n=4-8. Error bars represent SEM)**. B**. Denaturing gel showing successful separation on the products (8% acrylamide gel, see Methods). Note: Upper *env*8 transcription product is presumably a result of dsRNA self-templated addition by T7.[2]

### 2. Supporting Tables

**Table S1.** RNA sequences. The *anti*PNA and *anti*PNA$_{scr}$ fragments are color-coded with red and green, respectively.

| Name | Sequence |
|---|---|
| *env*8-FL-3'*anti*PNA | GGC CUA AAA GCG UAG UGG GAA AGU GAC GUG AAA UUC GUC CAG AUU ACU UGA UAC GGU UAU ACU CCG AAU GCC ACC UAG GCC AUA CAA CGA GCA AGG AGA CUC A |
| *env*8-FL-3'*anti*PNA_SHAPE | GGG CCU UCG GCC AAG GCC UAA AAG CGU AGU GGG AAA GUG ACG UGA AAU UCG UCC AGA UUA CUU GAU ACG GUU AUA CUC CGA AUG CCA CCU AGG CCA UAC AAC GAG CAA GGA GAC UCU CGA UCC GGU UCG CCG GAU CCA AAU CGG GCU UCG GUC CGG UUC |
| *env*8-FL-3'*anti*PNA$_{scr}$ | GGC CUA AAA GCG UAG UGG GAA AGU GAC GUG AAA UUC GUC CAG AUU ACU UGA UAC GGU UAU ACU CCG AAU GCC ACC UAG GCC ACA CUA AGA GCA AGG AGA CUC A |
| *env*8-FL-3'*anti*PNA$_{scr}$_SHAPE | GGG CCU UCG GGCC AAG GCC UAA AAG CGU AGU GGG AAA GUG ACG UGA AAU UCG UCC AGA UUA CUU GAU ACG GUU AUA CUC CGA AUG CCA CCU AGG CCA CAC UAA GAG CAA GGA GAC UCU CGA UCC GGU UCG CCG GAU CCA AAU CGG GCU UCG GUC CGG UUC |
| *env*8-AD-3'*anti*PNA | GGC CUA AAA GCG UAG UGG GAA AGU GAC GUG AAA UUC GUC CAG AUU ACU UGA UAC GGU UAU ACU CCG AAU GCC ACC UAG GCC AUA CAA C |
| *env*8-AD-3'*anti*PNA$_{scr}$ | GGC CUA AAA GCG UAG UGG GAA AGU GAC GUG AAA UUC GUC CAG AUU ACU UGA UAC GGU UAU ACU CCG AAU GCC ACC UAG GCC ACA CUA A |
| *env*8-AD-5'*anti*PNA | GAUA CAA C UUU GGC CUA AAA GCG UAG UGG GAA AGU GAC GUG AAA UUC GUC CAG AUU ACU UGA UAC GGU UAU ACU CCG AAU GCC ACC UAG GCC |
| *env*8-AD-5'*anti*PNA$_{scr}$ | GACA CUA A UUU GGC CUA AAA GCG UAG UGG GAA AGU GAC GUG AAA UUC GUC CAG AUU ACU UGA UAC GGU UAU ACU CCG AAU GCC ACC UAG GCC |
| *env*8$_{mut}$-FL-3'*anti*PNA | G GAG GC CUA AA UCA UAG UG GGG AAG UGA CGU GAA AUU CGU CCA GAU UAC UUG AUA CGG UUA UAC UCC GAG UGC CAC CUA GGC CAU ACA ACG AGC AAG GAG ACU CA |
| *env*8$_{mut}$-FL-3'*anti*PNA_SHAPE | GGC CUU CGG GCC AAG GCC UAA AUC AUA GUG GGG AAG UGA CGU GAA AUU CGU CCA GAU UAC UUG AUA CGG UUA UAC UCC GAG UGC CAC CUA GGC CAU ACA ACG AGC AAG GAG ACU CAU CGA UCC GGU UCG CCG GAU CCA AAU CGG GCU UCG GUC CGG UUC |
| *env*8$_{mut}$-FL-3'*anti*PNA$_{scr}$ | GGC CUA AA UCA UAG UG GGG AAG UGA CGU GAA AUU CGU CCA GAU UAC UUG AUA CGG UUA UAC UCC GAG UGC CAC CUA GGC CAC ACU AAG AGC AAG GAG ACU CA |
| *env*8$_{mut}$-FL-3'*anti*PNA$_{scr}$_SHAPE | GGC CUU CGG GCC AAG GCC UAA AUC AUA GUG GGG AAG UGA CGU GAA AUU CGU CCA GAU UAC UUG AUA CGG UUA UAC UCC GAG UGC CAC CUA GGC CAC ACU AAG AGC AAG GAG ACU CAU CGA UCC GGU UCG CCG GAU CCA AAU CGG GCU UCG GUC CGG UUC |

**Table S2.** Primer, ultramer and g-block sequences.

*Note: See next page for continuation.*

| RNA | primer/ultramer/gblock | Sequence |
|---|---|---|
| *env*8-FL-3'*anti*PNA | PCR'd plasmid | TAA TAC GAC TCA CTA TAG GGC CTA AAA GCG TAG TGG GAA AGT GAC GTG AAA TTC GTC CAG ATT ACT TGA TAC GGT TAT ACT CCG AAT GCC ACC TAG GCC ATA CAA CGA GCA AGG AGA CTC A |
| | Forward primer | GCG CGC GAA TTC TAA TAC GAC TCA CTA TAG GCC TAA AAG CGT AG |
| | Reverse primer | TGA GTC TCC TTG CTC GTT GTA TGG CCT AGG TGG CAT TCG GAG TAT A |
| *env*8-FL-3'*anti*$PNA_{scr}$ | Ultramer | TAA TAC GAC TCA CTA TAG GGC CTT CGG GCC AAG GCC TAA AAG CGT AGT GGG AAA GTG ACG TGA AAT TCG TCC AGA TTA CTT GAT ACG GTT ATA CTC CGA ATG CCA CCT AGG CCA CAC TAA GAG CAA GGA GAC TCT CGA TCC GGT TCG CCG GAT CCA AAT CGG GCT TCG GTC CGG TTC |
| | Forward primer | GCG CGC GAA TTC TAA TAC GAC TCA CTA TAG GCC TAA AAG CGT AG |
| | Reverse primer | TGA GTC TCC TTG CTC TTA GTG |
| *env*8-AD-3'*anti*PNA | Ultramer | TAA TAC GAC TCA CTA TAG GGC CTT CGG GCC AAG GCC TAA AAG CGT AGT GGG AAA GTG ACG TGA AAT TGG TCC AGA TTA CTT GAT ACG GTT ATA CTC CGA ATG CCA CCT AGG CCA TAC AAC TCG ATC CGG TTC GCC GGA TCC AAA TCG GGC TTC GGT CCG GTT C |
| | Forward primer | GCG CGC GAA TTC TAA TAC GAC TCA CTA TAG GCC TAA AAG CGT AG |
| | Reverse primer | GTT GTA TGG CCT AGG TGG C |
| *env*8-AD-3'*anti*$PNA_{scr}$ | Ultramer | TAA TAC GAC TCA CTA TAG GGC CTT CGG GCC AAG GCC TAA AAG CGT AGT GGG AAA GTG ACG TGA AAT TCG TCC AGA TAA CTT GAT ACG GTT ATA CTC CGA ATG CC ACCT AGG CCA CAC TAA TCG ATC CGG TTC GCC GGA TCC AAA TCG GGC TTC GGT CCG GTT C |
| | Forward primer | GCG CGC GAA TTC TAA TAC GAC TCA CTA TAG GCC TAA AAG CGT AG |
| | Reverse primer | TTA GTGT GGC CTA GGT GGC |
| *env*8-AD-5'*anti*PNA | Ultramer | TAA TAC GAC TCA CTA TAG GGC CTT CGG GCC AAT ACA ACT TTG GCC TAA AAG CGT AGT GGG AAA GTG ACG TGA AAT TCG TCC AGA TTA CTT GAT ACG GTT ATA CTC CGA ATG CCA CCT AGG CCT CGA TCC GGT TCG CCG GAT CCA AAT CGG GCT TCG GTC CGG TTC |
| | Forward primer | TAA TAC GAC TCA CTA TAG TAC AAC TTT GGC CTA AAA GCG TAG |
| | Reverse primer | GGC CTA GGT GGC ATT CGG |
| *env*8-AD-5'*anti*$PNA_{scr}$ | Ultramer | TAA TAC GAC TCA CTA TAG GGC CTT CGG GCC AAC ACT AAT TTG GCC TAA AAG CGT AGT GGG AAA GTG ACG TGA AAT TCG TCC AGA TTA CTT GAT ACG GTT ATA CTC CGA ATG CCA CCT AGG CCT CGA TCC GGT TCG CCG GAT CCA AAT CGG GCT TCG GTC CGG TTC |
| | Forward primer | TAA TAC GAC TCA CTA TAG CAC TAA TTT GGC CTA AAA GCG TAG TGG |
| | Reverse primer | GGC CTA GGT GGC ATT CGG |

**Table S2 continued.** Primer, ultramer and g-block sequences.

| RNA | primer/ultramer/gblock | Sequence |
|---|---|---|
| *env*8$_{mut}$-FL-3'*anti*PNA | Ultramer | GGA GCG CGC GAA TTC TAA TAC GAC TCA CTA TAG GCC TAA ATC ATA GTG GGG AAG TGA CGT GAA ATT CGT CCA GAT TAC TTG ATA CGG TTA TAC TCC GAG TGC CAC CTA GGC CAT ACA ACG AGC AAG GAG ACT C A |
| | Forward primer | GCG CGC GAA TTC TAA TAC GAC TCA CTA TAG GAG GCC TAA ATC ATA G |
| | Reverse primer | TGA GTC TCC TTG CTC GTT GTA TGG CCT AGG TGG CAC TCG GAG TAT A |
| *env*8$_{mut}$-FL-3'*anti*PNA$_{scr}$ | Ultramer | GGA GCG CGC GAA TTC TAA TAC GAC TCA CTA TAG GGC TAA ATC ATA GTG GGG AAG TGA CGT GAA ATT CGT CCA GAT TAC TTG ATA CGG TTA TAC TAA GAG TGC CAC CTA GGC CAC ACT AAG AGC AAG GAG ACT CA |
| | Forward primer | GCG CGC GAA TTC TAA TAC GAC TCA CTA TAG GAG GCC TAA ATC ATA G |
| | Reverse primer | TGA GTC TCC TTG CTC TTA GTG |
| *env*8-FL-3'*anti*PNA_SHAPE | gBlock | GGG CCT TCG GCC AAG GCC TAA AAG CGT AGT GGG AAA GTG ACG TGA AAT TCG TCC AGA TTA CTT GAT ACG GTT ATA CTC CGA ATG CCA CCT AGG CCA TAC AAC GAG CAA GGA GAC TCT CGA TCC GGT TCG CCG GAT CCA AAT CGG GCT TCG GTC CGG TTC |
| | Forward primer | GCG CGC GAA TTC TAA TAC GAC TCA CTA TAG |
| | Reverse primer | GAA CCG GAC CGA AGC CCG |
| *env*8-FL-3'*anti*PNA$_{scr}$_SHAPE | Ultramer | GGG CCT TCG GGC CAA GGC CTA AAA GCG TAG TGG GAA AGT GAC GTG AAA TTC GTC CAG ATT ACT TGA TAC GGT TAT ACT CCG AAT GCC ACC TAG GCC ACA CTA ATC GAT CCG GTT CGC CGG ATC CAA ATC GGG CTT CGG TCC GGT TC |
| | Forward primer | GCG CGC GAA TTC TAA TAC GAC TCA CTA TAG |
| | Reverse primer | GAA CCG GAC CGA AGC CCG |
| *env*8$_{mut}$-FL-3'*anti*PNA_SHAPE | Ultramer | GGA GCG CGC GAA TTC TAA TAC GAC TCA CTA TAG GCC TAA ATC ATA GTG GGG AAG TGA CGT GAA ATT CGT CCA GAT TAC TTG ATA CGG TTA TAC TCC GAG TGC CAC CTA GGC CAT ACA ACG AGC AAG GAG ACT C A |
| | Forward primer | GCG CGC GAA TTC TAA TAC GAC TCA CTA TAG GCC TTC GGG CCA AGG CCT AAA TCA TAG |
| | Reverse primer | GAA CCG GAC CGA AGC CCG ATT GGA TCC GGC GAA CCG GAT CGA TGG AGT CTC CTT GCT CGT |
| *env*8-AD-5'*anti*PNA-U1 | gBlock | AAG ATC TCA TAC TTA CCT GAT ACA ACT TTG GCC TAA AAG CGT AGT GGG AAA GTG ACG TGA AAT TCG TCC AGA TTA CTT GAT ACG GTT ATA CTC CGA ATG CCA CCT AGG CCG CAG GGG AGA TAC CAT GAT CA |
| | Forward primer | ATG CGA AGA TCT CAT ACT TAC CTG ATA C |
| | Reverse primer | ACT GTG ATC ATG GTA TCT CCC C |

**Table S3.** Reaction conditions for PCR and *in vitro* transcription.

| **PCR**[a] (Reaction volume = 100 μL) | |
|---|---|
| **Reagent** (stock concentration) | **Volume [μL]** |
| MilliQ $H_2O$ | 84 |
| Pfu Buffer[b] (10x) | 10 |
| dNTPs (10 mM) | 2 |
| Forward primer (100 μM) | 1 |
| Reverse primer (100 μM) | 1 |
| Template DNA (1 μM) | 1 |
| Pfu polymerase[e] | 1 |
| **Transcription**[c] (Reaction volume = 1 mL) | |
| **Reagent** (stock concentration) | **Volume [μL]** |
| MilliQ $H_2O$ | 590 |
| Transcription buffer[d] (10x) | 100 |
| $MgCl_2$ (2 M) | 12 |
| DTT (1 M) | 8 |
| rATP (100 mM) | 40 |
| rGTP (100 mM) | 40 |
| rCTP (100 mM) | 40 |
| rUTP(100 mM) | 40 |
| PCR template | 100 |
| Inorganic pyrophosphatase (IPP)[e] | 10 |
| T7 RNA polymerase[e] | 20 |

[a]The reaction is amplified in a PCR machine with the program: initial melt for 2 min at 95°C; 30 cycles of: 95°C for 30 s, 50°C for 30 s, and 72°C for 45 s; 10 min final extension at 72°C; hold at 4°C. [b]Pfu buffer: 200 mM Tris-HCl, 100 mM KCl, 100 mM $(NH_4)_2SO_4$, 20 mM $MgSO_4$, 1% Triton X-100, 1 mg/mL BSA. [c]The reaction is carried out at 37°C for 2.5 h. [d]Transcription buffer: 400 mM Tris pH 8.0, 100 mM DTT, 20 mM spermidine, 0.1% Triton X-100.[e]reagents made in house.

**Table S4.** Roadmap for fluorescence turn-on assay between Cbl-PNA-ATTO590 and *env*8-FL-3'*anti*PNA.

Each titration point contains: probe (1 nM), RNA (dissolved in water; concentration depends on the titration point), 1xRNA buffer, 0.01% nonidet P40 (surfactant; eliminates the issue of fluorophore sticking to eppendorf tubes and 384-well plates), 10% DMSO (ensures solubility of the probe)

The reactions are prepared in 1.5 mL eppendorfs. Two technical replicates are prepared in one eppendorf (total volume 2 x 60 μL = 120 μL). Every reaction contains equal volume of the master mix (see below) and different concentrations of the RNA. 55 μL of the final reaction mixture is pipetted twice into a Corning 384-well plate.

Master mix contains: probe, 10 x RNA buffer, DMSO, nonidet P40, $H_2O$ (enough to bring all the components to the expected concentrations, see below)

Notes:

- 10xRNA buffer contains: 1M KCl, 100 mM NaCl, 10 mM $MgCl_2$, 500 mM HEPES pH=8
- $c_{RNA\ stock}$= 40 μM (MilliQ water)
- $c_{probe\ stock}$= 335 μM (DMSO)

The table contains the volumes of reagents needed to perform the titration. For calculations, see below (color-coding is used for each reagent for easier text follow-up):

| Titration point | $c_{RNA}$ [nM] | $V_{RNA}$ [uL] (stock dilution) | $V_{H2O}$ [uL] | $V_{master\ mix}$ [uL] | $V_{total}$ [uL]= $V_{RNA}$ + $V_{H2O}$+ $V_{master\ mix}$ |
|---|---|---|---|---|---|
| 1 | 0 | 0 | 15 | 105 | 120 |
| 2 | 0.01 | 3 (1: 100000) | 12 | | |
| 3 | 0.05 | 15 (1: 100000) | 0 | | |
| 4 | 0.1 | 3 (1: 10000) | 12 | | |
| 5 | 0.2 | 6 (1: 10000) | 9 | | |
| 6 | 0.3 | 9 (1: 10000) | 6 | | |
| 7 | 0.5 | 15 (1: 10000) | 0 | | |
| 8 | 0.75 | 2.25 (1:1000) | 12.75 | | |
| 9 | 1 | 3 (1:1000) | 12 | | |
| 10 | 1.5 | 4.5 (1:1000) | 10.5 | | |
| 11 | 2 | 6 (1:1000) | 9 | | |
| 12 | 3 | 9 (1:1000) | 6 | | |
| 13 | 5 | 15 (1:1000) | 0 | | |
| 14 | 10 | 3(1:100) | 12 | | |
| 15 | 20 | 6 (1:100) | 9 | | |
| 16 | 50 | 15 (1:100) | 0 | | |

**How to calculate the volume of the RNA stock solution in each reaction ($V_{RNA}$ [uL])?**

Example for titration point 16:

$c_{RNA\ stock}$= 40 μM

$V_{reaction\ total}$ = 120 μL

$c_{RNA\ for\ point\ 16}$ = 50 nM

$V_{RNA}$ = ($c_{RNA}$/$c_{RNA\ stock}$)* $V_{reaction\ total}$ = [(50/1000)/40]*120 = 0.15 μL of RNA stock solution or 15 μL of RNA stock solution diluted at 1:100 ratio

Use the same approach to calculate the remaining titration points. See the table above for calculation results.

**How much of each RNA dilution is required?**

Start with diluting RNA stock to 1:10 and then proceed with serial dilutions. Prepare 30% more of each solution to have sufficient amount for the serial dilutions.

Below are summarized RNA volumes required for the titration:

$V_{RNA\ stock}$ (1:10) = 10 µL

$V_{RNA\ stock}$ (1:100, titration points 14-16) = 3+6+15+0.3*(3+6+15) = 32 µL

$V_{RNA\ stock}$ (1:1000, titration points 8-13) = (2.25+3+4.5+6+9+15)+0.3*(2.25+3+4.5+6+9+15)= 52 µL

$V_{RNA\ stock}$ (1:10000, titration points 4-7) = 3+6+9+15+0.3*(3+6+9+15)= 43 µL

$V_{RNA\ stock}$ (1:100000, titration points 2-3) = 3+15+0.3*(3+15)= 23 µL

**How to calculate the amount of water?**

The volume of the master mix remains constant for each titration point. However, since different volumes of RNA dilutions are used, additional water volume must be added to each eppendorf tube to bring the total volume in each well to the same level. Use the highest volume of the RNA dilution as a reference (in this case, 15 µL), and add this amount of water to the first eppendorf, which contains no RNA (the first titration point). For the remaining wells, subtract the volume of RNA from 15 µL (see the blue column in the table above for the results).

**How to prepare the master mix?**

Master mix contains: probe, 10x RNA buffer, DMSO, nonidet P40, $H_2O$

Begin by estimating the number of reactions. There are 16 titration points, but it's advisable to add extra volume in case of any repeats. Adding an extra 25% results in 20 reactions of volume 120 µL:

Probe:

$c_{probe\ stock}$= 335 µM

$C_{probe\ in\ reaction}$= 1 nM

$V_{probe\ stock\ for\ 20\ reactions}$= $[(c_{probe\ in\ reaction}/c_{probe\ stock})*\ V_{reaction\ total}]$*20 = [(1/(335*1000))*120]*20 = 0.0072 µL or 7.2 µL of probe stock solution diluted at 1:100 ratio

10x RNA buffer:

The reaction contains 1x RNA buffer, and the total reaction volume is 120 µL, so in each reaction there is 12 µL of 10x RNA buffer. Multiply 12 µL by the amount of the reactions:

$V_{10xRNAbuffer}$ = 12 µL * 20 = 240 µL

DMSO:

The final DMSO concentration in the reaction is 10%, so there is 12 µL of DMSO in each 120 µL reaction, analogous to the 10x RNA buffer.

$V_{DMSO}$= 12 µL * 20 = 240 µL

Nonidet P40:

Use 1% solution of nonidet. The final concentration of nonidet P40 in the reaction is 0.01%, so there is 1.2 μL of 1% nonidet P40 in each 120 μL reaction. For 20 reactions:

$V_{1\%nonidetP40}$= 1.2 μL * 20 = 24 μL

Water

This is the remaining amount of water that allows to obtain the expected volume and concentrations for all the ingredients.

$V_{H2O\ in\ the\ master\ mix\ for\ 20\ reactions}$= [$V_{reaction\ total}$ - ($V_{RNA}$+$V_{H2O}$) - $V_{10xRNAbuffer}$ - $V_{DMSO}$ - $V_{1\%nonidetP40}$ – ($V_{probe\ stock\ for\ 20\ reactions}$/20)*20= [120 - 15 – 12 – 12 – 1.2 – (7.2/20)]*20 = 1588.8 μL

Master mix = $V_{probe\ stock\ for\ 20\ reactions}$ + $V_{10xRNAbuffer}$ + $V_{DMSO}$ + $V_{1\%nonidetP40}$ + $V_{H2O\ in\ the\ master\ mix}$= 7.2 + 240 + 240 + 24 + 1588.8 = 2100 μL

The volume of master mix required for reach reaction:

$V_{master\ mix/reaction}$ = 2100/20 = 105 μL.

Other way of calculating $V_{master\ mix/reaction}$:

$V_{master\ mix/reaction}$ = $V_{reaction\ total}$ - ($V_{RNA}$+$V_{H2O}$) = 120 – 15 = 105 μL

**Preparing the reactions and plating**

- Prepare 16 numbered eppendorfs of 1.5 mL volume.
- Thaw RNA stock on ice; prepare RNA dilutions in $H_2O$; heat at 90°C for 3 minutes before incubating on ice for at least 10 min
- Prepare master mix during RNA incubation
- Pipet water volumes according to the amounts from the table above ($V_{H2O}$)
- Pipet master mix into each eppendorf (105 μL per reaction); for consistency vortex master mix every 3 eppendorfs
- Spin down all eppendorfs
- Spin down RNA dilutions and pipet into eppendorfs according to the amounts from the table above ($V_{RNA}$)
- Vortex and spin down all eppendorfs
- Plate 55 μL of each reaction on 384 well plate (each reaction is pipetted twice; vortex each eppendorf before plating)
- Pipet one buffer well (55 μL) for background subtracting purpose (buffer for background subtraction (1 mL): 100 μL of 10x RNA buffer, 100 μL DMSO, 10 μL of 1% nonidet P40, 790 μL $H_2O$)
- Incubate at room temperature in the dark for 1 hour before reading
- Read the plate using suitable program (see Methods)

**Note: The above roadmap was appropriately adjusted depending on the titration range required for each probe. Every RNA stock solution concentration used in the study was 40 μM and every probe stock concentration was 335 μM.**

**Table S5**. Imaging settings for live cell experiments. Laser wattages were collected with a PM100A Optical Power Meter (Thorlabs) and a S130C 400 nm – 1100 nm sensor (Thorlabs) on the 5 mW setting. Reported wattages were collected just before the objective in the light path (at an open slot on the objective wheel) and are reported as the max wattage in the manually scanned area.

| Experiment | Imaging Settings |
|---|---|
| **SG assay**<br>RNA: ACTB-4x*env8*-FL-3'*anti*PNA<br>Probe: 5 µM Cbl-PNA-ATTO590 or 5 µM Cbl-5xPEG-ATTO590 | 405 nm (NLS-TagBFP): 0.7 (0.065 mW) – 5% (0.090 mW), 90 gain<br>488 nm (GFP-G3BP1): 0.5 (0.029 mW) – 1.5% (0.035 mW), 40 gain<br>561 nm (Cbl-PNA-ATTO590, Cbl-5xPEG-ATTO590): 20% (0.344 mW), 40 gain<br>2X averaging, 12.1 µs dwell time<br>Nyquist sampling (1.2 AU) defined by 561-nm laser<br>Single plane image |
| **SG assay**<br>RNA: ACTB-4x*env8*-FL-3'*anti*PNA<br>Probe: 5 µM Cbl-4xGly-ATTO590 | 405 nm (NucBlue): 2% (0.072 mW), 90 gain<br>488 nm (GFP-G3BP1): 0.5 (0.029 mW) – 2% (0.041 mW), 40 gain<br>561 nm (Cbl-4xGly-ATTO590): 20% (0.344 mW), 40 gain<br>640 nm (PB-HaloTag-ACTB-0x + HaloTag-JF669 ligand): 0.5 (0.017 mW) – 5% (0.125 mW), 90 gain<br>2X averaging, 12.1 µs dwell time<br>Nyquist sampling (1.2 AU) defined by 561-nm laser<br>Single plane image |
| **SG assay**<br>RNA: (1/8)NORAD-3x*env8*-FL-3'*anti*PNA, ACTB-4x*env8*-FL-3'*anti*PNA, or (1/2)NORAD-4x*env8*-FL-3'*anti*PNA<br>Probe: 5 µM Cbl-PNA-ATTO590 | 405 nm (NLS-TagBFP): 1 (0.067 mW) – 10% (0.119 mW), 90 gain<br>488 nm (GFP-G3BP1): 0.3 (0.027 mW) – 1.2% (0.034 mW), 40 gain<br>561 nm (Cbl-PNA-ATTO590): 8% (0.144 mW), 40 gain<br>2X averaging, 12.1 µs dwell time<br>Nyquist sampling (1.2 AU) defined by 561-nm laser<br>Single plane image |
| **SG assay**<br>RNA: (1/2)NORAD-4x*env8*-FL-3'*anti*PNA<br>Probe: 5 µM Cbl-PNA-ATTO488 | 405 nm (NLS-TagBFP): 2 (0.072 mW) – 10% (0.119 mW), 90 gain<br>488 nm (Cbl-PNA-ATTO488): 20% (0.198 mW), 30 gain<br>640 nm (HaloTag-G3BP1 + HaloTag-JF669 ligand): 2 (0.055 mW) – 6% (0.148 mW), 90 gain<br>2X averaging, 12.1 µs dwell time<br>Nyquist sampling (1.2 AU) defined by 488-nm laser<br>Single plane image |
| **SG assay**<br>RNA: ACTB-4x*env8*-FL-3'*anti*PNA<br>Probe: 0.5 µM Cbl-PNA-ATTO590 or 0.5 µM Cbl-4xGly-ATTO590 | 405 nm (NLS-TagBFP): 0.7 (0.065 mW) – 10% (0.119 mW), 90 gain<br>488 nm (GFP-G3BP1): 0.3 (0.027 mW) – 6% (0.075 mW), 40 gain<br>561 nm (Cbl-PNA-ATTO590, Cbl-4xGly-ATTO590): 30% (0.512 mW), 40 gain<br>2X averaging, 12.1 µs dwell time<br>Nyquist sampling (1.2 AU) defined by 561-nm laser<br>Single plane image |
| **U-body assay**<br>RNA: *env8*-AD-U1 or *env8*-AD-5'*anti*PNA-U1<br>Probe: 50 µM Cbl-PNA-ATTO590 | 405 nm (NucBlue): 2 (0.072 mW) – 5%, 90 gain<br>488 nm (EGFP-SMN1): 0.2 (0.023 mW) – 1% (0.031 mW), 20 – 40 gain<br>561 nm (Cbl-PNA-ATTO590): 5% (0.093 mW), 40 gain<br>2X averaging, 12.1 µs dwell time<br>Nyquist sampling (1.2 AU) defined by 561-nm laser<br>Single plane image |

### 3. Supporting information: probe synthesis and characterization

#### 3.1. General Information

Commercially available reagents and solvents were used as received. PNA monomers were obtained from PNA Bio and Rink amide resin from Chem-Impex. ATTO590 propargylamide and ATTO488 propargylamide were obtained from Millipore Sigma. As supplied ATTO 590 consists of a mixture of two isomers with similar spectral properties (para and meta isomer). For simplicity only one isomer is presented on the schemes. The scale of the reaction with ATTO dyes did not provide sufficient amount of the products for NMR analyses, thus the HPLC and HR MS analyses were performed to characterize and confirm the purity of the probes. The synthesis of Cbl-PNA conjugates does not require purification of the respective PNA linker; therefore, crude material was used to synthesize these conjugates and only small amount of each PNA linker was purified via semipreparative HPLC and characterized via HPLC and HR MS. $^{1}H$ and $^{13}C$ NMR spectra were recorded at room temperature on a Bruker 400 MHz spectrometer with the residual solvent peak used as an internal standard. Data are reported as follows: chemical shift, peak multiplicity (s = singlet, d = doublet, t = triplet, q = quartet, m = multiplet), coupling constants (Hz), and number of protons. High-resolution ESI mass spectra were recorded on Waters Synapt G2 HDMS qTOF. All reactions and product purities were monitored using RP-HPLC techniques. Semipreparative chromatography was performed using LiChroprep RP-18 (40–63 mm) with HPLC grade water and MeCN as eluents. HPLC analytical measurement conditions: column, Kromasil 100-5-C18, 250 mm, 4.6 mm; detection, UV/Vis; pressure, 10 MPa; temperature, 22ºC, flow 1 mL/min or phenomenex Jupiter 5u C18 300A, 250 mm, 4.6 mm; detection, UV/Vis; pressure, 10 MPa; temperature, 22ºC, flow 1 mL/min. HPLC semipreparative measurement conditions: Kromasil 100-5-C18, 250 mm, 4.6 mm; detection, UV/Vis; pressure, 20 MPa; temperature, 22ºC, flow 3 mL/min. Abbreviations: AcOEt – ethyl acetate; Bhoc – benzhydryloxycarbonyl protecting group; CDT – 1,1′-Carbonyldi-(1,2,4-triazole); DIPEA – *N*,*N*-Diisopropylethylamine; DMAP – 4-Dimethylaminopyridine; $Et_2O$ – diethyl ether; Fmoc – fluorenylmethoxycarbonyl protecting group; HATU – 1-[Bis(dimethylamino)methylene]-1H-1,2,3-triazolo[4,5-b]pyridinium 3-oxide hexafluorophosphate; HOAt – 1-Hydroxy-7-azabenzotriazole; MeCN – acetonitrile; MeOH – methanol; NMM – 4-Methylmorpholine; NMP – *N*-Methyl-2-pyrrolidone; RP HPLC – Reverse-phase high-performance liquid chromatography; TBTA – Tris[(1-benzyl-1H-1,2,3-triazol-4 yl)methyl]amine; TFA – trifluoroacetic acid.

#### 3.2. Synthesis of PNA linkers

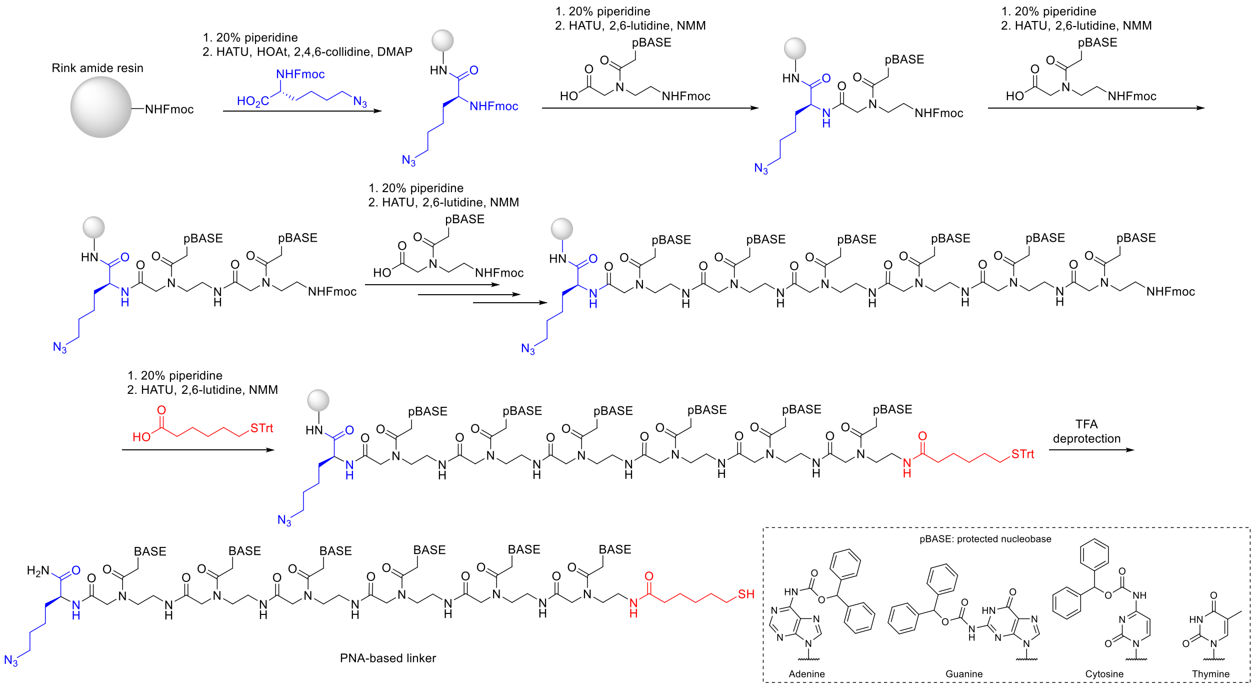

**Scheme S1.** Schematic representation of PNA linker synthesis using Fmoc chemistry.

PNA linkers were synthesized according to previously published protocol[3] with modifications. PNA linkers were synthesized manually by Fmoc chemistry on 0.05 mmol scale using 2.5 molar excess of the Fmoc/Bhoc protected PNA monomers, 3.0 molar excess of Fmoc-Lys($N_3$)-OH, 3.0 molar excess of 6-(tritylthio)hexanoic acid and Rink amide resin (loading 0.4 mmol/g). Fmoc deprotection of the resin was carried with 20% piperidine in DMF (1x for 5 min and 1x for 15 min). Fmoc-Lys($N_3$)-OH was activated with the mixture of HATU (3.0 equiv), HOAt (3.0 equiv.), 2,4,6-collidine (6.0 equiv.) and catalytic amount of DMAP in DMF/NMP (1:1; v/v). Coupling of Fmoc-Lys($N_3$)-OH was carried for 2 h. Deprotection of the Fmoc form the lysine was carried with 20% piperidine in DMF (1x for 5 and 1x for 15 min). PNA momomers were activated with the mixture of HATU (2.3 equiv), NMM (2.5 equiv) and 2,6-lutidine (3.75 equiv) in DMF/NMP (1:1; v/v). Each coupling of the PNA monomer was performed twice for 40 min. Fmoc deprotection of each PNA monomer was performed with 20% of piperidine in DMF (2 × 2 min). 6-(tritylthio)hexanoic acid was activated with HATU (2.8 equiv), NMM (3.0 equiv) and 2,6-lutidine (4.5 equiv) in DMF/NMP (1:1; v/v). Coupling of 6-(tritylthio)hexanoic acid was performed twice for 30 min. The resin was washed consecutively with DCM (2x), DMF (5x) and DCM (3x) after each Fmoc deprotection and with DMF (3x) and DCM (3x) consecutively after each coupling step. To deprotect and cleave the product the resin was treated with TFA/ triisopropylsilane/m-cresol mixture (95:2.5:2.5 v/v/v) for 1 h. The reaction mixture was then precipitated with ice-cold $Et_2O$, centrifuged and dried.

*Note: The crude PNA linkers had sufficient purity to be successfully used in the next step (Cbl-PNA conjugate synthesis). Purification of a small sample (approx. 5 mg) for characterization purposes was performed using semipreparative RP-HPLC (see HPLC method below).*

HPLC purification method for PNA linkers (λ= 254 nm):

| Time [min] | Water + 0.02%TFA [%] | Acetonitrile [%] |
|---|---|---|
| Initial | 90 | 10 |
| 5 | 90 | 10 |
| 24 | 50 | 50 |
| 25 | 90 | 10 |
| 27 | 90 | 10 |

HPLC analytical method for PNA linkers (λ= 254 nm):

| Time [min] | Water + 0.02%TFA [%] | Acetonitrile [%] |
|---|---|---|
| Initial | 90 | 10 |
| 10 | 30 | 70 |
| 13 | 30 | 70 |
| 14 | 90 | 10 |
| 16 | 90 | 10 |

#### 3.3. Characterization of PNA linkers

PNA (sequence G-T-T-G-T-A)

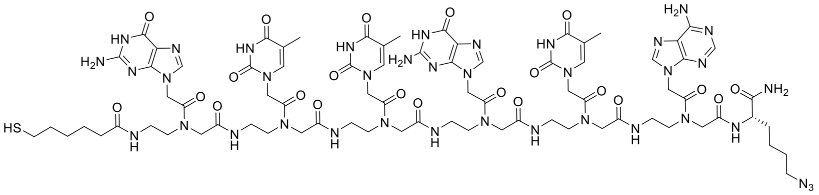

The compound was obtained as a white solid; crude mass = 60 mg. HRMS (ESI) m/z $[M + 2H]^{2+}$ calculated for $C_{78}H_{106}N_{38}O_{22}S$, 979.4027; found, 979.4064. $t_R$ (RP-HPLC): 8.19 min.

HPLC chromatogram (λ= 254 nm)

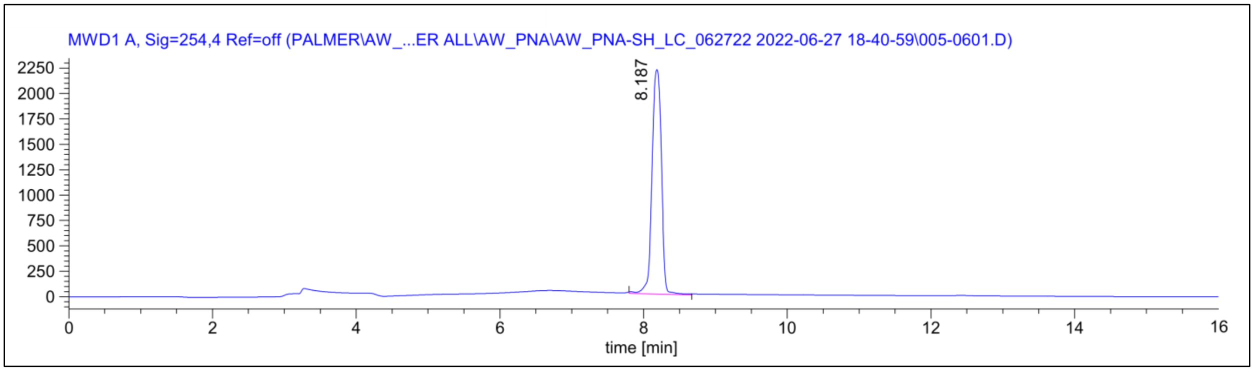

ESI MS spectrum

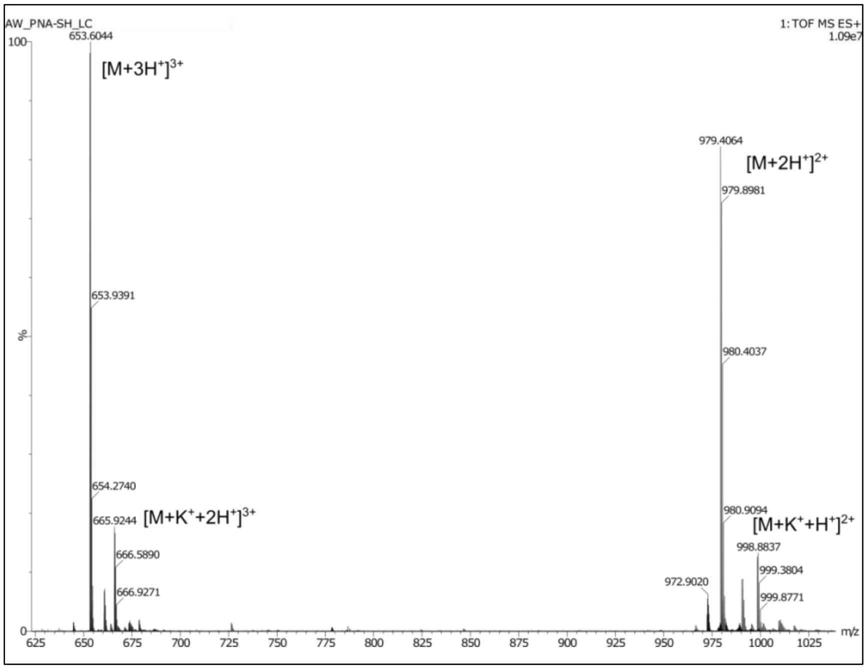

PNA$_{scr}$ (sequence T-T-A-G-T-G)

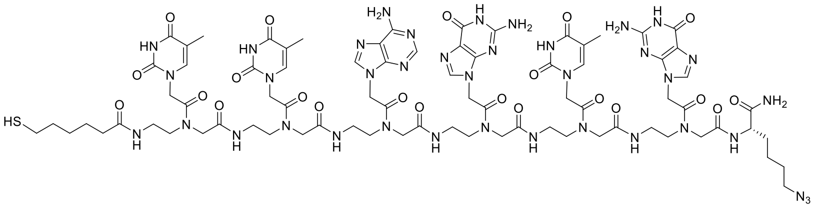

The compound was obtained as a white solid crude mass = 73 mg. HRMS (ESI) m/z $[M + 2H]^{2+}$ calculated for $C_{78}H_{106}N_{38}O_{22}S$, 979.4027; found, 979.4006. $t_R$ (RP-HPLC): 9.91 min.

HPLC chromatogram

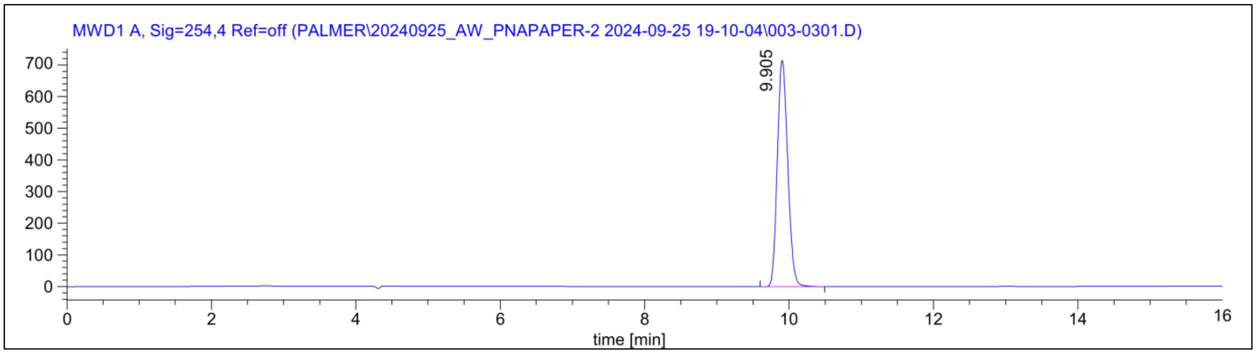

ESI MS spectrum

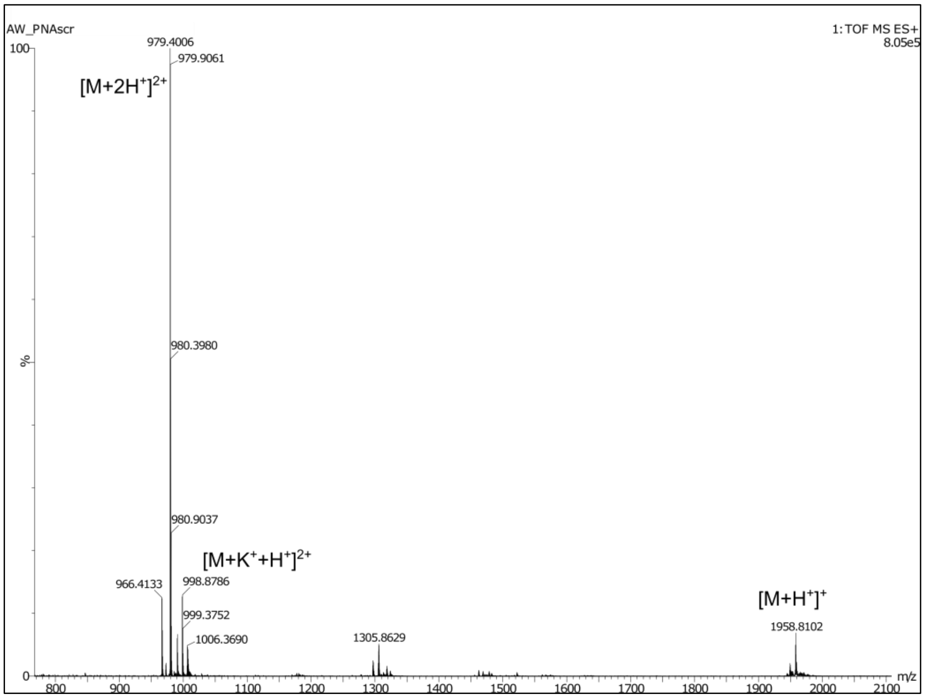

PNA$_{truncated}$ (sequence G-T-T-G-T-A)

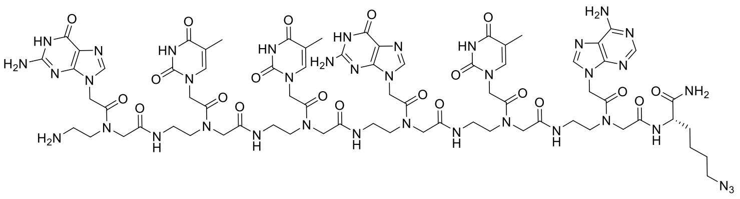

The compound was obtained as a white solid; crude mass = 65 mg. HRMS (ESI) m/z $[M + H]^+$ calculated for $C_{72}H_{95}N_{38}O_{21}$, 914.3801; found, 914.3806. $t_R$ (RP-HPLC): 8.02 min.

HPLC chromatogram

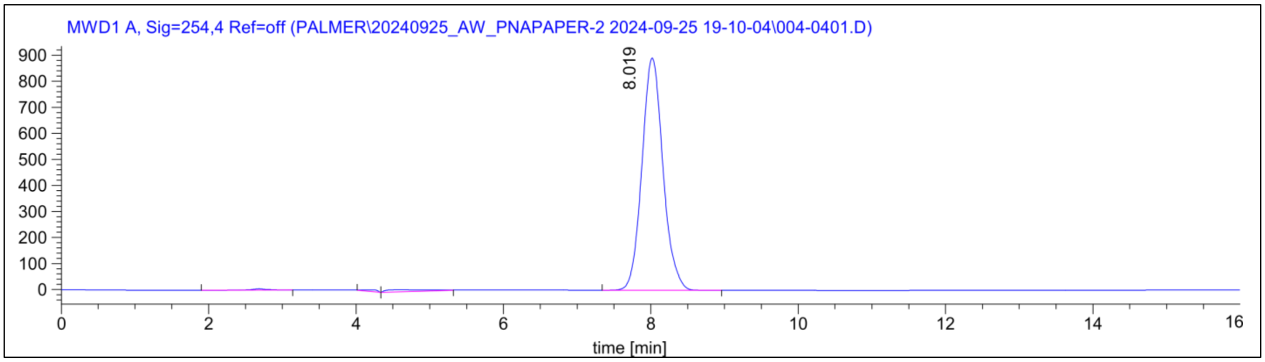

ESI MS spectrum

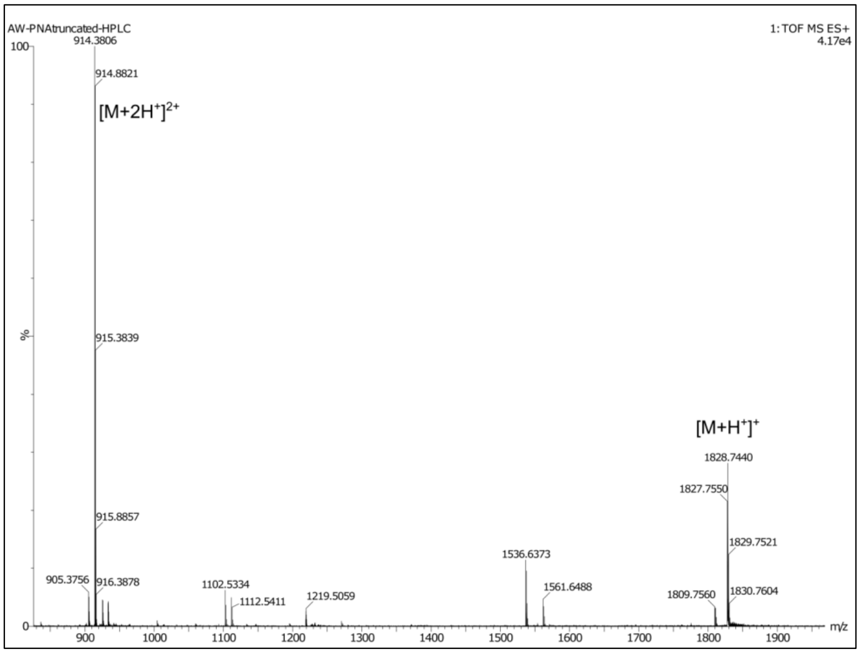

#### 3.4. Synthesis of cobalamin derivative with maleimide functionality

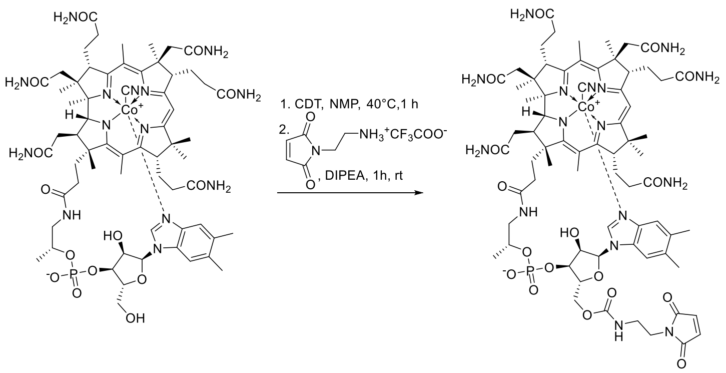

**Scheme S2.** Synthesis of maleimide-functionalized cobalamin derivative.

Cobalamin (70 mg, 0.05 mmol, 1 equiv.) was dissolved in dry *N*-Methyl-2-pyrrolidone (NMP, 3.0 mL) at 40 °C under an argon atmosphere. To a stirring solution under argon solid CDT (21 mg, 0.125 mmol, 2.5 equiv.) was added. When full consumption of the substrate (monitored by the RP HPLC) was observed (approx. 1.0 h), heating bath was removed and 1-(2-aminoethyl)-1*H*-pyrrole-2,5-dione as TFA salt (12.7 mg, 0.05 mmol, 1 equiv.) was added in one portion along with DIPEA (13 µL, 0.075 mmol, 1.5 equiv.). The resulting solution was stirred for 1h at room temperature. Subsequently, the reaction mixture was poured into AcOEt (10 mL), and centrifuged. The solid residue was redissolved in MeOH and precipitated with $Et_2O$ (10 mL), and centrifuged. After drying, the remaining solid was dissolved in water and purified by RP column chromatography with a mixture of MeCN and $H_2O$ as eluents (gradually from 10% to 25% v/v).

HPLC analytical method for Cbl-maleimide ($\lambda$= 254 and 361 nm):

| Time [min] | Water + 0.02%TFA [%] | Acetonitrile [%] |
|---|---|---|
| Initial | 90 | 10 |
| 10 | 30 | 70 |
| 13 | 30 | 70 |
| 14 | 90 | 10 |
| 16 | 90 | 10 |

#### 3.5. Characterization of cobalamin derivative with maleimide functionality

Cbl-maleimide

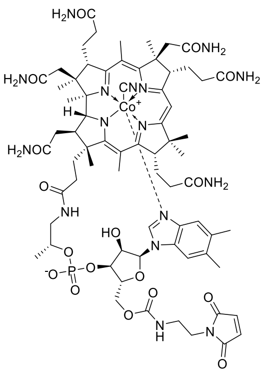

The compound was obtained as a red solid; yield: 40%. HRMS (ESI) m/z $[M + 2H]^{2+}$ calculated for $C_{70}H_{96}CoN_{16}O_{17}P$, 761.3099; found, 761.3108. $t_R$ (RP-HPLC): 7.44 min.

$^1$H NMR (400 MHz, $CD_3OD$) δ 7.27 (s, 1H), 7.16 (s, 1H), 6.66 (s, 1H), 6.58 (s, 1H), 6.28 (d, *J* = 2.9 Hz, 1H), 6.06 (s, 1H), 4.83 – 4.71 (m, 2H), 4.58 (s, 1H), 4.53 (d, *J* = 9.0 Hz, 1H), 4.39 – 4.32 (m, 1H), 4.25 – 4.18 (m, 2H), 4.14 (d, *J* = 11.4 Hz, 1H), 4.09 (dd, *J* = 12.3, 2.3 Hz, 1H), 3.70 – 3.59 (m, 3H), 3.57 – 3.50 (m, 1H), 3.41 – 3.32 (m, 1H), 3.28 – 3.18 (m, 1H), 2.96 – 2.83 (m, 2H), 2.72 – 1.82 (m, 17H) 2.59 (s, 6H), 2.38 (d, *J* = 2.7 Hz, 2H), 2.28 (d, *J* = 8.2 Hz, 6H), 2.07 (d, *J* = 13.8 Hz, 2H), 1.89 (s, 3H), 1.48 (s, 3H), 1.37 (d, 3H), 1.39 (d, 3H), 1.32 – 1.23 (m, 1H) 1.24 (d, *J* = 6.3 Hz, 3H), 1.19 (s, 3H), 1.14 – 1.07 (m, 1H), 0.48 (s, 3H). $^{13}$C NMR (100 MHz, $CD_3OD$) δ 181.56, 180.15, 177.59, 177.33, 176.59, 175.55, 175.50, 175.31, 174.62, 174.04, 172.53, 167.19, 166.90, 158.68, 143.40, 138.25, 135.58, 135.34, 133.93, 131.35, 127.47, 117.87, 112.60, 108.66, 105.25, 95.60, 88.21, 86.40, 81.53, 76.36, 75.01, 73.40, 70.50, 66.90, 64.37, 63.67, 60.34, 60.31, 57.65, 56.91, 55.05, 52.63, 52.58, 52.50, 46.58, 43.89, 43.03, 40.26, 40.07, 38.68, 36.21, 35.01, 33.15, 32.40, 29.54, 27.43, 20.90, 20.57, 20.33, 20.18, 20.14, 19.90, 17.49, 17.14, 16.39, 16.13, 15. 44.

$^{1}H$ and $^{13}C$ NMR spectra recrded in $CD_3OD$ (for clarity only selected signals were integrated in $^{1}H$ NMR spectra)

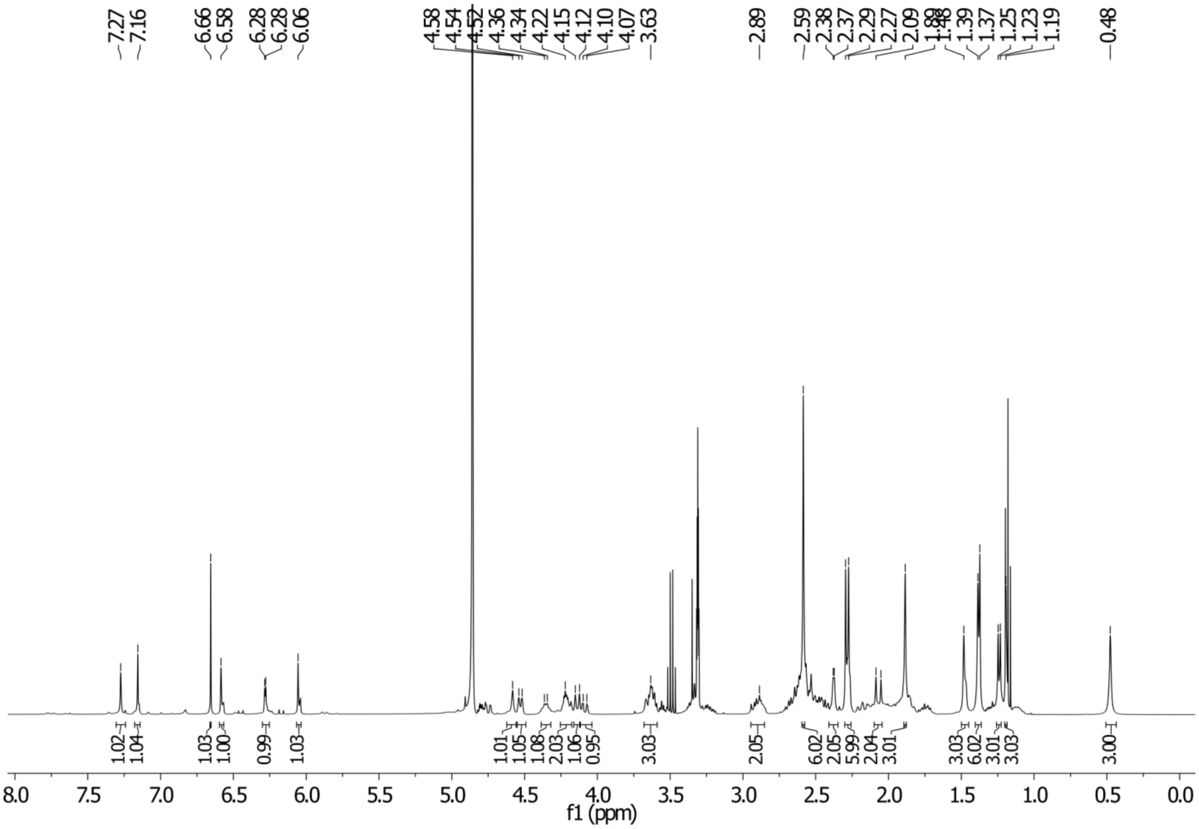

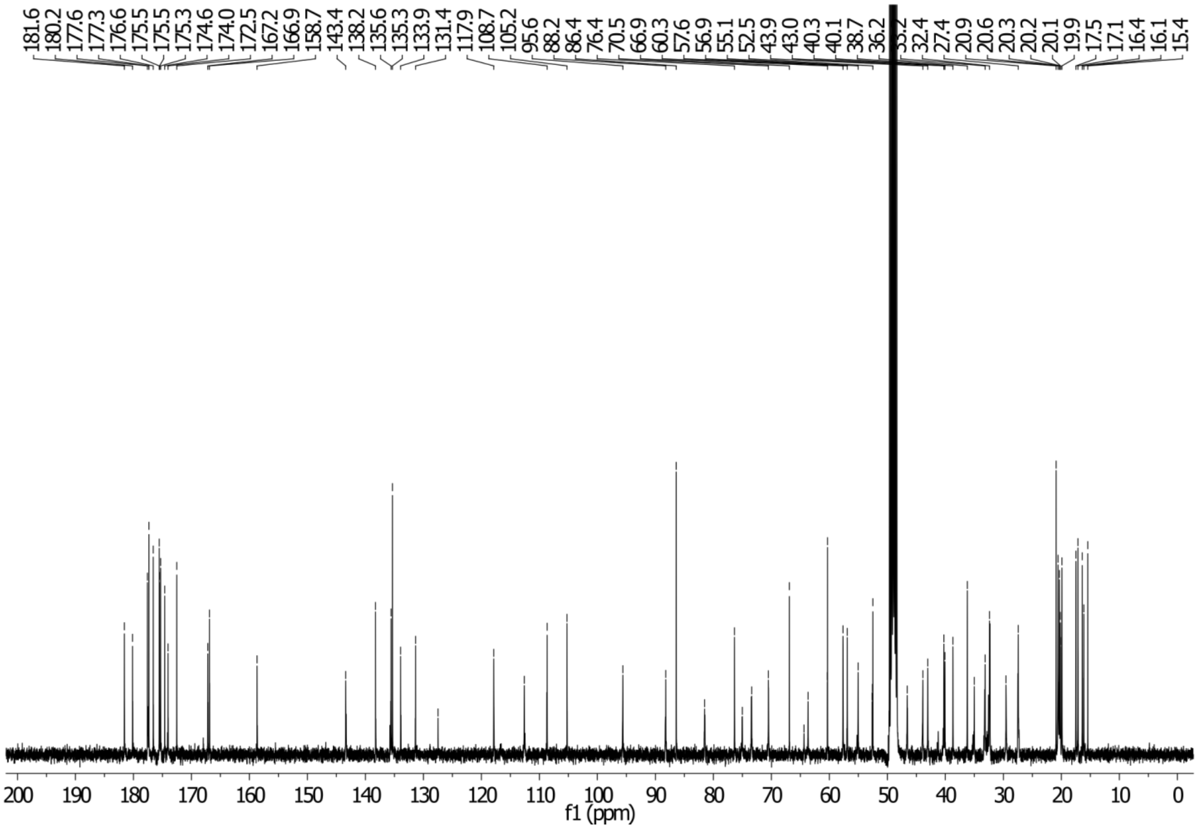

HPLC chromatogram

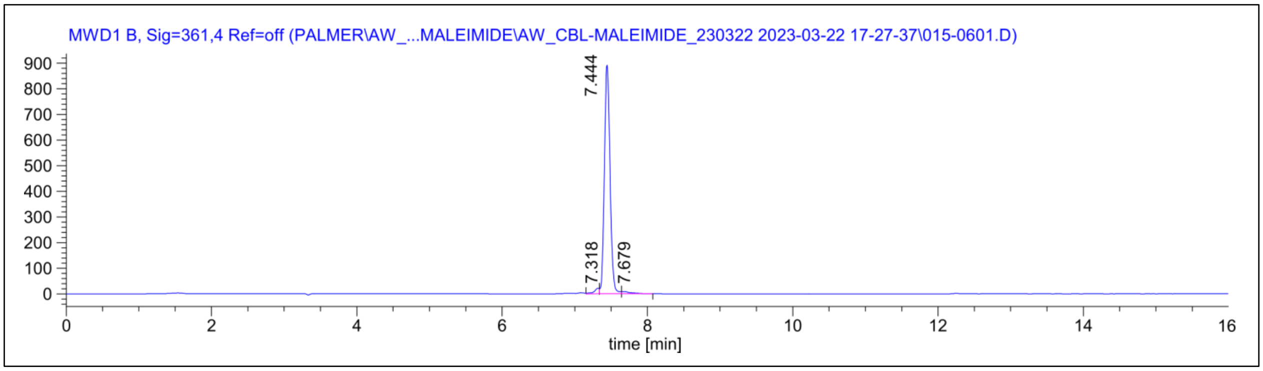

ESI MS spectrum

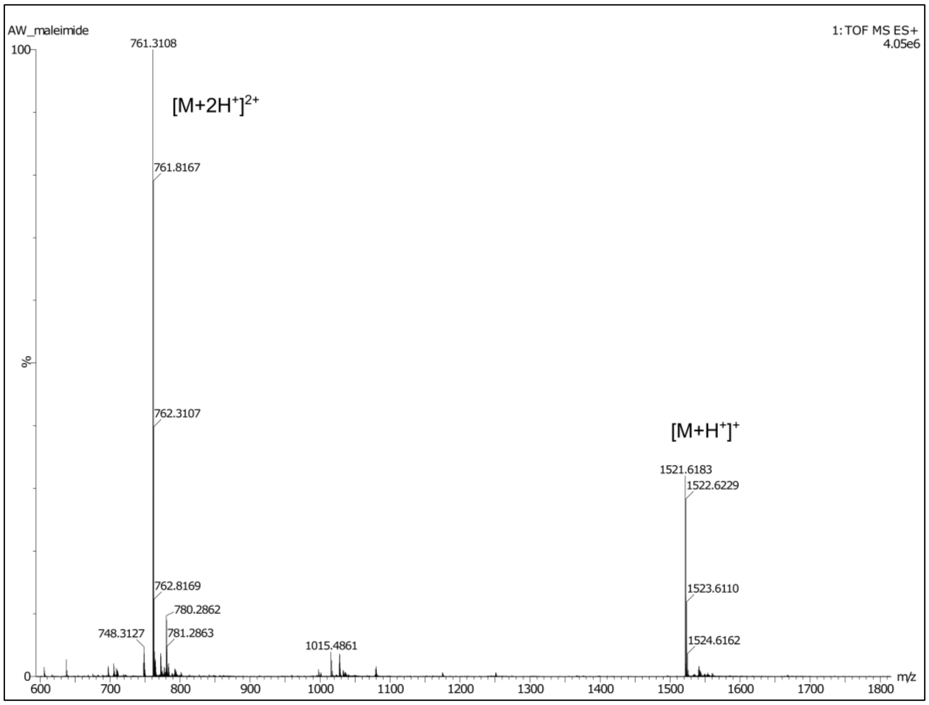

#### 3.6. Synthesis of Cbl-PNA conjugates

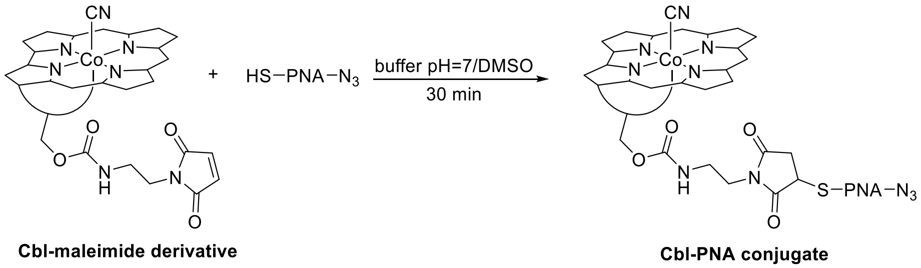

**Scheme S3.** Schematic representation of Cbl-PNA conjugate synthesis.

Cbl-maleimide derivative (3 µmol, 2 equiv.) was dissolved in a potassium phosphate monobasic buffer (pH=7, 200 µL), while the PNA linker (1.5 µmol, 1 equiv.) was separately dissolved in DMSO (200 µL). The Cbl solution in the buffer was then added to the PNA solution in DMSO (*Note: the order is crucial, as PNA linker is insoluble in the buffer and tends to precipitate if added in reverse*), and the mixture was stirred for 30 min. Subsequently, the reaction mixture was diluted with MeOH (up to 2 mL), precipitated with $Et_2O$ (10 mL), and centrifuged. After drying, the remaining solid was dissolved in water and purified by RP column chromatography with a mixture of MeCN and $H_2O$ as eluents (gradually from 10% to 25% v/v) or via semipreparative HPLC (see method below).

HPLC purification method for Cbl-PNA conjugates (λ= 254 nm and 361 nm):

| Time [min] | Water + 0.02%TFA [%] | Acetonitrile [%] |
|---|---|---|
| Initial | 90 | 10 |
| 5 | 90 | 10 |
| 24 | 50 | 50 |
| 25 | 90 | 10 |
| 27 | 90 | 10 |

HPLC analytical method for Cbl-PNA conjugates (λ= 254 and 361 nm):

| Time [min] | Water + 0.02%TFA [%] | Acetonitrile [%] |
|---|---|---|
| Initial | 90 | 10 |
| 10 | 30 | 70 |
| 13 | 30 | 70 |
| 14 | 90 | 10 |
| 16 | 90 | 10 |

#### 3.7. Characterization of Cbl-PNA conjugates

<u>Cbl-PNA</u>

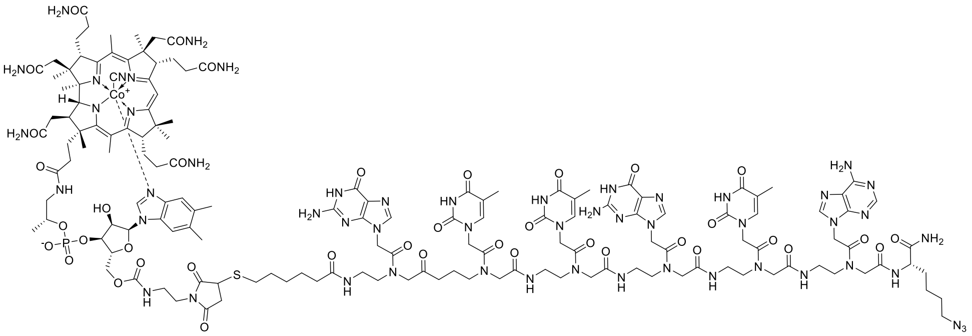

The compound was obtained as a red solid; yield: 30% (*Note: The yield is affected by the use of crude PNA linker as the limiting reagent and can be improved by using purified PNA linker*). HRMS (ESI) m/z [M + 2H]$^{2+}$ calculated for $C_{148}H_{200}CoN_{54}O_{39}PS$, 1739.7053; found, 1739.7042. $t_R$ (RP-HPLC): 7.74 min.

HPLC chromatogram

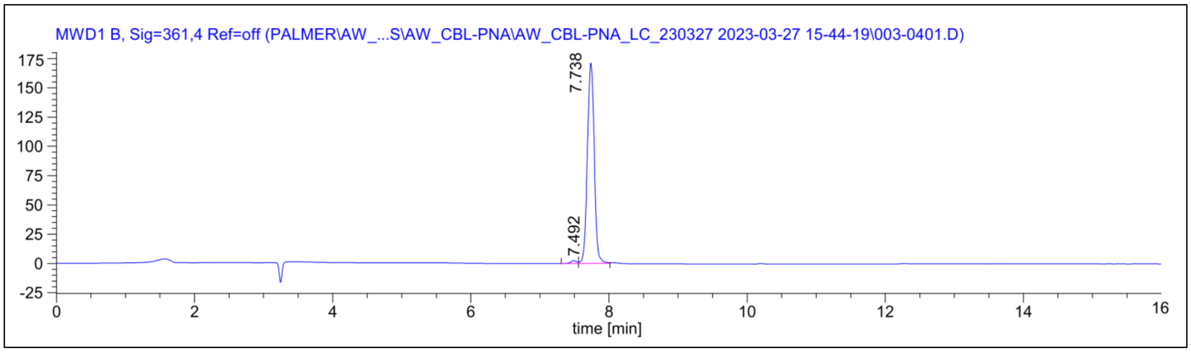

ESI MS spectrum

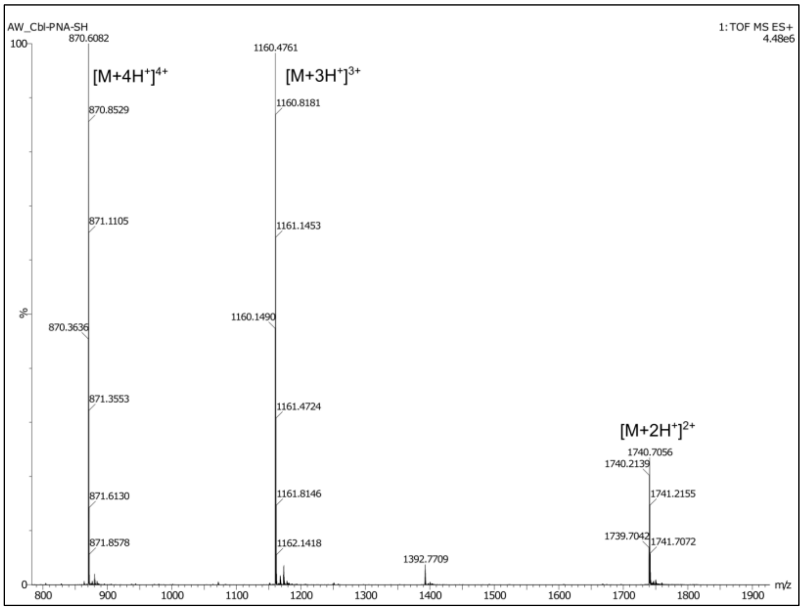

Cbl-PNA$_{scr}$

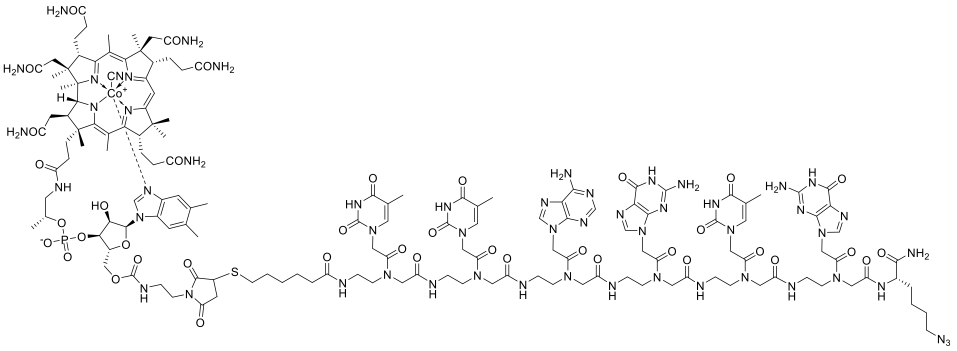

The compound was obtained as a red solid; yield: 42%. (*Note: The yield is affected by the use of crude PNA linker as the limiting reagent and can be improved by using purified PNA$_{scr}$ linker* HRMS (ESI) m/z $[M + 2H]^{2+}$ calculated for $C_{148}H_{200}CoN_{54}O_{39}PS$, 1739.7053; found, 1739.7042. $t_R$ (RP-HPLC): 7.73 min.

HPLC chromatogram

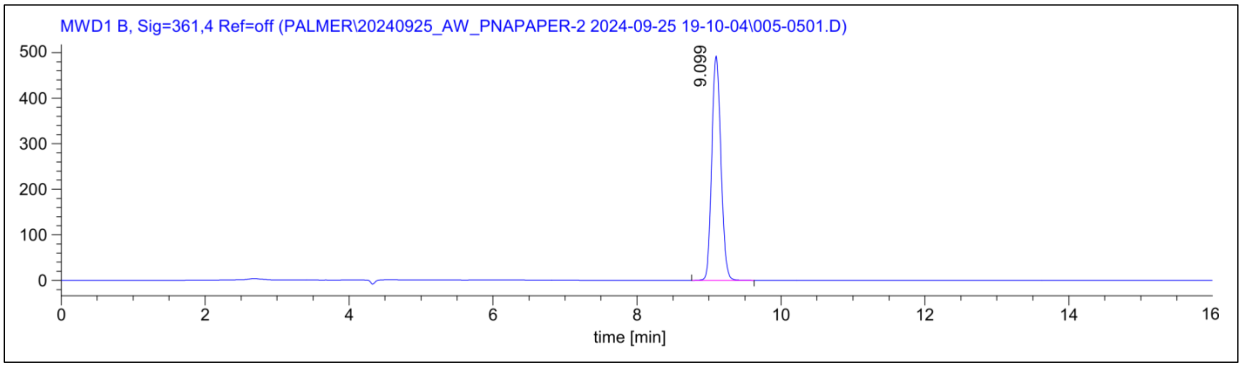

ESI MS spectrum

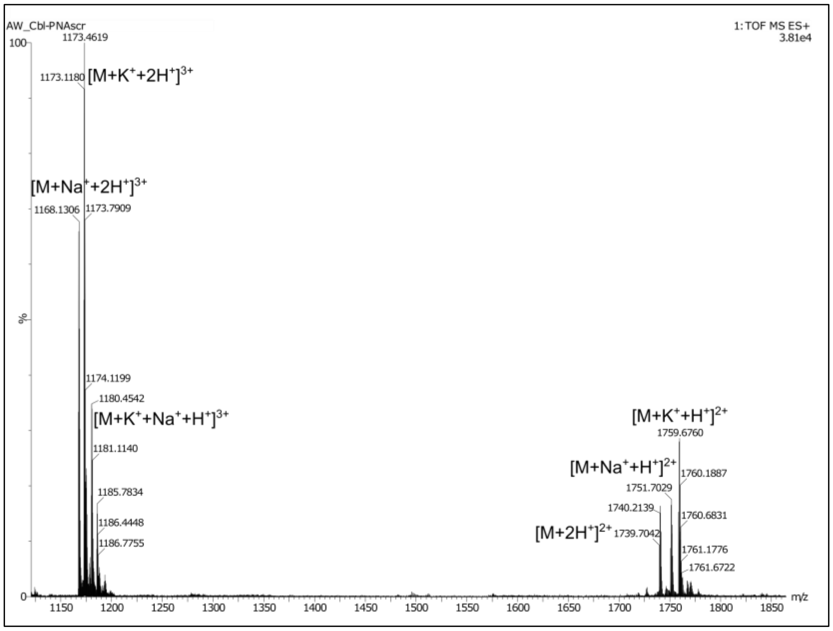

#### 3.8. Synthesis of Cbl-PNA-dye probes

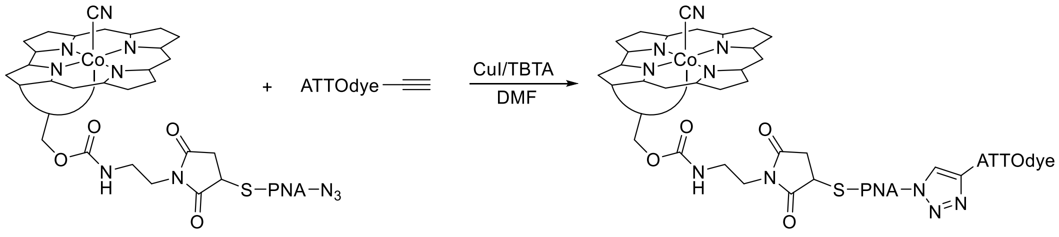

**Scheme S4.** Schematic representation of Cbl-PNA-dye probe synthesis.

Synthesis of Cbl-fluorophore probes was performed as previously described[4] with modifications. Catalyst solution: CuI (1 mg, 5 µmol) and TBTA (5 mg, 10 µmol) were dissolved in DMF (250 µL) and stirred for 20 min. Cbl-PNA conjugate (2 µmol, 2 equiv.) was dissolved in DMSO (20 µL) and subsequently diluted with DMF (200 µL) (*Note: DMSO addition is required as Cbl-PNA conjugates exhibit challenging solubility in DMF*). Dye propargylamide (1 µmol, 1 equiv.) was dissolved in DMF (30 µL) and added to the solution of Cbl-PNA conjugate. Subsequently the catalyst solution was added (see above) and the mixture was stirred at 35°C overnight. When full conversion of the dye was achieved (determined via HPLC, see *Notes* below) the reaction mixture was diluted with MeOH (5 mL), poured into $Et_2O$ (15 mL) and the precipitate was centrifuged and dried. The dried solid was then dissolved in minimal amount of DMSO, and purified via semipreparative HPLC (see method below).

*Notes: All reactions were run until full conversion of the dye was achieved (determined via HPLC). Full conversion of ATTO590 propargylamide was achieved by running the reaction overnight at 35°C. In contrast, the reaction with ATTO488-propargylamine required heating to 40°C and an extended reaction time of 48 h to reach full conversion. Reactions involving Cbl-PNA conjugates generally demand a high catalyst load, which results in the formation of a byproduct with an iodine atom incorporated into the triazole ring, forming alongside the desired product in approximately a 1:1 ratio (confirmed by HPLC and HR MS; data not shown). Further reaction optimization was not pursued due to the high cost of the ATTO fluorophores. Although the byproduct exhibits performance comparable to the desired probe (data not shown), all experiments in this study were conducted using the purified probe, which does not contain iodine.*

| HPLC analytical method for probes with ATTO590 dye (λ= 254, 361, 590 nm): | | | | | |
|---|---|---|---|---|---|
| Purification method | | | Analytical method | | |
| Time [min] | Water + 0.02%TFA [%] | Acetonitrile [%] | Time [min] | Water + 0.02%TFA [%] | Acetonitrile [%] |
| Initial | 90 | 10 | Initial | 90 | 10 |
| 5 | 90 | 10 | 10 | 30 | 70 |
| 23 | 35 | 65 | 13 | 30 | 70 |
| 24 | 20 | 80 | 14 | 20 | 80 |
| 27 | 20 | 80 | 15 | 20 | 80 |
| 28 | 90 | 10 | 16 | 50 | 50 |
| 30 | 90 | 10 | 17 | 90 | 10 |
| | | | 20 | 90 | 10 |

| HPLC analytical method for probes with ATTO488 dye (λ= 254, 361, 488 nm): | | | | | |
|---|---|---|---|---|---|
| Purification method | | | Analytical method | | |
| Time [min] | Water + 0.02%TFA [%] | Acetonitrile [%] | Time [min] | Water + 0.02%TFA [%] | Acetonitrile [%] |
| Initial | 90 | 10 | Initial | 90 | 10 |
| 3 | 90 | 10 | 2 | 80 | 20 |
| 4 | 80 | 20 | 15 | 80 | 20 |
| 18 | 80 | 20 | 16 | 70 | 30 |
| 19 | 50 | 50 | 17 | 70 | 30 |
| 20 | 50 | 50 | 18 | 90 | 10 |
| 21 | 90 | 10 | 20 | 90 | 10 |
| 25 | 90 | 10 | | | |

#### 3.9. Characterization of Cbl-PNA-dye probes

Cbl-PNA-ATTO590

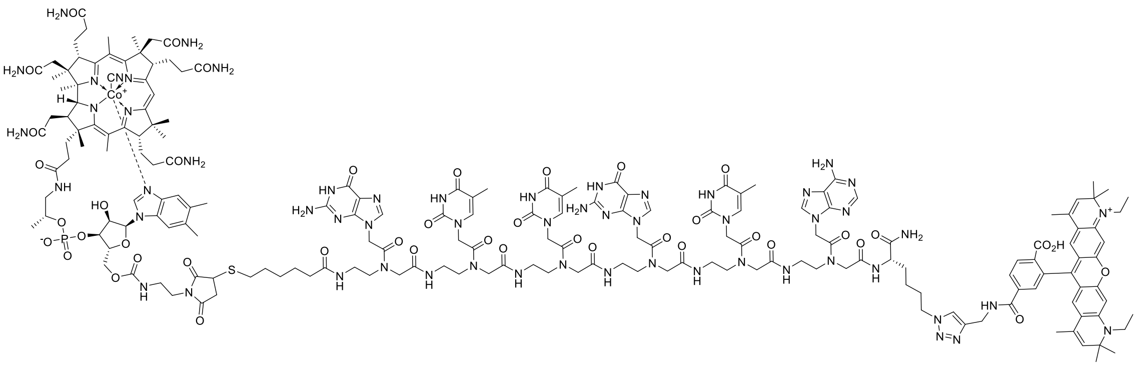

The compound was obtained as a violet solid. HRMS (ESI) m/z $[M^{+} + 3H^{+}]^{4+}$ calculated for $C_{188}H_{243}CoN_{57}O_{43}PS^{+}$, 1027.1837; found, 1027.1840. $t_R$ (RP-HPLC): 9.34 min.

HPLC chromatogram

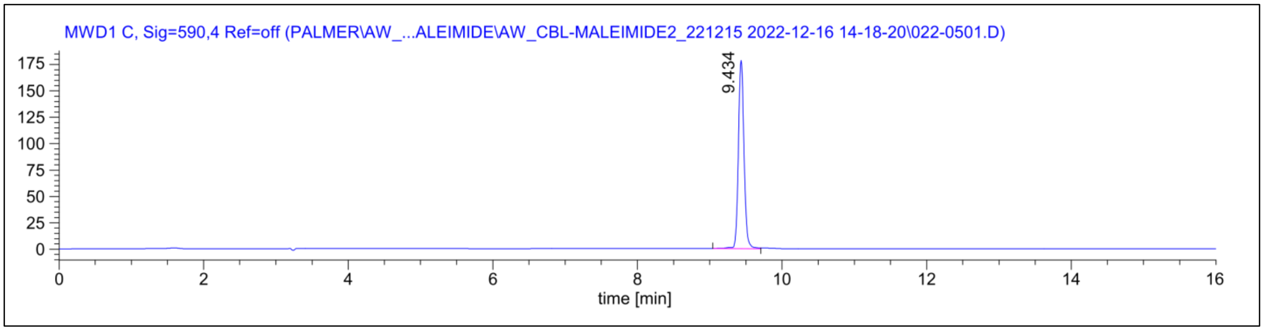

ESI MS spectrum

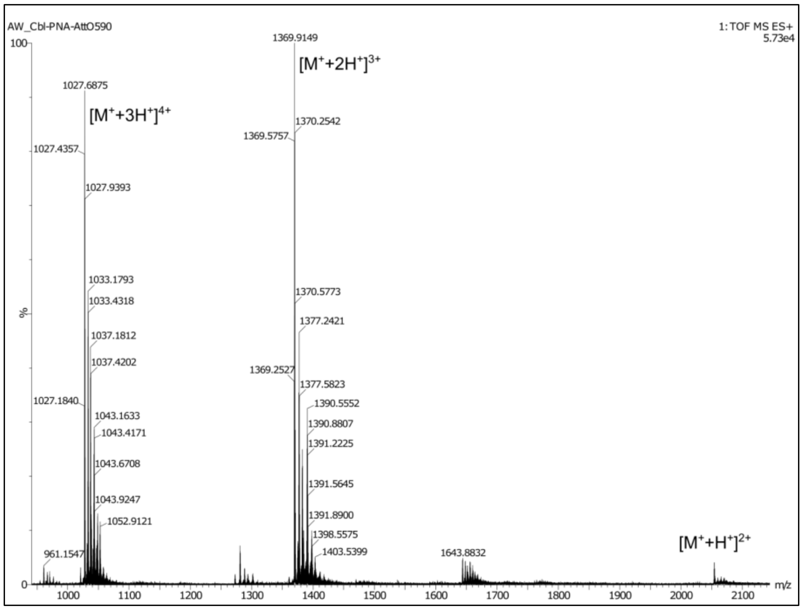

Cbl-$PNA_{scr}$-ATTO590

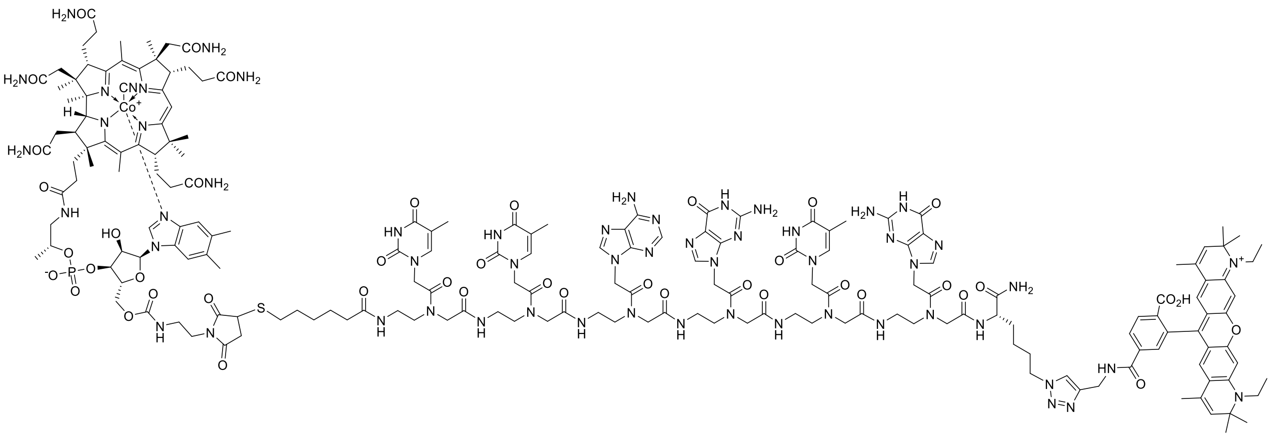

The compound was obtained as a violet solid. HRMS (ESI) m/z $[M^{+} + K^{+} + H^{+}]^{3+}$ calculated for $C_{188}H_{241}CoN_{57}O_{43}PSK^{+}$, 1381.8945; found, 1381.8956. $t_R$ (RP-HPLC): 9.47 min.

HPLC chromatogram

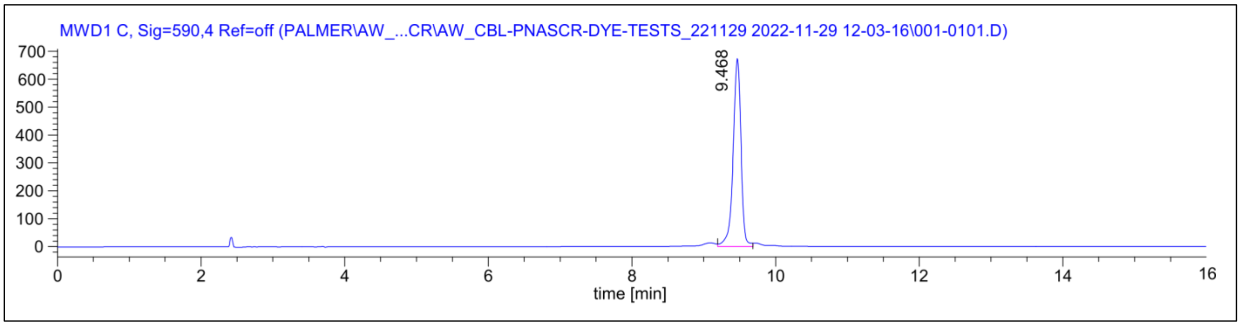

ESI MS spectrum

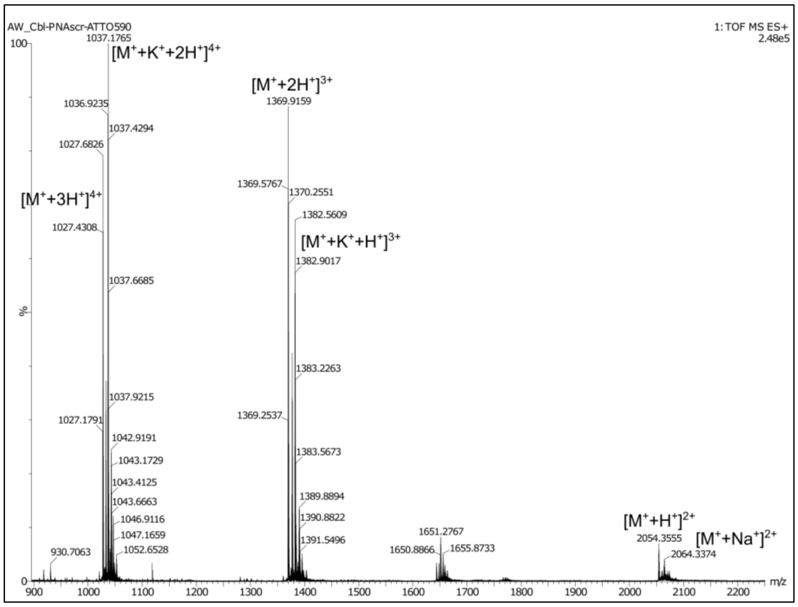

Cbl-PNA-ATTO488

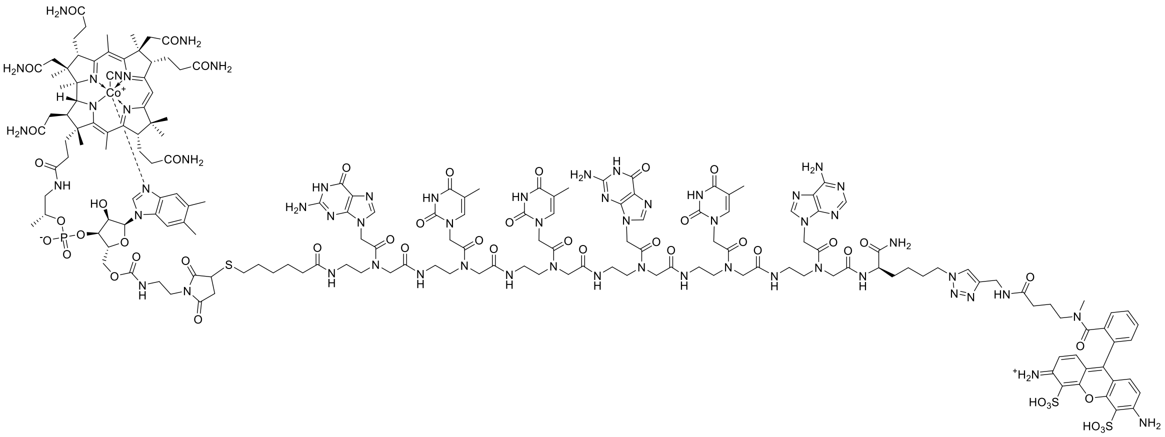

The compound was obtained as a red solid. HRMS (ESI) m/z $[M^+ + 3H^+]^{4+}$ calculated for $C_{176}H_{228}CoN_{58}O_{48}PS_3^+$, 1026.8848; found, 1026.8810. $t_R$ (RP-HPLC): 9.73 min.

HPLC chromatogram

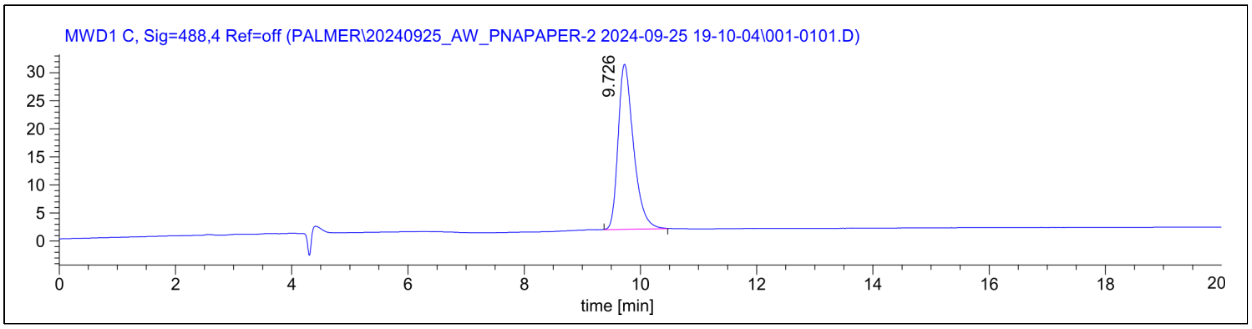

ESI MS spectrum

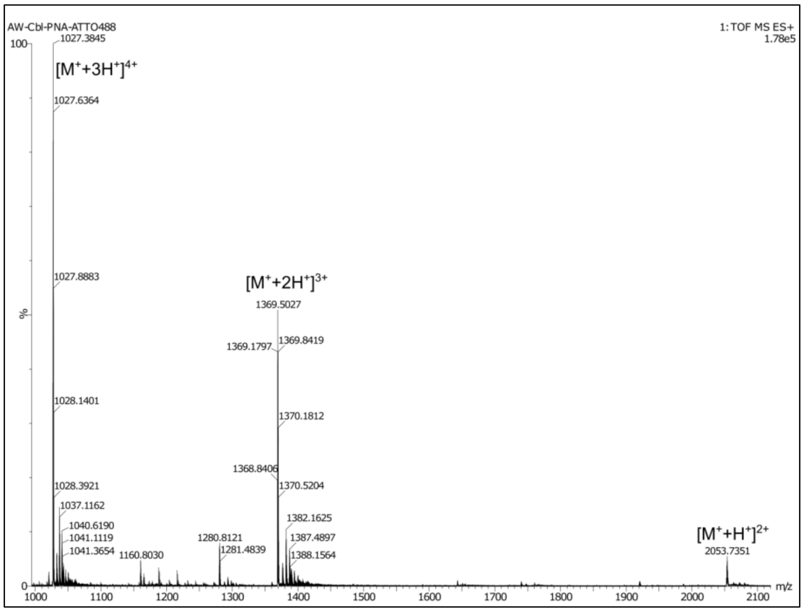

#### 3.10. Synthesis and characterization of PNA-ATTO590 probe

PNA-ATTO590 conjugate

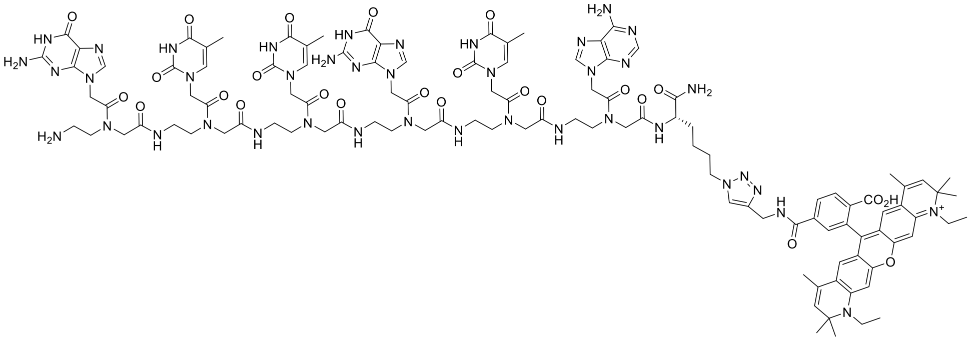

The compound was synthesized according to the protocol 3.8 with the following modifications: 1) 6 equiv. of $PNA_{trunc}$ were used. 2) 125 µL of the catalyst solution was used. 3) The reaction was performed at room temperature overnight. The compound was obtained as a violet solid. HRMS (ESI) m/z $[M^{+}+H^{+}]^{+}$ calculated for $C_{112}H_{137}N_{41}O_{25}^{+}$, 1228.0349; found, 1228.0865. $t_R$ (RP-HPLC): 11.81 min.

HPLC chromatogram

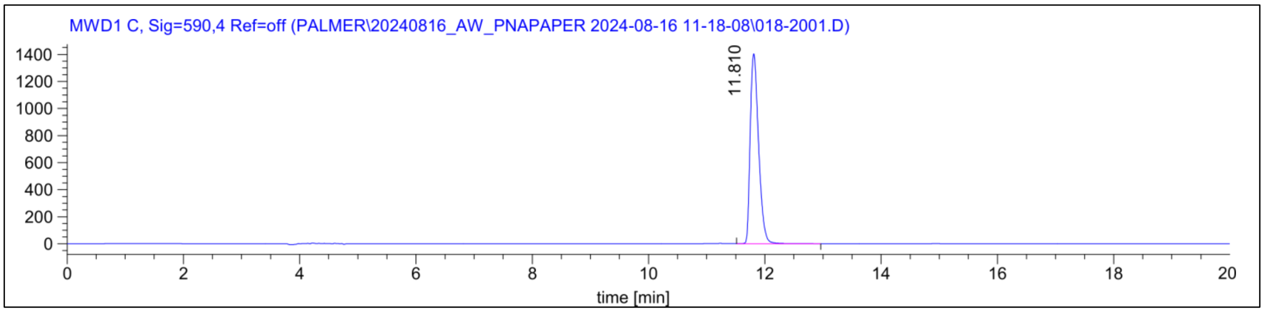

ESI MS spectrum

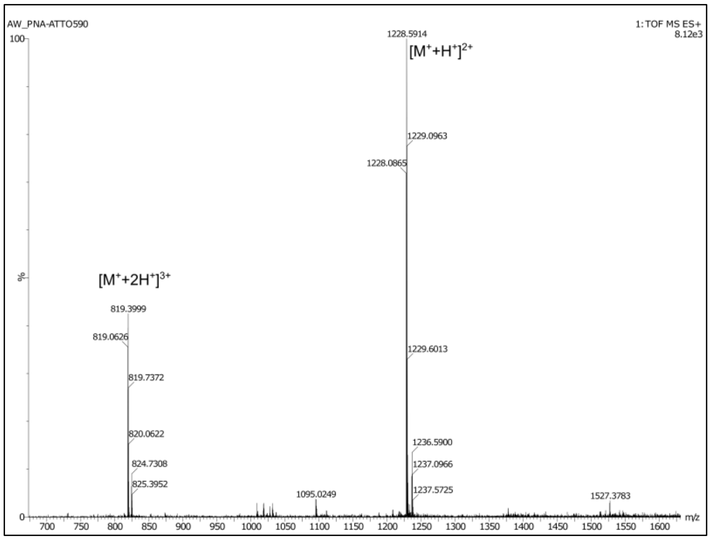

### 4. References

(1) Johnson Jr, J. E.; Reyes, F. E.; Polaski, J. T.; Batey, R. T. B12 Cofactors Directly Stabilize an MRNA Regulatory Switch. *Nature* **2012**, *492* (7427), 133–137.

(2) Gholamalipour, Y.; Karunanayake Mudiyanselage, A.; Martin, C. T. NAR Breakthrough Article 3 End Additions by T7 RNA Polymerase Are RNA Self-Templated, Distributive and Diverse in Character—RNA-Seq Analyses. *Nucleic Acids Res.* **2018**, *46* (18), 9253–9263.

(3) Wierzba, A. J.; Wojciechowska, M.; Trylska, J.; Gryko, D. Vitamin B12 – Peptide Nucleic Acid Conjugates BT - Peptide Conjugation: Methods and Protocols; Hussein, W. M., Stephenson, R. J., Toth, I., Eds.; Springer US: New York, NY, **2021**; pp 65–82.

(4) Braselmann, E.; Wierzba, A. J.; Polaski, J. T.; Chromiński, M.; Holmes, Z. E.; Hung, S. T.; Batan, D.; Wheeler, J. R.; Parker, R.; Jimenez, R.; et al. A Multicolor Riboswitch-Based Platform for Imaging of RNA in Live Mammalian Cells. *Nat. Chem. Biol.* **2018**, *14* (10), 964–971.
